# Symmetries and Continuous Attractors in Disordered Neural Circuits

**DOI:** 10.1101/2025.01.26.634933

**Authors:** David G. Clark, L.F. Abbott, Haim Sompolinsky

## Abstract

A major challenge in neuroscience is reconciling idealized theoretical models with complex, heterogeneous experimental data. We address this challenge through continuous-attractor networks, which model how neural circuits represent continuous variables such as head direction or spatial location through collective dynamics. Classical continuous-attractor models rely on continuous symmetry in the recurrent weights to generate a manifold of stable states, predicting tuning curves that are identical up to shifts. However, mouse head-direction cells exhibit substantial heterogeneity in their responses, seemingly incompatible with this classical picture. We demonstrate that mammalian circuits could nevertheless rely on the same dynamical mechanisms as classical continuous-attractor models. We construct recurrent neural networks directly from experimental head-direction tuning curves that exhibit quasi-continuous-attractor dynamics, then develop a statistical generative process quantitatively capturing the structure of tuning heterogeneity. This enables large-*N* analysis, where we show through dynamical mean-field theory that these networks become equivalent to classical ring-attractor models, with Mexican-hat interactions and continuous symmetry that is spontaneously broken, leading to bump states. In the seemingly disordered weights, the continuous symmetry essential to classical models is reflected through eigenvalue degeneracies, positioning spectral structure as a target for detecting continuous-attractor circuits in connectome data. We extend this framework to two-dimensional symmetries, constructing grid-cell models that similarly reduce to classical toroidal attractors. Our work demonstrates that the dynamical mechanisms of classical continuous-attractor models may operate not only in small brains or idealized systems but also in complex mammalian circuits.

## 1 Introduction

Organisms from flies to humans maintain mental representations of continuous variables. These include navigationally relevant variables, such as self-orientation and location, as well as more abstract cognitive variables. Continuous-attractor models explain how networks of interacting neurons, each individually lacking memory, can maintain these representations through their collective dynamics. In these models, synaptic weights are configured to generate a manifold of stable states that matches the group structure of the encoded variable, providing persistent working memory of this variable. Furthermore, by incorporating velocity inputs that modify network states, such systems can integrate these inputs.

Classical continuous-attractor models generate marginally stable manifolds of fixed points through three ingredients (Fig. 2A, B) [1]. First, neurons must transform input currents into firing rates nonlinearly. Second, the recurrent weights must support Turing pattern formation, typically through a Mexican-hat profile with local excitation and longer-range inhibition. Third, these weights must possess a continuous symmetry corresponding to the encoded variable: one-dimensional circular symmetry for head direction, two-dimensional symmetry for grid-cell responses, and so on. The essential dynamical mechanism is that pattern formation spontaneously breaks the continuous symmetry, and the resulting activity patterns form a continuous manifold with the same symmetry as the weights and thus as the encoded variable. As a consequence of this mechanism, the tuning curves of individual neurons (i.e., their responses as a function of the encoded variable) are identical up to neuron-specific shifts (Fig. 2C).

**Figure 1:**
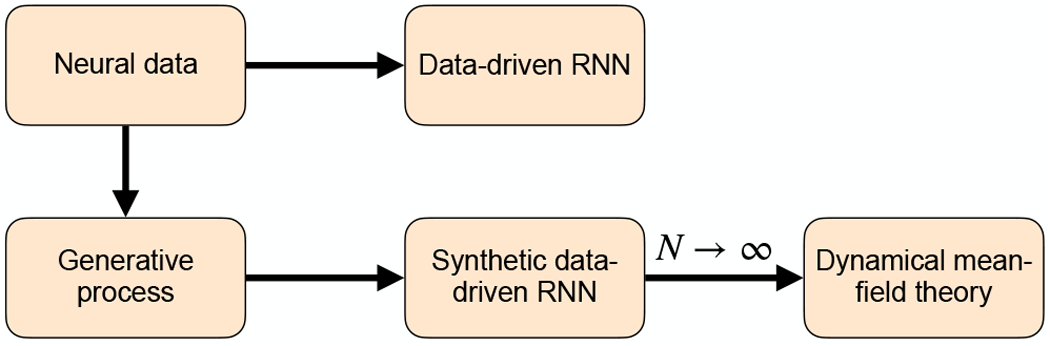
Flow of this paper. We use neural data (top left) to construct a recurrent neural network model directly (rightward arrow) and to build a generative process that captures the data statistics (downward arrow). Such a generative process enables construction of larger network models and dynamical mean-field analysis.

**Figure 2:**
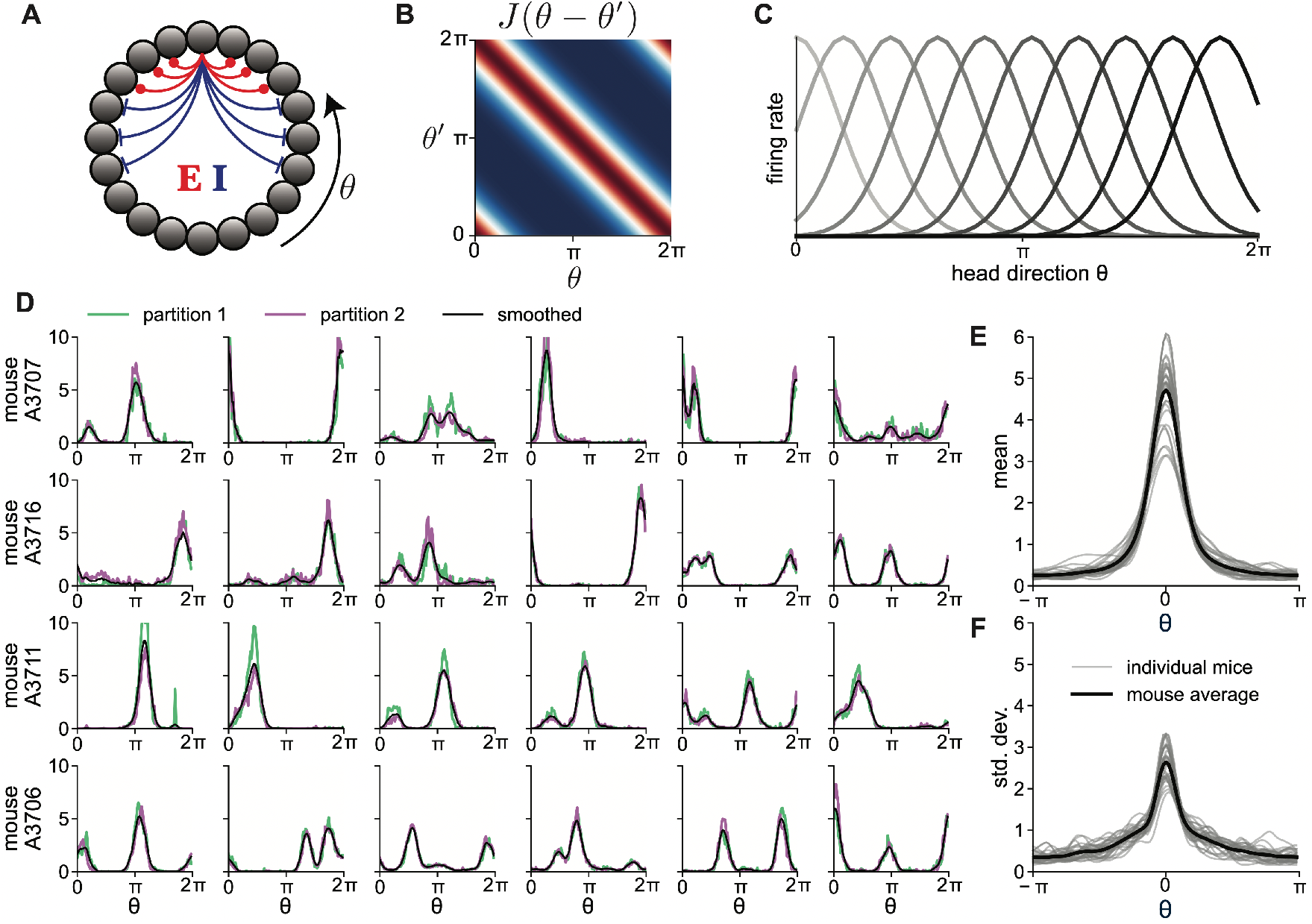
Classical continuous-attractor models fail to capture experimental data. **(A)** Schematic of a classical ring-attractor model with rotation-invariant weights. **(B)** Circulant, Mexican-hat synaptic weight profile characteristic of classical ring-attractor models. **(C)** Translation-invariant tuning curves predicted by classical ring-attractor models. Tuning curves have identical shapes with neuron-specific shifts, tiling the *θ* axis. **(D)** Randomly selected single-neuron tuning curves (arbitrary units) from postsubicular head-direction cells (*N*_data_ = 1533 neurons across 31 mice) [19, 45]. Data shown are from the four mice with the most recorded neurons. Green and purple traces show unsmoothed tuning curves computed from disjoint data partitions (Sec. B2). Black traces show cross-validated smoothed tuning curves. **(E)** Mean of center-of-mass-aligned tuning curves for each mouse (thin lines, 31 mice). Thick line represents the average across all mice. **(F)** Standard deviation of center-of-mass-aligned tuning curves for each mouse (thin lines, 31 mice). Thick line represents the average across all mice.

A canonical example of continuous-attractor models is the ring-attractor model for one-dimensional circular variables, such as head direction or visual-stimulus orientation [2–8]. The experimental discovery of a ring-attractor circuit in the brain of *Drosophila melanogaster* several decades after the formulation of this model provided a notable validation of theoretical approaches. In the central complex, a region called the ellipsoid body has an anatomical ring structure that produces a localized bump of activity that accurately tracks the animal’s orientation, appearing to be a nearly exact instantiation of a classical ring-attractor model [9–11]. Ring-attractor-like circuitry has also been discovered in zebrafish [12].

In this paper, we investigate how mammalian brains might implement stable representations of continuous variables. Rodents, for example, possess neurons that encode head direction, analogous to the heading representation in flies, in regions including the anterodorsal nucleus of the thalamus and the postsubiculum [13–15]. Similarly, recordings from the medial entorhinal cortex have revealed grid cells that represent spatial location through hexagonally arranged firing fields, with each cell’s activity peaking at the vertices of a hexagonal lattice that spans the environment [16, 17]. An open question is whether these are examples of the classical continuous-attractor mechanism described above. Evidence suggests that the intrinsic circuit structure constrains neuronal activity to low-dimensional manifolds, and that this manifold confinement persists even during sleep, a hallmark prediction of continuous-attractor models. Such structured activity patterns have been demonstrated in both rodent head-direction cells, where activity is confined to a manifold with ring topology [18, 19], and grid cells, where activity traces out a manifold with toroidal topology [20–22].

While mammalian neural circuits possess the requisite nonlinear neurons and strong excitatory and inhibitory recurrent connectivity, their tuning to continuous variables is markedly different. Instead of having tuning curves that are identical up to neuron-specific shifts—a direct consequence of continuous symmetry in the weights of classical models—mammalian circuits show diverse response profiles. This tuning heterogeneity is evident in the rodent head-direction system, which we examine in detail; in the medial entorhinal cortex, where neurons display a continuous spectrum of “grid scores” [23–25], with only a small subset exhibiting the periodicity predicted by classical models; and elsewhere in the brain [26].

These observations raise two interconnected questions:

1. What is the nature of heterogeneous tuning in biological neural circuits?
2. To what extent are classical models, based on dynamical breaking of continuous symmetry, relevant to such circuits?

Question (2) has a practical corollary in light of the nascent but growing availability of large-scale electron-microscopy reconstructions of detailed synaptic connectivity, or “connectomes,” in mammalian cortex [27, 28]. In classical continuous-attractor models, the connectivity exhibits easily identifiable structure (e.g., a circulant weight matrix for ring-attractor models), providing clear signatures to search for in connectomics data. In heterogeneous mammalian circuits, it remains unknown which connectivity features, if any, are conserved from classical models and, relatedly, what signatures should be targeted in connectome analyses.

Recent work has begun to separately address aspects of these questions. Regarding question (1), Mainali et al. [29] demonstrated that sampling tuning curves from a Gaussian process and passing them through a nonlinear activation function provides a quantitatively accurate model of topological features of hippocampal place-cell firing fields across dimensionalities and environments. Relatedly, Gaussian process-based latent-variable models of neuronal tuning that allow for unsupervised discovery of topologically nontrivial latent variables have been proposed [30, 31].

Regarding question (2), various constructions of network models with heterogeneous continuous-attractor manifolds have been proposed, though no explicit correspondence to classical continuous-attractor models has been established. Darshan and Rivkind [32] applied an earlier result of Rivkind and Barak [33] to circular manifolds, showing that a combination of high- and low-rank weights generates ring attractors with heterogeneous tuning. Pollock and Jazayeri [34] developed methods for constructing recurrent neural networks with prescribed dynamics, providing another approach for generating ring attractors with heterogeneous tuning. Within the broader context of low-rank recurrent neural networks, Mastrogiuseppe and Ostojic [35] and Beiran et al. [36] showed that random low-rank weights with appropriate correlations can generate ring attractors with amplitude variability in tuning. While these works provide specific constructions of continuous-attractor models with heterogeneous tuning, such constructibility is guaranteed on general grounds by the universal approximation capability of recurrent neural networks (Sec. A5) [37]. These approaches neither establish any relationship to classical models nor identify which connectivity features are conserved between classical and heterogeneous models. Furthermore, they have not been constrained by the statistical structure of tuning heterogeneity observed in experiential data.

This paper jointly addresses questions (1) and (2) within a unified framework, demonstrating that heterogeneous mammalian circuits could rely on the same symmetry-based dynamical mechanism as classical continuous-attractor models. Briefly, we construct recurrent neural networks that generate experimentally observed tuning curves as stable attractor manifolds, develop a statistical generative process capturing biological tuning heterogeneity (question (1)), show through dynamical mean-field theory that these heterogeneous networks are equivalent to classical models at large *N* (question (2)), and identify spectral degeneracies as the key conserved connectivity signature between classical and heterogeneous models.

### 1.1 Outline of the paper

The flow of this paper is schematized in Fig. 1. We focus on the rodent head-direction system. By examining recordings from head-direction cells in the postsubiculum of freely moving mice [19], we first demonstrate that the heterogeneity in neuronal tuning curves rules out the possibility that the recorded neurons themselves comprise a classical ring-attractor circuit. The observed tuning could arise from feedforward input from a classical ring-attractor circuit elsewhere in the brain [38], distributed attractor dynamics across multiple coupled regions, or recurrent connections among the heterogeneous cells themselves. Regardless of which mechanism operates in this particular system, understanding how recurrent connectivity alone can generate continuous attractor manifolds with heterogeneous tuning is of broad interest, given the prevalence of such tuning across mammalian brain regions [26]. Using an optimization procedure (building on neural activity-constrained modeling approaches [39–41]; see also nascent connectome-constrained modeling approaches [42–44]), we construct recurrent neural networks directly from the experimentally observed tuning curves, demonstrating that appropriate recurrent connections among the heterogeneous cells can maintain this representation and exhibit quasi-continuous-attractor dynamics.

To extend this optimization procedure beyond the limited population sizes available in experimental recordings, we developed a statistical generative process for sampling synthetic tuning curves that addresses question (1). This generative process is based on a Gaussian process with circular symmetry, motivated by statistical analysis of the experimental tuning-curve distribution. The resulting model reproduces the statistical structure of biological head-direction cells with quantitative accuracy across multiple measures, enabling analysis of network properties at scales reflecting mammalian neural circuits.

Using the generative process, we analyze the structural properties of optimal weight matrices at large *N*. Despite appearing disordered, these matrices reflect an underlying circular symmetry that manifests as doubly degenerate eigenvalues. This symmetry, however, is realized differently than in classical ring-attractor models [2] and in other models [32]. While classical models embed circular symmetry through orderly Fourier eigenvectors across neurons, our model achieves the same symmetry through disordered, random vectors. This distinction has direct implications for connectomics: eigenvalue degeneracies, rather than eigenvector structure, may provide a more reliable signature for identifying continuous-attractor circuits in mammalian brains. We formalize the relationships between continuous symmetries, eigenvalue degeneracies, and symmetries of the dynamics in Sec. A1.

Finally, we develop a dynamical mean-field theory to analyze these networks at large *N*. This reveals a central result, answering question (2): the network dynamics become equivalent to a classical ring-attractor model. The mean-field description recovers all essential features of classical continuous-attractor models—a nonlinear activation function, Mexican-hat interactions, and continuous symmetry—with spontaneous symmetry breaking generating stable bump states through a Turing instability. The effective activation function has interesting differences from a conventional activation function that we discuss. Angular-velocity inputs are faithfully integrated within these mean-field dynamics. We extend this framework to higher dimensions by modeling grid cells in the medial entorhinal cortex, which are thought to comprise a toroidal attractor manifold.

## 2 Examining a dataset of head-direction cells

We analyzed recordings of head-direction cells from the postsubiculum of freely moving mice [19] (data publicly available at [45]). Both the postsubiculum and the anterodorsal thalamic nucleus [18] contain neurons that encode head direction in a manner suggestive of ring-attractor dynamics, with activity patterns confined to a stable manifold with ring topology across different environments and during sleep [18, 19]. Previous analysis by Duszkiewicz et al. [19] emphasized the relative stereotypy of head-direction cells by contrasting them with more heterogeneous fast-spiking neurons. Nevertheless, head-direction cells themselves exhibit substantial heterogeneity in their responses, deviating markedly from the predictions of classical continuous-attractor models, as we now show.

We analyzed *N*_data_ = 1533 excitatory neurons, pooled across 31 mice, that were identified as head-direction cells by Duszkiewicz et al. [19]. These cells were classified based on encoding significant head-direction information relative to a time-reversed control distribution, without imposing constraints on tuning-curve shapes such as unimodality. We constructed tuning curves by dividing head-direction angles into 100 bins and computing occupancy-normalized spike counts for each neuron. This binning introduces high-frequency artifacts (Fig. 2D, “partition 1” and “partition 2”). To reduce noise while preserving structure, we applied cross-validated smoothing using a Gaussian kernel. For each neuron, we split the recording into two temporal partitions, computed separate tuning curves, and selected the kernel width that maximized the cross-partition Poisson log-likelihood. Optimal smoothing widths were correlated across partitions (*r* = 0.16, *p <* 0.001). The final tuning curve was obtained by averaging the two smoothed partition curves (Fig. 2D, “smoothed”). All tuning curves were normalized to unit mean across head direction *θ* to enable comparison across neurons and animals. These smoothed, normalized tuning curves form the basis for subsequent analyses.

### 2.1 Failure of classical ring-attractor models

Classical ring-attractor models (Fig. 2A, B) predict tuning curves that are identical up to neuron-specific shifts (Fig. 2C), a direct consequence of circular symmetry in the weights. However, tuning curves of mouse head-direction cells exhibit significant heterogeneity, with neurons displaying variable numbers of irregularly spaced peaks (Fig. 2D). To quantify this heterogeneity, we aligned tuning curves within each of the 31 mice by their circular centers of mass and computed the mean profile (Fig. 2E) and fluctuations (standard deviation) around this within-mouse mean profile (Fig. 2F). While the mean profiles were unimodal and symmetric, the fluctuations were comparable in magnitude to the means, starkly violating the classical prediction of zero fluctuations. Both the mean and fluctuation profiles exhibited remarkable consistency across mice: the gray curves (individual mice) closely track the black curve (cross-mouse average) in both cases. This consistency suggests a universal statistical structure (Sec. 3).

### 2.2 Network construction

The observed heterogeneity rules out the possibility that the recorded neurons themselves comprise a classical ring-attractor circuit. To test whether intrinsic recurrent connectivity alone can generate the observed responses as a continuous attractor manifold, we constructed network models directly from the experimental tuning curves using an optimization principle. This setup is summarized in Sec. B3.1 (Setting 1).

The model we constructed consists of a firing rate-based recurrent neural network of *N* neurons. While we use a rate-based formulation for analytical tractability, key behaviors of these networks generalize to their spiking counterparts [46]. The state of each neuron *i*, where *i* = 1, …, *N*, is described by an input current *x*_*i*_(*t*) and firing rate *ϕ*_*i*_(*t*). The network dynamics are governed by

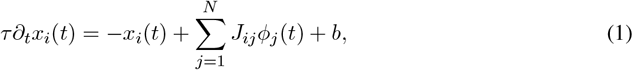

where *J*_*ij*_ is the synaptic weight from neuron *j* to *i, b* is a bias current, and *τ* = 50 ms is the network time constant. The term −*x*_*i*_(*t*) represents passive decay of neuronal activity, resulting in exponential relaxation to rest, with timescale *τ*, in the absence of input. Because *τ* is on a sub-second timescale, stable dynamics on timescales of seconds, minutes, or hours require collective network dynamics. The firing rate of a neuron is related to its input current *x* through a nonlinear function, *ϕ*(*x*). We used a smooth and invertible activation function to enable the conversion between experimentally measured firing rates and the corresponding input currents. Specifically, we used a “softplus” function that interpolates between zero firing for strongly negative inputs and linear firing with unit gain for strongly positive inputs (Sec. B3.1, Eq. B2).

The dataset of normalized tuning curves, pooled across mice, defines a target manifold of activity patterns, 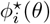, where *i* = 1, …, *N*_data_. This manifold is embedded in *N*_data_-dimensional neuronal space and parameterized by the coordinate *θ*, with the periodicity of *θ* endowing it with the topology of a ring. To incorporate the target manifold into a network of *N* = *N*_data_ neurons, we first obtained the corresponding input-current manifold 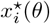 by applying the inverse of the activation function to the observed tuning curves 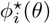. While these target patterns 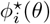 and 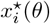 are *a priori* unrelated to the time-dependent network activities *ϕ*_*i*_(*t*) and *x*_*i*_(*t*), our goal is to configure the synaptic weights *J*_*ij*_ and bias *b* such that network dynamics become confined to the target manifold. We formulated this as an optimization problem: at each point 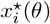 along the manifold, the time derivative should vanish: ∂_*t*_*x*_*i*_(*t*) = 0.

To enable a simple method for preventing runaway activity, which can occur under the non-saturating activation function, we performed an additional normalization of the target manifold along the neuron dimension such that the neuron-averaged firing rate is exactly unity for all *θ* (Sec. B4.1, Fig. B1). As a result, on the target manifold, the right-hand side of Eq. 1 is invariant to a shift in the weights, *J*_*ij*_ → *J*_*ij*_ *− c/N*, if this is accompanied by a shift in the bias, *b* → *b* + *c*. Thus, to prevent runaway activity, we first optimized the weights *J*_*ij*_ with *b* = 0 and then selected a sufficiently large *c >* 0 whose effect is to provide network-wide inhibition that stabilizes the mean activity mode (Sec. B4.2).

We optimized the weights for the vanishing-flow condition using least-squares optimization, in particular, by minimizing the objective function

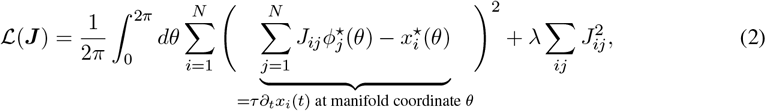

subject to the constraint *J*_*ii*_ = 0 for all *i*. The first term in Eq. 2 ensures that when network activity lies on the target manifold, its time derivative is small (Eq. 1). It involves a sum over neurons and an integral over the manifold. The second term is a regularizer that prevents the weights from growing large. The solution to this optimization problem, explained in Sec. A2, can be expressed for *i*≠ *j* as

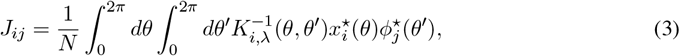

where the kernel *K*_*i,λ*_(*θ, θ*^*′*^) is given by 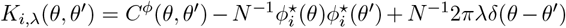. Here, *C*^*ϕ*^(*θ, θ*^*′*^) is the two-point correlation function of the target firing rates,

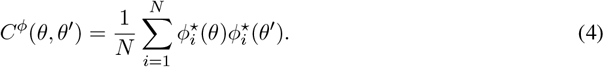

The second term in the kernel accounts for the constraint *J*_*ii*_ = 0, which necessitates that the solution for *J*_*ij*_ does not include neuron *i* as a regressor (this term has a vanishingly small effect for large *N*) and the third is the regularizer. Eq. 3 has a structure reminiscent of Hebbian learning, with 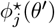 as a presynaptic factor and 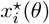 as a postsynaptic factor. Note that the activation function appears only in the presynaptic term. Consequently, the weights do not form a transpose-symmetric matrix, *J*_*ij*_ ≠ *J*_*ji*_, and the resultant network behavior does not lend itself to the analysis of an energy function [47, 48].

At first glance, this optimization problem appears overconstrained: for each neuron *i*, setting 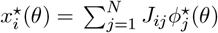 while continuously varying *θ* generates infinitely many constraints, but there are only *N* free parameters per row, namely, *J*_*ij*_ for *j* = 1, …, *N*. However, the problem is actually underconstrained due to the smoothness of the tuning curves, which leads to an approximate frequency cutoff at some *D* ≪ *N* in the space of Fourier coefficients, reducing the effective number of constraints to 2*D* + 1 (Sec. 4). Since the problem is underconstrained, infinitely many weight matrices satisfy the constraints, achieving ∂_*t*_*x*_*i*_(*t*) = 0 at all points *θ* on the target manifold. As *λ* → 0, the unique minimum-norm (pseudoinverse) solution is selected from this set of possibilities.

To enable angular-velocity integration, a capability of biological head-direction systems, we incorporated angular-velocity inputs into the model. Classical ring-attractor models generally achieve this by adding an analogous derivative term to the weights, related to the angular derivative of the rotation-invariant weight profile [4]. We generalized this approach to the data-derived network model by adding an additional term to the weights, yielding

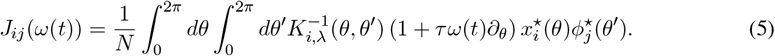

Here, *ω*(*t*) represents angular velocity. This expression can be derived by solving a least-squares optimization problem similar to Eq. 2, but with the objective of setting 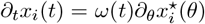, where 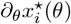 is the tangent vector to the manifold at the coordinate *θ*, instead of ∂_*t*_*x*_*i*_(*t*) = 0. While this approach nominally requires the recurrent weight matrix to be modulated by angular velocity, various biologically plausible mechanisms exist to mimic this effect [4, 9–11, 49, 50].

### 2.3 Dynamics of the data-derived network model

We now examine the dynamics of the data-derived network model, focusing on its ability to generate a quasi-continuous ring attractor manifold. We used regularizer *λ* = 10^*−*6^ and uniform inhibition with strength *c* = 1 to stabilize the mean activity mode (Sec. B4.4). For visualization, we show the *c* = 0 weights in Fig. 3. The effects of *c* and *λ* on the dynamics are shown in Fig. B2.

**Figure 3:**
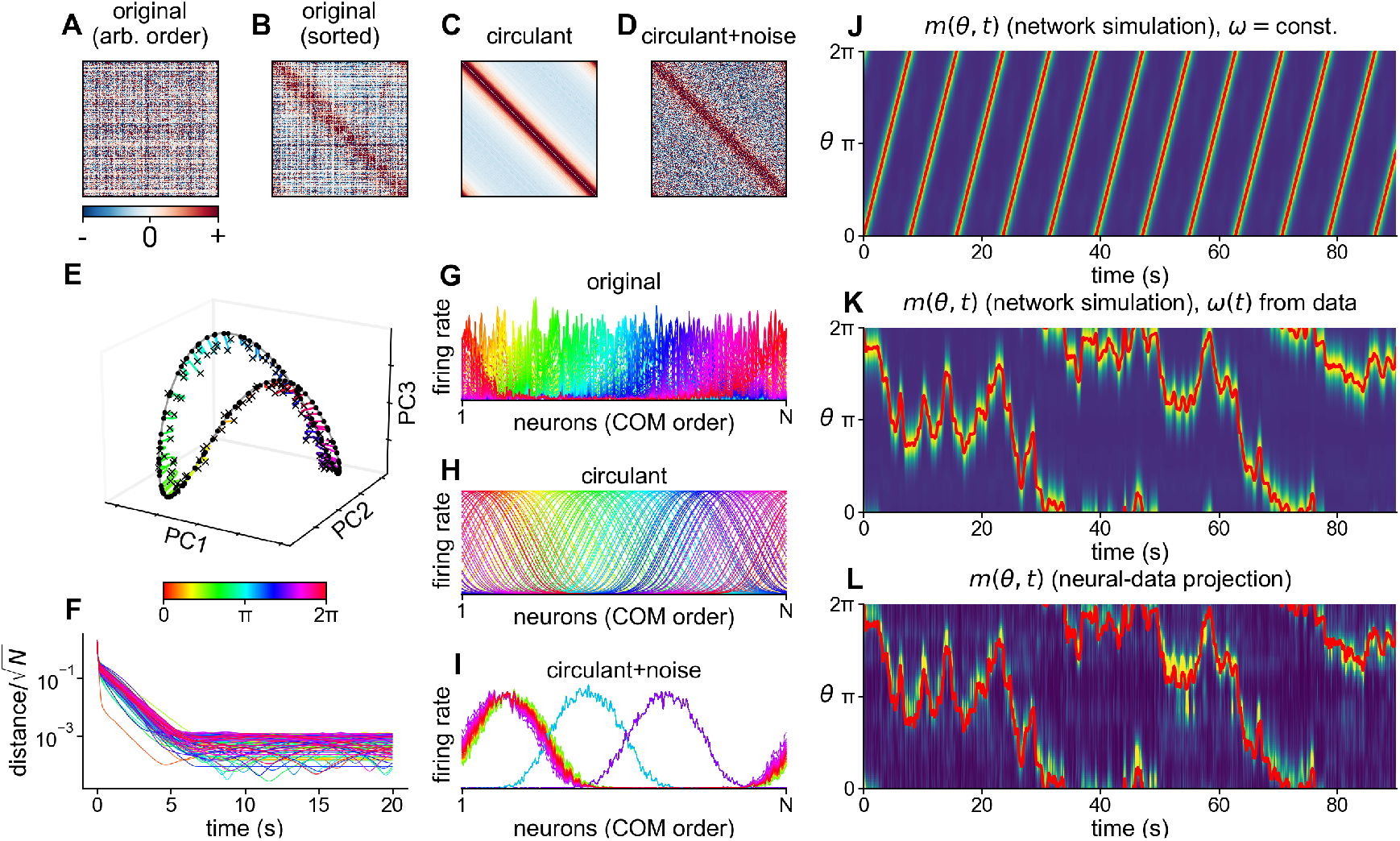
Quasi-continuous-attractor dynamics of the data-derived network model (*N* = *N*_data_ = 1533, uniform inhibition *c* = 1). **(A)** Optimal data-derived weights *J*_*ij*_ with neurons in random order, showing no apparent structure. **(B)** Same weight matrix as (A), with neurons ordered according to the circular centers of mass of their tuning curves. **(C)** Circulant version of matrix (B), obtained by averaging weights along diagonals corresponding to fixed angular differences between neurons. **(D)** Matrix from (C) with added noise, constructed by randomly permuting the residuals between matrices (B) and (C), demonstrating that random noise atop circulant weights destroys quasi-continuous-attractor behavior. **(E)** Network dynamics visualized in the space of the first three principal components of the target manifold. Black crosses mark initial conditions, colored lines show trajectories over 30 s (colors indicate angular locations of initial conditions), and black dots indicate final states. In this and subsequent plots (G, H, I, J), colors indicate the angular locations of initial conditions. **(F)** Distance between each trajectory and its nearest point on the target manifold versus time, demonstrating convergence to the manifold. **(G)** Steady-state activity patterns across neurons (ordered as in B, using data-derived weights) for different initial conditions, showing convergence to stable bump configurations. Panels G, H, and I show activity across neurons at fixed *θ*, not tuning curves across *θ*. **(H)** Steady-state activity patterns using the circulant matrix from (C), showing bump configurations identical up to translation, characteristic of classical ring-attractor models. **(I)** Steady-state activity patterns using the noise-corrupted matrix from (D), showing collapse to a small number of stable states, demonstrating that the structure of matrix (B) is essential for quasi-continuous-attractor dynamics. **(J)** Overlap order parameter *m*(*θ, t*) (Eq. 6) for constant angular-velocity input (*ω* = 0.8 rad/s), demonstrating integration. **(K)** Overlap order parameter *m*(*θ, t*) for actual mouse head-direction trajectories, demonstrating integration of a more complex signal. **(L)** Overlap order parameter *m*(*θ, t*) computed directly from mouse head-direction cell recordings (binned spike counts from time series data, 100 ms bins, mouse A3707).

As described in Sec. 1, classical ring-attractor models rely on circular symmetry in the weights to generate a continuous manifold of fixed points (Fig. 2A). The optimized weights *J*_*ij*_ do not exhibit this symmetry in an obvious way. When neurons are arranged in arbitrary order, the weight matrix *J*_*ij*_ appears, necessarily, unstructured (Fig. 3A). However, when neurons are arranged according to their circular centers of mass, an approximate Mexican-hat structure is revealed (Fig. 3B). One can sort neurons in a similar manner using the singular vectors of the weights (Sec. A3.3) [51].

In this work, we are interested in quasi-continuous-attractor dynamics defined through a separation-of-timescales criterion. Specifically, we seek an invariant manifold with fast, attractive normal flow and slow tangential flow (this dynamical definition is more robust than counting discrete fixed points, which can be unreliable indicators of attractor quality) [52]. Due to the overparameterized nature of the optimization problem, the selected weights produced vanishingly small values of *τ*∂_*t*_*x*_*i*_(*t*) everywhere on the target manifold (Fig. B10). However, achieving near-zero flow on the manifold alone does not guarantee quasi-continuous-attractor dynamics, which requires that network activity both remains confined to this manifold during evolution and returns to it when perturbed away. The uniform inhibition stabilizes the (*N* − 1)-dimensional hyperplane on which the neuron-averaged firing rate is unity, but does not guarantee confinement to the one-dimensional target manifold embedded within this hyperplane.

Despite not being explicitly encouraged within the optimization problem, the optimal weights empirically produce quasi-continuous-attractor dynamics. Initializing the system at regularly spaced points along the target manifold perturbed by noise, we observe rapid convergence back to the manifold within ∼1 s (Fig. 3F), followed by persistence over 30 s. This order-of-magnitude separation between convergence and drift timescales (Fig. 3E) exemplifies quasi-continuous attractor behavior. This stability results from the minimum-norm prescription for the weights (i.e., it is implicitly encouraged). Small weights ensure that most eigenvalues of the Jacobian evaluated on the target manifold, 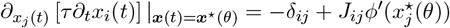, are substantially negative, corresponding to rapid relaxation back to the manifold. Non-minimum-norm constructions by definition add structure to the weight subspace unconstrained by tuning curves, potentially hindering stability. For example, the construction of Darshan and Rivkind [32] generates tuning heterogeneity by operating near chaotic instability (Sec. 5.4, Fig. B12). The stabilizing effect of minimum-norm solutions requires overparameterization, and this relationship between overparameterization and dynamical stability has been noted in previous work [53].

One hypothesis, that we now rule out, is that the optimal weights comprise a circulant matrix with weak noise that does not substantially affect the dynamics. We constructed two alternative weight matrices from the sorted optimal weights (Sec. B4.3). We examined their activity patterns, with neurons sorted by their center-of-mass angles, following after 30 s of evolution from regularly spaced manifold points perturbed by noise.

As we have shown, the optimal weights (Fig. 3B) maintain a continuum of stable, heterogeneous states (Fig. 3G). We first constructed a circulant matrix (Fig. 3C) by averaging the sorted optimal weights along diagonal bands. This produces translation-invariant bump configurations characteristic of classical ring-attractor models (Fig. 3H). We then constructed a circulant-plus-noise matrix (Fig. 3D) by adding back the scrambled residuals between the sorted data-derived and circulant matrices. This causes the network to collapse onto a small number of discrete stable states (Fig. 3I). This degradation in performance implies that the optimal weights are not merely a noisy circulant matrix, but possess structure within their disorder that is essential for maintaining quasi-continuous-attractor dynamics.

To characterize the alignment between network firing rates *ϕ*_*i*_(*t*) and the target manifold 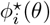, we define an overlap order parameter, given by

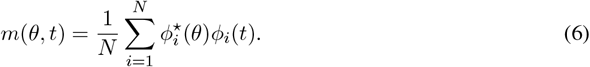

When network activity lies exactly on the manifold at coordinate *ψ*, we have 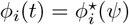, resulting in *m*(*θ, t*) = *C*^*ϕ*^(*θ, ψ*) (Eq. 4). As a function of *θ*, this produces a localized bump of activity peaked at *ψ*. This order parameter is evocative of classical ring-attractor models, which are also described by a time-dependent order parameter that exhibits steady-state localized bumps. Similarly, when the network state moves along the target manifold in response to angular-velocity input, *m*(*θ, t*) takes the form of a traveling bump of activity centered around the integrated angular displacement.

Using *m*(*θ, t*) as a readout of network activity, we tested the network’s ability to integrate angular-velocity inputs. For a constant input *ω* = 0.8 rad/s, visualization of *m*(*θ, t*) shows accurate tracking (Fig. 3J). The network also successfully tracks more complicated trajectories derived from the mouse head-direction data (Fig. 3K), demonstrating robust integration of realistic velocity inputs.

The overlap order parameter *m*(*θ, t*) can also be computed directly from neural recordings by projecting measured firing rates onto measured tuning curves. Using the definition in Eq. 6, we again take 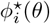 to be the measured tuning curve for neuron *i*, now restricted to a single mouse; and we now take 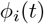 to be the binned spike counts of neuronal time-series data within this mouse. The resulting bump pattern accurately tracks the animal’s head direction (Fig. 3L) and closely resembles the patterns generated by the network model (Fig. 3K). This correspondence provides strong evidence that neuronal activity remains confined to a manifold closely aligned with that defined by the tuning curves [18]. Furthermore, the peak of *m*(*θ, t*) serves as an effective decoder of head direction; in fact, it provides the maximum *a posteriori* estimate of head direction under the assumption of Gaussian noise around tuning curves and a uniform prior over *θ* (Sec. B4.5).

## 3 Statistical generative process

Single-mouse recording sessions contain at most 𝒪(10^2^) neurons, and pooling across all 31 mice yields only *N*_data_ = 𝒪(10^3^) neurons, a small fraction of the head-direction cell population in the postsubiculum. A generative process for tuning curves would allow construction and analysis of network dynamics at scales characteristic of mammalian neural circuits by applying the optimization approach of the previous section to synthetic tuning curves sampled from this process. Characterizing such a process would answer question (1).

We propose a general structure in which tuning curves are generated by passing samples from a rotation-invariant Gaussian process on a ring through a nonlinear activation function. In this section, we fit this model to experimental data using a soft-rectification activation function, achieving quantitatively accurate fits. In subsequent sections, we use the same fundamental structure with an analytically tractable and dynamically convenient saturating activation function to obtain a theoretical understanding of network connectivity and dynamics.

### 3.1 Evidence for distributional circular symmetry

Even when individual tuning curves are heterogeneous, they may arise from a generative process with underlying circular symmetry. Such a symmetry would have two implications. First, it would enable a simplification or “inductive bias” in the generative process itself. Second, it would suggest a mechanism through which circular symmetry could emerge at the population level, despite the disorder present in single-neuron responses.

To test this possibility, we first examined the distribution of center-of-mass angles of tuning curves within each mouse (Fig. 4A). Relative phase relationships cannot be meaningfully compared between neurons recorded in different animals, necessitating single-mouse analysis. Visual inspection suggests that these distributions could be uniform (Fig. 4B), which we confirmed using the Kuiper test for circular uniformity. Across 31 mice, only 2 sessions showed significant (*p <* 0.05) deviations from uniformity before multiple-comparison correction (uncorrected *p*-values ranging from 0.044 to 0.97). After applying the Benjamini-Hochberg correction for multiple comparisons, no sessions showed significant deviations from uniformity (corrected *p*-values ranging from 0.59 to 0.97). This uniform distribution of center-of-mass angles is consistent with circular symmetry in the distribution of tuning curves.

**Figure 4:**
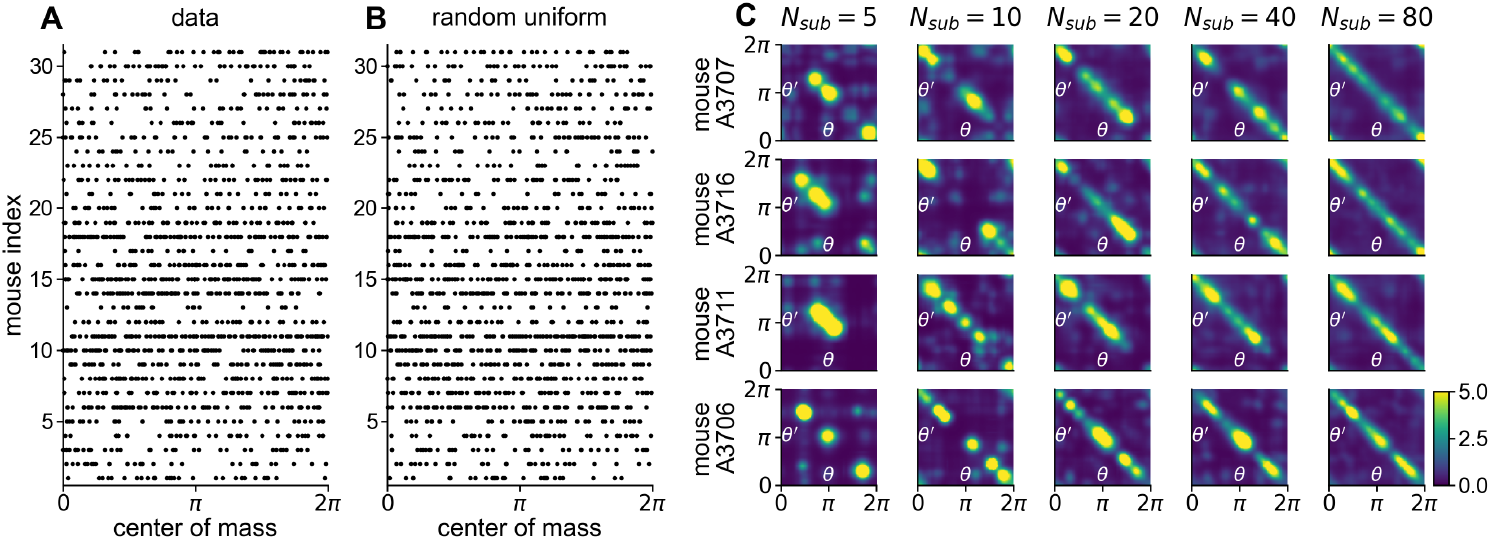
Evidence of distributional circular symmetry in head-direction cell responses (*N*_data_ = 1533 neurons across 31 mice). **(A)** Distribution of center-of-mass angles across neurons. Each row represents a different mouse, with dots indicating the center-of-mass angles of individual tuning curves. **(B)** Random uniform angle distributions matched to the number of neurons in each session, demonstrating visual similarity to the experimental data in (A). Statistical analysis confirms that the experimental distributions are consistent with uniformity (main text). **(C)** Empirical uncentered two-point correlation functions, *C*^*ϕ*^(*θ, θ*^*′*^) (Eq. 4), computed from three example mice, shown for increasing subset sizes (*N*_subset_). The correlation structure becomes increasingly circulant as more neurons are included.

As a stronger test of this distributional symmetry, we examined the two-point correlation function of tuning curves, *C*^*ϕ*^(*θ, θ*^*′*^) (Eq. 4). When computed for increasingly large random subsets of neurons (again, within individual mice), this function appears to approach a circulant form (Fig. 4C), consistent with circular symmetry. The same behavior is seen in normalized correlation functions (i.e., correlation rather than covariance functions), the eigenvectors of which converge to Fourier modes (Fig. B4), as expected for circulant matrices.

### 3.2 Structure of the generative process

Motivated by these observations, we developed a two-step generative process for sampling tuning curves. This is summarized in B3.1 (Setting 2). The first step involves sampling an input-current tuning curve *x*^⋆^(*θ*) from a circularly symmetric Gaussian process with zero mean and correlation function Γ^*x*^(Δ*θ*),

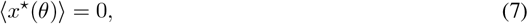

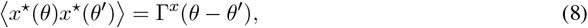

where the correlation function depends solely on the angular difference Δ*θ* = *θ* − *θ*^*′*^ in accordance with the evidence for distributional circular symmetry presented above. It is parameterized as a wrapped Gaussian,

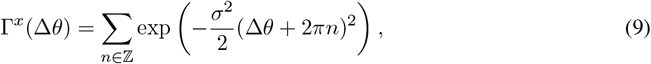

where *n* runs over the integers ℤ. Here, *σ* controls the typical number of peaks in the generated tuning curves (Fig. 5A, B), or equivalently, the number of large Fourier coefficients in these tuning curves (in particular, the Fourier coefficients are given by 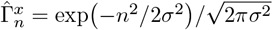 Sec. A6). The second step transforms these input currents into firing rates using a normalized softplus function (Fig. 5C; Sec. B3.1),

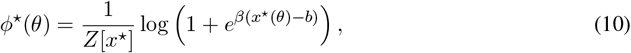

where *β* controls the sharpness of the activation function and *b* sets a soft threshold. *Z*[*x*^⋆^] is a functional that normalizes the mean over *θ* of the firing rate to unity, serving two purposes. First, it places all tuning curves on the same scale (across different mice within the data, and between these data and the generative process). Second, tuning curve features result from positive deviations of the input-current Gaussian process; for *b >* 0, these deviations can be small and require amplification through normalization to produce pronounced peaks.

**Figure 5:**
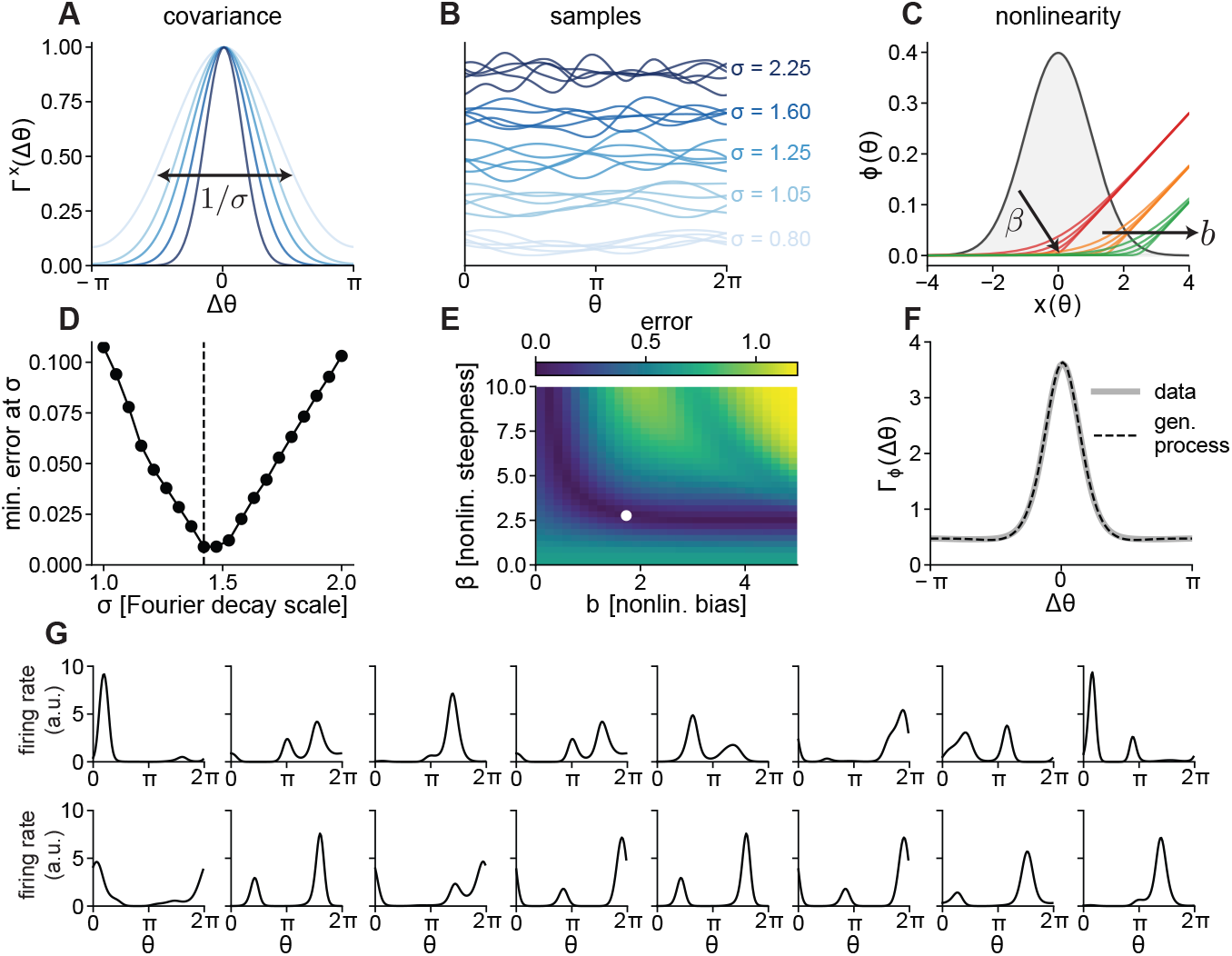
Statistical generative process for head-direction tuning curves. **(A)** Angular correlation functions Γ^*x*^(Δ*θ*) for different values of the Fourier decay scale *σ*. **(B)** Example samples from the Gaussian process for different values of *σ*. Increasing *σ* makes sampled tuning curves more wiggly. **(C)** Effect of activation function parameters *β* (sharpness) and *b* (threshold) on the input-output relationship. Gray shading shows the Gaussian distribution of input currents. **(D)** Parameter selection curve showing the minimal error (minimized over *b* and *β*) as a function of *σ*. Error measures the difference between generative process and experimental two-point correlation functions across the first 20 distinct Fourier coefficients. **(E)** Error landscape at optimal *σ* as a function of *b* and *β*. **(F)** Two-point correlation functions of the data and generative process agree at the optimal parameter values. **(G)** Random samples from the optimized generative process, exhibiting heterogeneity visually reminiscent of experimental tuning curves.

### 3.3 Quantitative validation of the generative process

Rather than fitting session-specific parameters, we sought a single parameter set (*σ, β, b*) for the generative process that could capture the statistical structure across all recordings. We fit these parameters by minimizing the difference between the empirical two-point correlation function, pooled across mice, and the generative correlation function, obtaining fitted values (*σ, β, b*) ≈ (1.42, 2.76, 1.73) (Fig. 5D, E, F). The optimized generative process yields tuning curves that closely resemble the experimental observations (Fig. 5G).

To quantitatively validate the generative process, we compared several statistical features of generated and experimental tuning curves (Fig. 6). First, we computed the center-of-mass-aligned mean tuning-curve profile, as well as the profile of fluctuations, for the generative process (as done for the data in Sec. 2.1). These predicted profiles match those observed experimentally (Fig. 6A, B).

**Figure 6:**
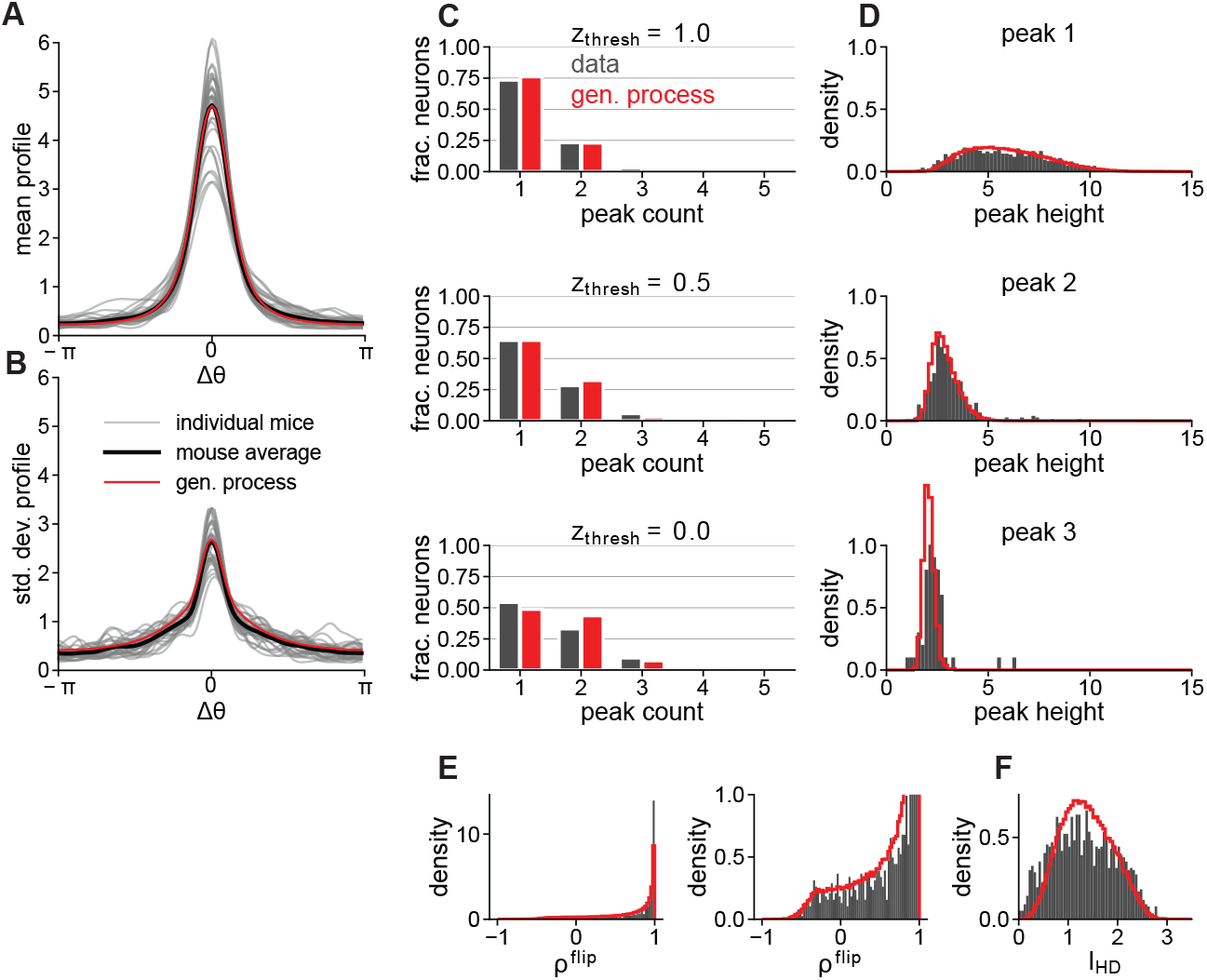
Quantitative validation of the generative process against experimental data (*N*_samples_ = 100,000 from generative process, *N*_data_ = 1533 from experiments). **(A)** Mean tuning-curve profile predicted by the generative process overlaid on experimental data from Fig. 2E (individual mice, mouse average). **(B)** Standard deviation of tuning curves around mean profile predicted by the generative process overlaid on experimental measurements from Fig. 2F (individual mice, mouse average). **(C)** Distribution of peak counts per tuning curve for decreasing detection thresholds (*z*_thresh_, top to bottom). Here and in the following panels (D, E, F), experimental data (gray) and generative process samples (red) are compared. **(D)** Distributions of tuning curve values at the three highest local maxima. **(E)** Distribution of flip symmetry correlations (*ρ*^flip^, left), with an expanded view of the asymmetric tail (right). **(F)** Distribution of head-direction information content (*I*_HD_).

We next analyzed the distribution of peak counts in individual tuning curves, where a peak is counted only if its *z*-score (computed using the tuning curve’s own mean and standard deviation) exceeds a detection threshold *z*_thresh_ (Fig. 6C). The model accurately reproduces the experimental peak-count distributions, with particularly strong agreement at higher detection thresholds *z*_thresh_.

We further characterized the peak structure by comparing the full distributions of tuning-curve values at the first, second, and third highest peaks, using *z*_thresh_ = 1 (Fig. 6D). Again, we find close agreement, now for full distributions, between the generative process and the data.

Going beyond peak structure, we examined the deviation from reflection symmetry in individual tuning curves. For each tuning curve, we computed the correlation *ρ*^flip^ between the original tuning curve and its reflection around its circular center of mass. The model accurately reproduces the distribution of these correlations (Fig. 6E, left), capturing particularly well the tail of strongly asymmetric tuning curves (Fig. 6E, right).

Finally, we compared the distributions of head-direction information content *I*_HD_, a commonly used measure of how precisely individual neurons encode head direction (Fig. 6F) [54]. The model shows good agreement with the data, especially for the tail of neurons with high information content.

In summary, the generative process accurately reproduces an array of features of the experimental tuning curves, indicating that the parsimonious three-parameter model has identified universal statistical structure present in the rodent head-direction system. The particularly strong agreement in the tails of these distributions suggests possible universal extreme value statistics at play, though agreement is seen for non-extreme quantities as well.

## 4 Large-network limit: weights

Inspired by the success of the “Gaussian plus nonlinearity” generative process in capturing tuning curve statistics, we now use a similar generative process to study optimal weight matrices and network dynamics at large *N*, typical of mammalian circuits. We retain the input-current correlation function Γ^*x*^(Δ*θ*) with rotational symmetry and Fourier decay scale *σ*. For analytical tractability, we adopt the saturating activation function 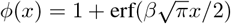, where *β* controls response sharpness. This choice enables closed-form evaluation of certain integrals. From a dynamical perspective, saturation at 0 and 2 for large negative and positive inputs, respectively, precludes runaway activity, removing the need for the uniform inhibition parameter *c* used in Sec. 2.3. Finally, since we have eliminated the soft threshold *b* used in Sec. 3.2, firing rates are non-sparse over *θ* and the normalization functional *Z*[*x*^⋆^] is not needed to amplify small tuning features. This error-function generative process has two parameters, *σ* and *β*, and is summarized in Sec. B3.1 (Setting 3).

We now analyze the optimal weight matrix ***J*** at large *N* (in this section, we use bold symbols for vectors and matrices). We first point out that, by virtue of the distributional circular symmetry in the generative process, the target manifold possesses a circular geometry at large *N*; that is, the distributional symmetry becomes a geometric symmetry. We then show that the optimal weights that encode the target manifold reflect this circular geometry in their eigenvalues, which exhibit doublet degeneracy. A general analysis of the relationships between continuous symmetries, eigenvalue degeneracies, and symmetries of the dynamics is provided in Sec. A1. This spectral structure addresses a corollary to question (2), identifying which connectivity features are conserved between classical and heterogeneous continuous-attractor models. Details are provided in Sec. A3.2. Note that while we analyze the optimal weight matrix using the generative process, the same structure derived here holds in the data-derived weight matrix (Fig. B5).

The target manifold consists of the desired patterns of input currents 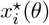 and firing rates 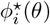 across neurons. We organize these patterns into matrices ***X*** and **Φ** of size *N*_*θ*_*× N*, where *θ* has been discretized into *N*_*θ*_ bins. Although tuning curves contain infinitely many nonzero Fourier components, their smoothness allows us to truncate at frequency *D* without introducing significant error, where *D* remains fixed as *N* increases. This truncation ensures that both matrices have rank 2*D* + 1, admitting singular value decompositions

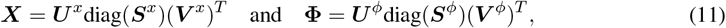

where 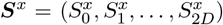 and 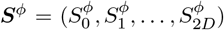 are vectors of singular values. The left singular vectors comprise the columns of matrices ***U*** ^*x*^, ***U*** ^*ϕ*^ of size *N*_*θ*_ *×* (2*D* + 1), and the right singular vectors comprise the columns of matrices ***V*** ^*x*^, ***V*** ^*ϕ*^ of size *N ×* (2*D* + 1). These singular-vector matrices have orthonormal columns.

As *N* → ∞, the singular value decompositions become Fourier decompositions. To see this, note that the empirical *N*_*θ*_ *× N*_*θ*_ covariance matrices converge to

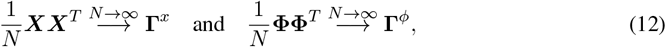

where the *N*_*θ*_ *× N*_*θ*_ matrices **Γ**^*x*^ and **Γ**^*ϕ*^ are circulant. Their (*θ, θ*^*′*^) elements are given by Γ^*x*^(*θ* − *θ*^*′*^) (the generative process input-current correlation function) and 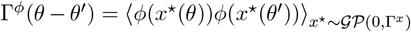 (the generative process firing-rate correlation function).

The left singular vectors ***U*** ^*x*^ and ***U*** ^*ϕ*^ are equivalently eigenvectors of the respective empirical covariance matrices and thus converge as *N*→ ∞ to eigenvectors of **Γ**^*x*^ and **Γ**^*ϕ*^, which are the Fourier modes: ***U*** ^*x*^ = ***U*** ^*ϕ*^ = ***F***, where the columns of ***F*** consist of the constant mode followed by sine-cosine pairs for each frequency from 1 to *D*. We adopt this sine-cosine Fourier basis (with mode index *n* = 0, …, 2*D*) in this section to make the singular vectors real; otherwise, we prefer the complex-exponential basis (with *n* = −*D*, …, *D*; Sec. A3.1).

The singular values ***S***^*x*^ and ***S***^*ϕ*^ converge to the square roots of the eigenvalues of **Γ**^*x*^ and **Γ**^*ϕ*^, respectively. These eigenvalues are the Fourier coefficients of the respective correlation functions, denoted by 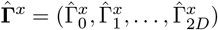 and 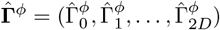. Due to circular symmetry, consecutive pairs of coefficients are equal (excluding the constant mode *n* = 0), exhibiting a doublet degeneracy.

Finally, the right singular vectors embed the Fourier modes, which comprise a geometric ring, into the neuronal spaces of input currents and firing rates. Specifically, the *i*-th row of ***V*** ^*x*^ contains the 2*D* + 1 Fourier coefficients of 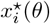 scaled to have unit variance, and similarly for ***V*** ^*ϕ*^. These coefficients are real and expressed with respect to the sine-cosine basis defined by ***F***.

### 4.1 Spectral structure of J

The optimal weight matrix ***J*** connects the target manifolds of input currents and firing rates. The least-squares solution (Eq. 3) can be written in matrix form as 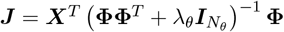, where we additionally dropped the constraint *J*_*ii*_ = 0 in anticipation of the large-*N* limit and defined *λ*_*θ*_ = *N*_*θ*_*λ*. Replacing ***X*** and **Φ** with their singular value decompositions and applying a linear-algebra identity (Sec. A2) gives

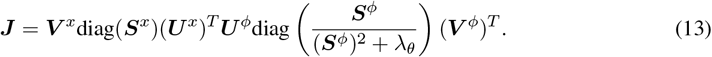

Taking the *N→ ∞* limit and applying the relation (***U*** ^*x*^)^*T*^ ***U*** ^*ϕ*^ = ***F*** ^*T*^ ***F*** = ***I***_2*D*+1_, then taking *λ*_*θ*_ → 0 to yield the minimum-norm solution, gives the singular value decomposition of ***J*** :

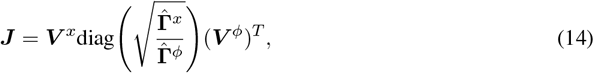

where ***V*** ^*x*^ and ***V*** ^*ϕ*^ are the left and right singular vectors, respectively, of ***J***.

Using the relation (***U*** ^*x*^)^*T*^ ***U*** ^*ϕ*^ = ***I***_2*D*+1_ from above, one can show that the left-right singular-vector overlaps are diagonal, with

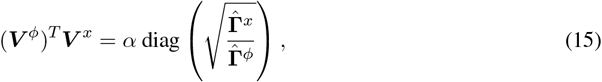

where

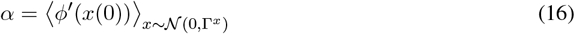

is the average slope of the neuronal activation function (Sec. A2). Consequently, ***V*** ^*x*^ and ***V*** ^*ϕ*^ comprise the right and left eigenvectors, respectively, of ***J***, with eigenvalues

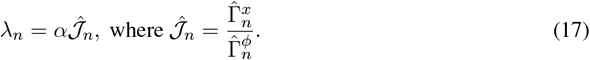

These eigenvalues are real and exhibit doublet degeneracy, features inherited from 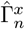 and 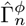. This degeneracy directly reflects the geometric symmetry of the activity manifold that the network generates, which is itself a consequence of the distributional symmetry of tuning curves at large *N→ ∞* (Fig. 7A, C, D). These features of the eigenvalues are characteristic of a matrix that is transpose-symmetric and circulant, despite the fact that ***J*** lacks both of these features. Indeed, ***J*** maintains a finite level of transpose-asymmetry even as *N* (Sec. A4.5). Nevertheless, these features suggest that the dynamical behavior of the system governed by ***J*** is related to that of a classical ring-attractor model with transpose-symmetric circulant weights. Furthermore, a reasonable conjecture is that the effective interactions in the effective classical model would be proportional to 𝒥(Δ*θ*), given by the inverse Fourier transform of 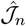. We show that this is correct in Sec. 5.

**Figure 7:**
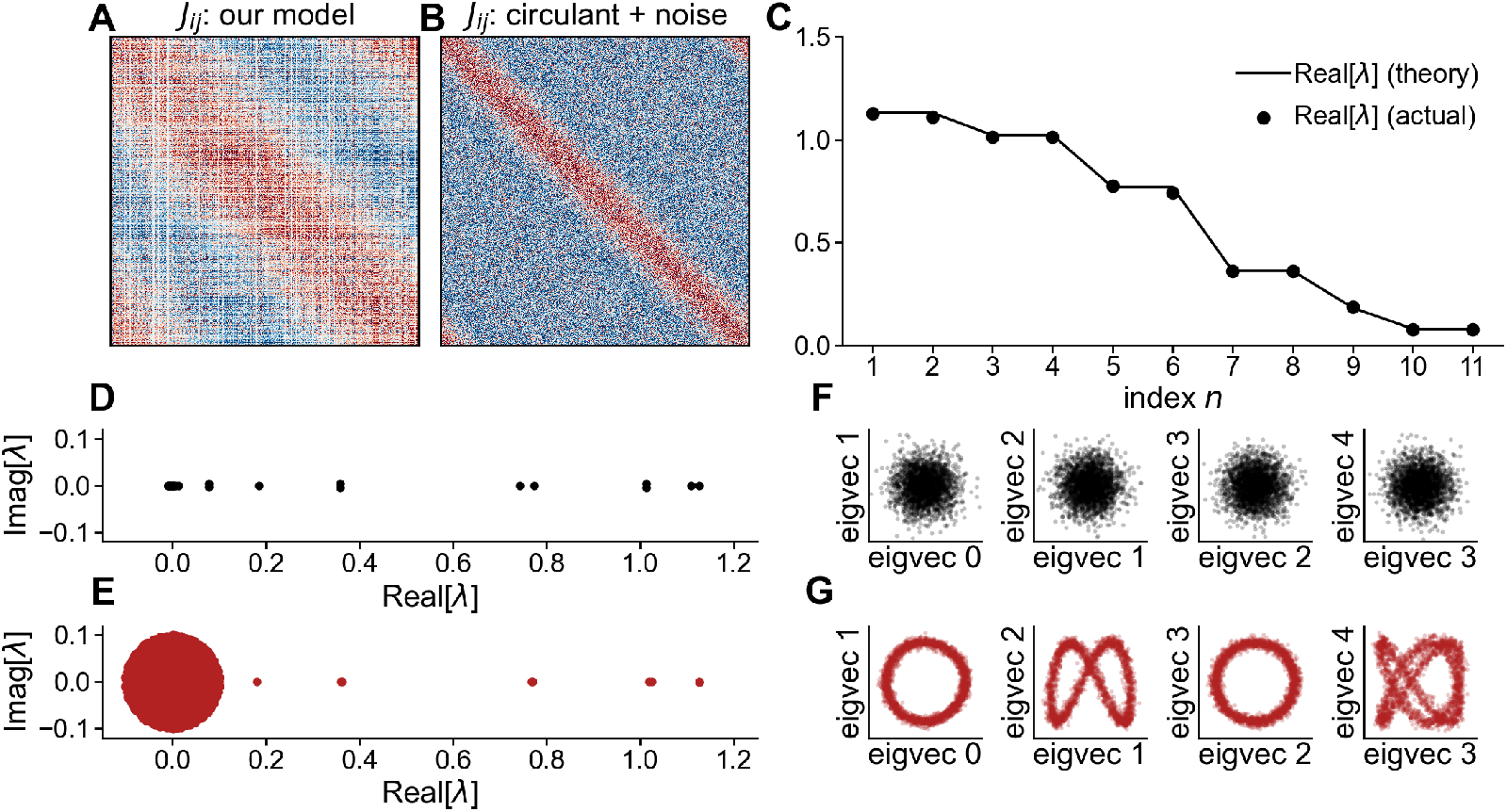
Spectral structure of the optimal weight matrix (*N* = 2500, generative-process Fourier decay scale *σ* = 1.42, response sharpness *β* = 2.76). **(A)** Optimal weight matrix using tuning curves drawn from the generative process. Neurons are ordered by their tuning-curve centers of mass. **(B)** Circulant matrix with added noise, where the circulant component has the same eigen values as ***J*** at large *N* (Eq. 17). Matrix elements are of order 1*/N*. Independent Gaussian noise with standard deviation 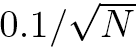 is added to this circulant matrix. **(C)** Sorted real parts of the eigenvalues of the optimal matrix from (A). Theoretical predictions (lines) are shown alongside numerical results (dots). In this example, mode 9 is the constant mode and thus does not occur in a doublet. **(D)** Complex-plane view of eigenvalues of the optimal weight matrix from (A), with real parts as in (C). **(E)** Complex-plane view of eigenvalues from the circulant-plus-noise matrix in (B). **(F)** Scatterplots comparing the components of consecutive pairs of eigenvectors of ***JJ*** ^*T*^ (equivalently, the left singular vectors of ***J***), where ***J*** is the optimal matrix from (A). No structure is apparent. **(G)** Analogous scatterplots for the circulant-plus-noise matrix from (B), where the leading eigenvectors retain their Fourier structure despite partial corruption by noise.

### 4.2 Disordered vs. ordered embeddings and implications for connectome analysis

Because the spectrum of ***J*** is, at large *N*, identical to that of a transpose-symmetric circulant matrix (as in a classical ring-attractor model), the difference between such a matrix and ***J*** must lie in the eigenvectors. For circulant weights, the eigenvectors are Fourier modes: in our notation, ***V*** ^*ϕ*^ = ***V*** ^*x*^ = ***F***. The eigenvalue structure (real, doubly degenerate) combined with Fourier eigenvectors produces a circulant matrix. In contrast, ***J*** has the same eigenvalue structure but disordered eigenvectors ***V*** ^*x*^ and ***V*** ^*ϕ*^, obscuring the structure present in the spectrum.

We now describe the nature of these disordered eigenvectors more precisely. Recall that, as *N → ∞*, ***V*** ^*x*^ and ***V*** ^*ϕ*^ contain Fourier coefficients of tuning curves (scaled to have unit variance). The elements of ***V*** ^*x*^ (whose *i*-th row contains the Fourier coefficients of 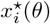 exhibit statistical independence across both the rows and columns. This double independence arises because each neuron’s tuning curve is sampled independently from the same generative process (independence across rows, *i* = 1, …, *N*) and the Gaussian process construction implies that different Fourier coefficients are uncorrelated by definition (independence across columns, *n* = 0, …, 2*D*). While the elements of ***V*** ^*ϕ*^ (whose *i*-th row contains the Fourier coefficients of 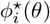 are similarly independent across rows, they are not independent across columns. This is because applying the nonlinear activation function in *θ*-space induces higher-order correlations in Fourier space, although pairwise correlations vanish due to rotation invariance. Equivalently, 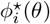 constitutes a non-Gaussian process, which generically has non-independent Fourier coefficients.

This disorder has important implications for connectome analysis. Consider the challenge of detecting ring-attractor structure in an observed synaptic weight matrix. A common method is spectral embedding, which involves computing the top eigenvectors of symmetric matrices derived from the weights, assigning coordinates to each neuron based on these eigenvectors, and visualizing the resulting coordinates to detect geometric structure. Candidate matrices include ***JJ*** ^*T*^ and ***J*** ^*T*^ ***J***, which measure similarity of neuronal inputs and outputs, respectively. Their eigenvectors are the left and right singular vectors of ***J***, converging to ***V*** ^*x*^ and ***V*** ^*ϕ*^ as *N→ ∞*, respectively. Although this method successfully reveals rings in circulant matrices, where the eigenvectors are Fourier modes, it fails for our model because both ***V*** ^*x*^ and ***V*** ^*ϕ*^ are disordered (Fig. 7F, Fig. B6). Despite ***V*** ^*ϕ*^ having some structure in the form of higher-order correlations between entries (Fig. B6), the resulting scatter plots appear similarly disordered to those from ***V*** ^*x*^. In Sec. A3.3, we attempt to extract information about the latent ring structure from the eigenvectors despite their disorder, finding that this approach hallucinates spurious patterns from low-rank structure alone. The ring-attractor models of Beiran et al. [36] and Mastrogiuseppe and Ostojic [35] and of Pollock and Jazayeri [34] also feature disordered embeddings; one could also construct such a disordered-embedding ring-attractor model within the Neural Engineering Framework (see Barak and Romani [8], though they used an ordered embedding; Sec. A5).

A natural alternative model, also explored in the data-derived network case (Sec. 2.3, Fig. 3D,I) is one in which a circulant weight matrix is added to a background of independent, random weights. While 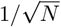 noise affects the eigenvalue spectrum by adding a bulk (Fig. 7E) [55], the eigenvectors retain their Fourier structure provided the noise is not too strong (Fig. 7G). Similar Fourier-structured eigenvectors arise in more sophisticated models where the added low-rank component is correlated with random background, such as that of Darshan and Rivkind [32] (Sec. 5.4). In our model, by contrast, the eigenvectors are structureless by construction. Eigenvalue degeneracy is present in all of these cases.

In summary, we propose that eigenvalue degeneracy is the most reliable signature for detecting ring-attractor structures in synaptic weight matrices, such as those from future connectome data. Unlike eigenvectors, this spectral structure is conserved between classical continuous-attractor models and various other models since this degeneracy directly encodes the geometry of the activity manifold.

## 5 Large-network limit: dynamical mean-field theory

The previous section identified conserved connectivity features between classical continuous-attractor models and our model, but did not relate their dynamics. In this section, we develop a theory of the full network dynamics in the limit *N → ∞* using dynamical mean-field theory. This answers question (2) by showing that all ingredients of classical ring-attractor models are recovered in the effective description: a nonlinear activation function (though modulated by global activity, as we will explain), Mexican-hat interactions, and, crucially, continuous symmetry. The dynamical mechanism, spontaneous breaking of continuous symmetry leading to bump states, is therefore conserved. We present the main components here, with details in Sec. A4.1. We continue to use the error-function generative process (Setting 3; Sec. B3.1). Convergence of the dynamics to this limiting behavior with increasing *N* is visualized in Fig. B9.

The dynamical mean-field theory reduces the high-dimensional network dynamics to equations governing the dynamics of a low-dimensional order parameter, 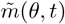, which is a temporally low-pass filtered version of the overlap order parameter *m*(*θ, t*) defined in Eq. 6. Specifically, for heading movements with small angular acceleration (*τ*^2^|∂_*t*_*ω*(*t*)| ≪ 1, which is satisfied by the behavioral conditions), this order parameter obeys

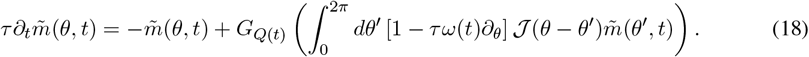

This equation is identical to that of classical ring-attractor models, with circulant interactions 𝒥 (Δ*θ*) and activation function *G*_*Q*_(*h*) [2]. We now unpack each of the components.

### 5.1 Interactions

The effective interactions appearing in Eq. 18 are in general proportional to **Γ**^*x*^(**Γ**^*ϕ*^)^*−*1^, where **Γ**^*x*^ and **Γ**^*ϕ*^ are the *N*_*θ*_ *× N*_*θ*_ input-current and firing-rate correlation matrices, respectively. When these are circulant, the effective interactions themselves are circulant and defined in Fourier space by coefficients 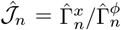, which are proportional to the eigenvalues of *J* as anticipated (Eq. 17). We denote the circulant effective interactions by 𝒥(Δ*θ*), visualized in Fig. 9A. In addition to being circulant, these interactions are transpose symmetric, 𝒥(Δ*θ*) = 𝒥(*−* Δ*θ*). In contrast, the full network weights *J*_*ij*_ are non-circulant and lack transpose symmetry, with *J*_*ij*_ ≠ *J*_*ji*_ even as *N→ ∞*, where corr(*J*_*ij*_, *J*_*ji*_) = *α*^2^[𝒥^2^](0)*/*𝒥(0) (Sec. A4.5).

**Figure 8:**
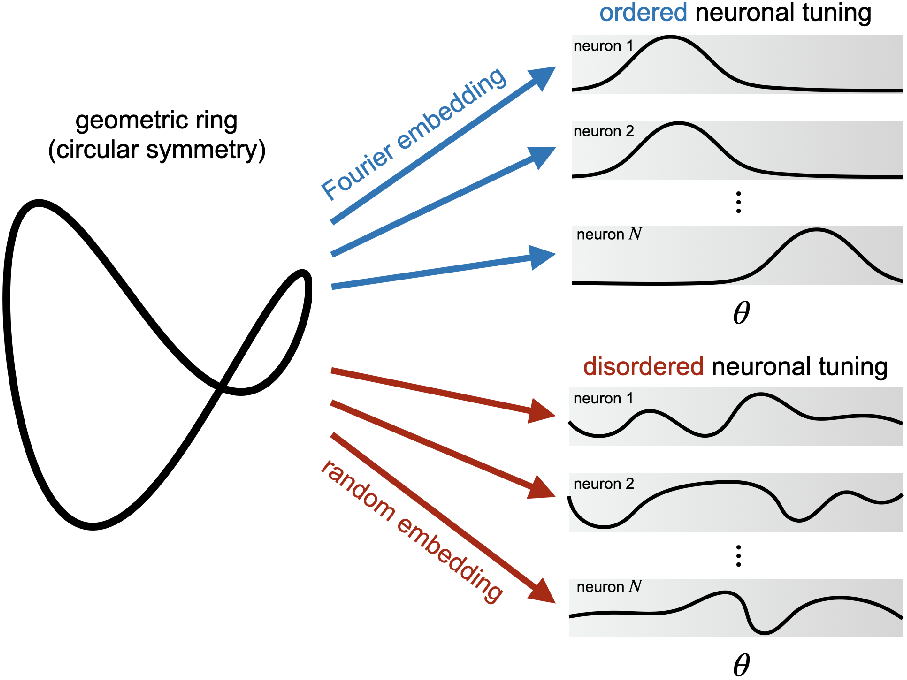
Both classical continuous-attractor models (top) and our model (bottom) generate circular activity manifolds with identical geometry. Classical models achieve this through ordered Fourier patterns across neurons, producing circulant weight matrices with Fourier eigenvectors. Our model achieves the same manifold geometry through disordered, random embeddings that produce heterogeneous tuning curves and weight matrices with disordered eigenvectors. Despite this difference in embedding, both models share identical weight-matrix eigenvalue structure (doublet degeneracy), reflecting the circular geometry of the activity manifold.

**Figure 9:**
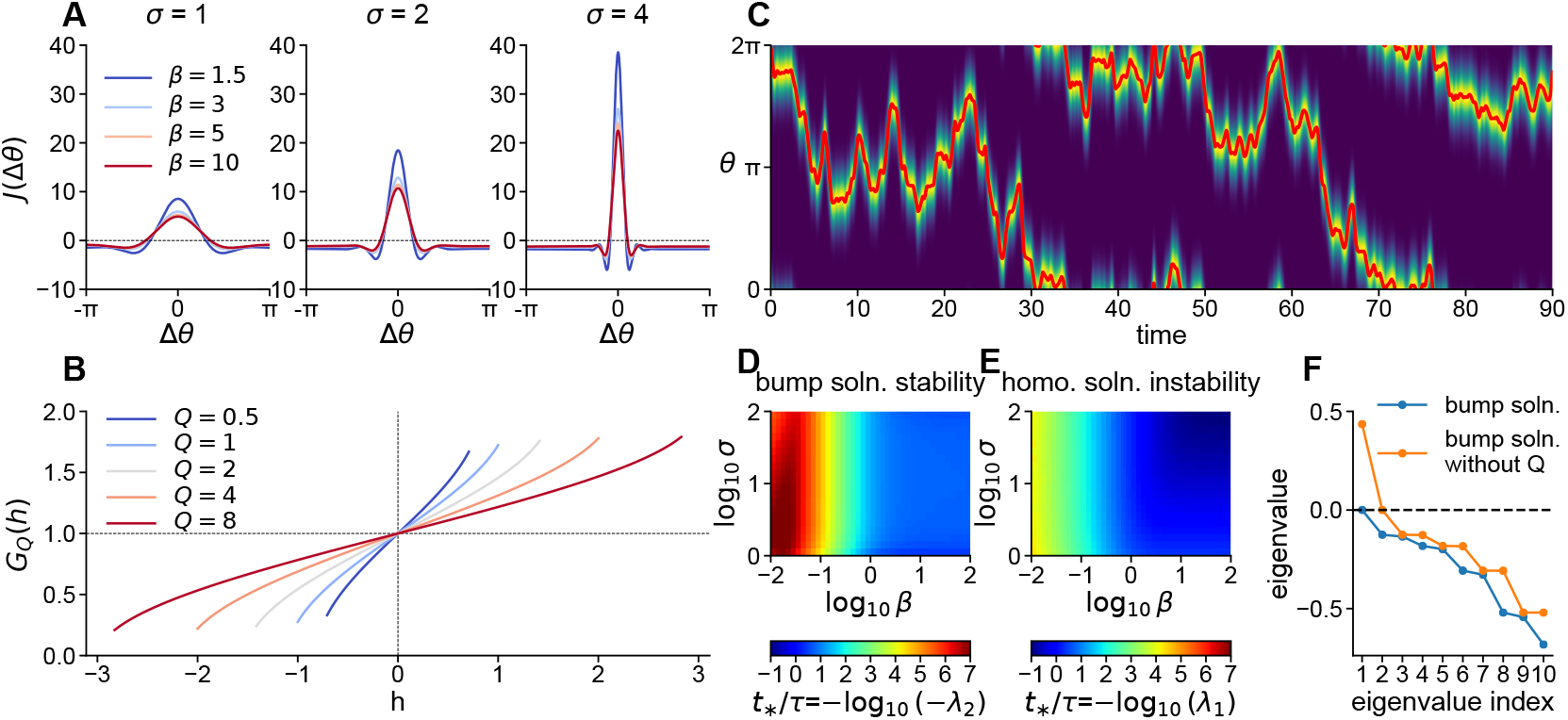
Dynamical mean-field theory recovers classical ring-attractor dynamics in the large-*N* limit. **(A)** Mean-field interactions 𝒥(Δ*θ*) are circulant and exhibit a Mexican-hat profile that varies with the Fourier decay scale *σ* (panels) and response sharpness *β* (curves). Higher *β* increases the amplitude of excitation and inhibition, while larger *σ* narrows the profile. **(B)** Activation function *G*_*Q*_(*h*) for different values of the global activity-dependent parameter *Q*. As network activity increases, *Q* grows, reducing the slope of the activation function. This function has a finite domain for *h* (Sec. A4.4). **(C)** Solution of the dynamical mean-field theory equations showing a traveling bump tracking head-direction angular velocity input, matching the dynamics observed in the high-dimensional network. **(D)** Timescale of convergence to the ring manifold (*t*_*\**_*/τ* = −log(−*γ*_2_)) as a function of *σ* and *β*, where *γ*_2_ is the eigenvalue with second-largest real part of the Jacobian *M* (*θ, θ*^*′*^) evaluated at the bump state. Rapid convergence (*t*_*\**_ ∼ *τ*) occurs for *β* ≳ 1 across all *σ*. **(E)** Timescale of divergence from the uniform state (*t*_*\**_*/τ* = −log(*γ*_1_), where *γ*_1_ is the leading eigenvalue of the Jacobian evaluated at the uniform state), demonstrating a pattern-forming (Turing) instability. **(F)** Real parts of the leading eigenvalues of the Jacobian *M* (*θ, θ*^*′*^) at the bump state for the full dynamical mean-field theory equations and modified equations with fixed *Q* = Γ^*x*^(0). Dynamic *Q*(*t*) is required for bump stability; fixing *Q* = const. introduces a positive eigenvalue that destabilizes the bump.

The interactions 𝒥(Δ*θ*) exhibit a Mexican-hat profile characteristic of pattern-forming systems, with shape controlled by the generative-process Fourier decay scale *σ* and activation function sharpness *β*, as shown in Fig. 9A. This shape emerges from the statistical structure of tuning curves (Sec. A4.6, Fig. A3), with the excitatory peak and inhibitory surround arising from second and fourth moments, respectively. Increasing *β* amplifies both excitation and inhibition, while larger *σ* produces a narrower profile.

### 5.2 Activation function

The activation function *G*_*Q*_(*h*) in Eq. 18, defined in Eq. 69, is modulated by a factor *Q* that measures the level of global excitation across angles *θ*. Specifically, at time *t, Q*(*t*) is a quadratic functional of the form

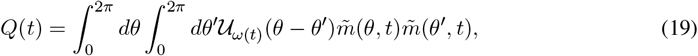

where 𝒰_*ω*(*t*)_(Δ*θ*) is an angular velocity-dependent kernel (Sec. A4.1).

Both *G*_*Q*_(*h*) and the single-neuron sigmoidal activation function *ϕ*(*x*), present in both classical ring-attractor models and our *N*-dimensional model, exhibit local slope (or “gain”) dependence on their inputs: ∂_*h*_*G*_*Q*_(*h*) and ∂_*x*_*ϕ*(*x*) are not constant functions of *h* and *x*, respectively. However, *G*_*Q*_(*h*) has an additional global gain dependence on activity across all *θ* through *Q*. As *Q* increases, the local gain ∂_*h*_*G*_*Q*_(*h*) decreases for all values of *h*, as shown in Fig. 9B. A further difference is that, for fixed *Q*, ∂_*h*_*G*_*Q*_(*h*) increases monotonically with |*h*|, while ∂_*x*_*ϕ*(*x*) decreases monotonically with |*x*| due to saturation. The global gain control in *G*_*Q*_(*h*) is necessary to stabilize the system, compensating for the absence of local saturation. Despite these differences, both mechanisms prevent runaway activity through gain control that depends on the instantaneous state of the system and are therefore similar in this regard. As discussed below, this instantaneous state dependence is not a universal feature of dynamical mean-field theories.

### 5.3 Solutions

We now discuss solutions of the dynamical mean-field equations and their stability. For *ω*(*t*) = 0 the equations exhibit static solutions 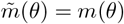. The stable solutions are bump states of the form

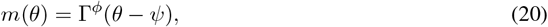

where *ψ* sets the angular location of the peak of the bump, reflecting the rotation-invariance of the dynamical mean-field equations. For *ω*(*t*) ≠ 0, there are traveling-bump solutions,

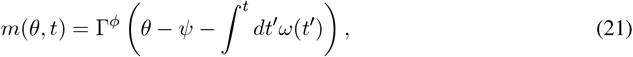

with 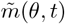 following a temporally smoothed version of this profile. These solutions track head direction through integration of angular velocity (Fig. 9C), matching the behavior observed in both network simulations (Fig. 3K, L) and experimental recordings (Fig. 3M). We note that, for both static and traveling-bump solutions, *Q* = Γ^*x*^(0), independent of time.

The stability properties and related timescales of the mean-field dynamics can be analyzed through the Jacobian 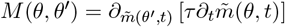 (Sec. A4.3). When evaluated at a particular equilibrium state 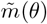, the absence or presence of an eigenvalue with a positive real part indicates stability or instability, respectively. We always expect a single zero eigenvalue due to the continuous symmetry of the system.

For a bump state 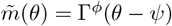 (where *ψ* is arbitrary and can be set to *ψ* = 0), diagonalization of *M*(*θ, θ*^*′*^) reveals that all eigenvalues have negative real parts except for a single zero eigenvalue corresponding to rotation symmetry (Fig. 9F, blue curve). The ultimate convergence timescale to the ring manifold is determined by the eigenvalue *λ*_2_ with the second-largest real part, which is itself real. Specifically, perturbations decay with a characteristic timescale *t*_*_*/τ* = − log(−*λ*_2_) (Fig. 9D). The timescale *t*_*_ is of order *τ* (i.e., convergence is fast) for *β* ≳ 1.

The system exhibits a Turing instability that destabilizes the uniform state 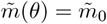, where 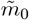 can be determined numerically. Diagonalizing the Jacobian reveals a real, positive eigenvalue *λ*_1_ for all *σ* and *β*, with the instability timescale *t*_*_*/τ* = − log(*λ*_1_) (Fig. 9E). This timescale *t*_*_ is of order *τ* (i.e., destabilization is fast) for *β* ≳ 1, with a weak dependence on *σ*.

The global gain control through *Q*(*t*) is essential for these pattern-forming dynamics. Suppose we fix *Q* = Γ^*x*^(0), removing its dynamical dependence on the system state (this manipulation has no analogue in the high-dimensional system as it would correspond to fixing the *L*^2^ norm of input currents while letting the overlap with the target manifold evolve as though this norm were not fixed). The result is that the bump becomes unstable, evidenced by the real, positive eigenvalue in Fig. 9F (orange curve). The system instead converges to a new uniform state 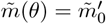, where 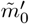 can be determined numerically.

Finally, we note that the overlap order parameter 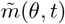 determines not only the macroscopic behavior of the system but also the dynamics of the local activities, through

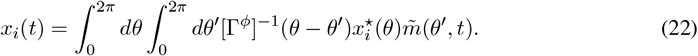

That is, neuronal activities are a simple linear embedding of the overlap order parameter.

In summary, the main components of the dynamical mean-field theory are

1. A nonlinear activation function *G*_*Q*_(*h*) that, unlike classical ring-attractor models, depends on a measure of the network’s instantaneous global activity *Q*(*t*) through an emergent gain-control mechanism.
2. Effective weights 𝒥(Δ*θ*) supporting Turing pattern formation, with local excitation and longer-range inhibition forming a Mexican-hat profile.
3. A continuous circular symmetry, reflected in the effective weights 𝒥(Δ*θ*) depending only on angular differences Δ*θ*, that is spontaneously broken, leading to the emergence of bump states.

As described in the introduction, these are the essential ingredients of a classical continuous-attractor model. Thus, the model we have constructed, whose tuning statistics share the fundamental structure indicated by experimental data, relies on the same dynamical mechanisms as classical continuous attractors. This provides a precise answer to question (2): classical continuous-attractor dynamics are not merely compatible with biological heterogeneity but are the exact effective description at the population level for systems with this statistical structure.

### 5.4 A taxonomy of dynamical mean-field theories

One way heterogeneous representations could arise in a recurrent neural network is through the combination of random initialization and task-driven learning. In this spirit, Rivkind and Barak [33] analyzed a two-stage reservoir-computing process [56, 57]: first, a low-rank addition to random background weights is learned to produce a desired set of discrete fixed points; then, the system operates autonomously in closed-loop mode. They computed a response function of the closed-loop system at large *N* whose poles control local stability around fixed points. We extended this analysis by deriving the dynamical mean-field description of the closed-loop dynamics using a path-integral formulation (Supplement: Dynamics of learned attractors). This description captures the global nonlinear dynamics, extending beyond the local linear analysis of Rivkind and Barak [33]. This theory involves order parameters given by overlaps of network states with learned patterns, analogous to order parameters in Hopfield networks. We recover the response-function poles computed by Rivkind and Barak [33] as eigenvalues governing the linearized dynamics of these overlaps.

The structure of the dynamical mean-field theory depends on the nature of the feedback weights that embed fixed points into neuronal space. For Gaussian feedback weights, the overlap dynamics reduce to an effective recurrent neural network with interactions determined by pattern statistics. For non-Gaussian feedback weights, such as Fourier embeddings, no such reduction occurs. Within the Gaussian case, the effective activation function depends on the history of the system’s global state through a two-time correlation function if *g >* 0, but depends only on the instantaneous global state if *g* = 0, yielding a more direct correspondence to an effective recurrent neural network. Note that *g* = 0 is necessary for the weights to be minimum-norm with respect to the tuning curves they produce.

When specialized to continuous circular manifolds of fixed points, this framework provides a unifying taxonomy of continuous-attractor models in which our model, classical models, the model of Darshan and Rivkind [32], and the model of Beiran et al. [36] and Mastrogiuseppe and Ostojic [35] appear as special cases. This taxonomy is based on three parameters: the rank *K* of the low-rank weights, the strength *g* of random background weights, and the embedding type (Gaussian versus Fourier). For models with Gaussian embeddings, such as ours, rotational symmetry of tuning curves produces rotational symmetry of the effective recurrent interactions.

Within this taxonomy, we pinpoint the differences between our model and that of Darshan and Rivkind [32]. Our model achieves tuning heterogeneity through multiple modes (*K >* 2), without random background weights (*g* = 0), and a Gaussian embedding. The Darshan and Rivkind [32] model achieves heterogeneity through two modes (*K* = 2), random background weights (*g >* 0), and a structured embedding. These differences have important conceptual and functional consequences described in Sec. S8 (see Sec. S8.1).

## 6 Grid cells

Our approach for constructing continuous-attractor models from heterogeneous tuning curves generalizes to symmetries of higher dimensions. We expect this method to extend to general Lie groups. Here we focus on a two-dimensional torus, demonstrating its application to grid cells in the medial entorhinal cortex, which are thought to support a two-dimensional toroidal attractor [22].

Classical continuous-attractor models of grid cells, such as the model of Burak and Fiete [49], involve a two-dimensional sheet of neurons with translation-invariant weights. These models generate multiple activity bumps through spatial pattern formation, with two-dimensional shifts of the pattern induced by movement in the environment. Alternative models with a single bump of activity on a twisted torus have also been proposed [59]. These classical models produce idealized, perfectly periodic hexagonal grid patterns. In contrast, experimental observations show that medial-entorhinal cells exhibit more complex, imperfect patterns (Fig. 10A). Grid cells are typically identified by selecting neurons with large “grid scores,” a metric quantifying the degree of hexagonal periodicity in spatial firing patterns. In data from various experiments, there often exists a continuous distribution of grid scores with no clear boundary between grid and non-grid responses (Fig. 10B). This contrasts sharply with classical continuous-attractor models, which predict high grid scores corresponding to a dedicated grid-cell circuit or “module.”

**Figure 10:**
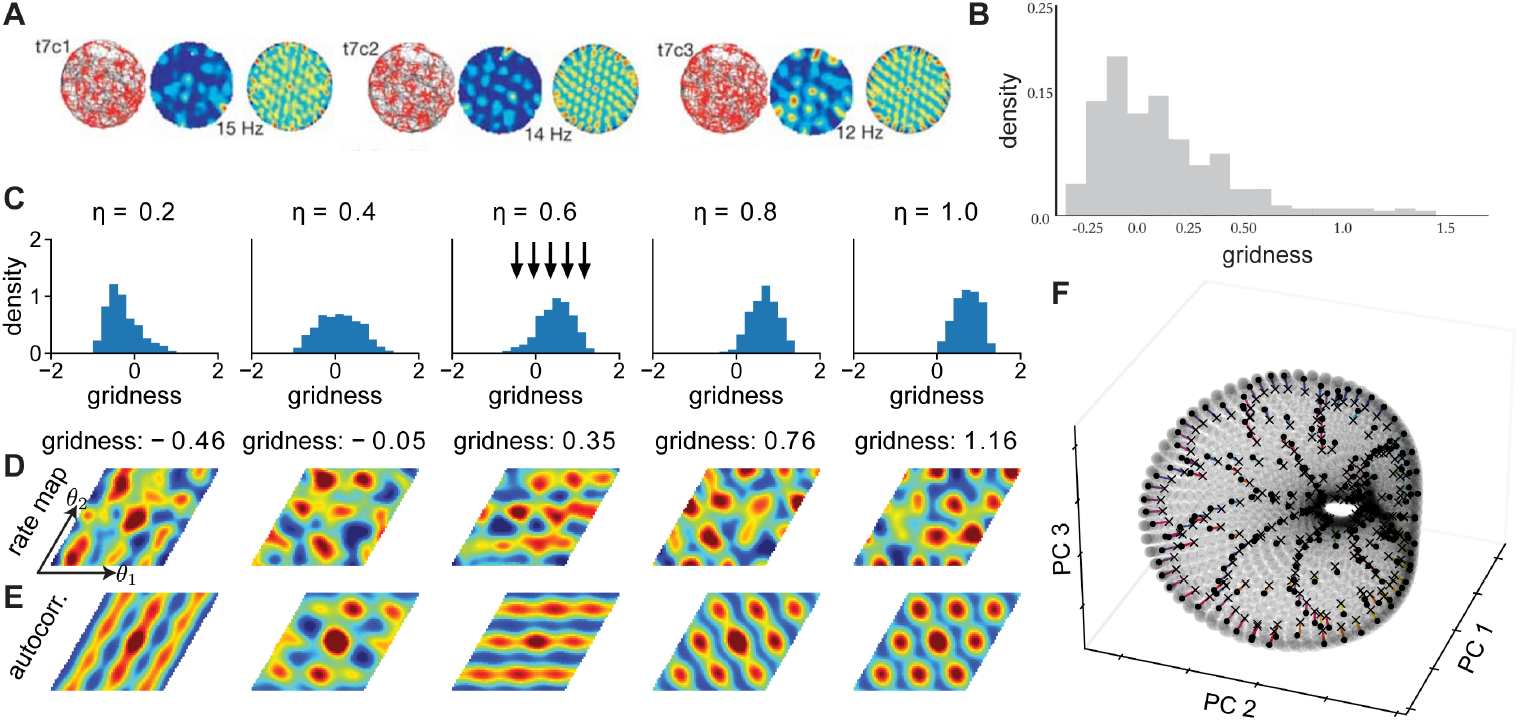
Grid-cell model derived from heterogeneous tuning curves. **(A)** Example experimental grid-cell responses, reproduced from Hafting et al. [16]. Each example shows spiking activity atop the rat’s two-dimensional spatial trajectory (left), the firing-rate map (middle), and the spatial autocorrelation (right). **(B)** Distribution of grid scores from experimental recordings, adapted from Nayebi et al. [25], which analyzed data from Mallory et al. [58]. **(C)** Grid-score distributions computed from input currents sampled from the Gaussian generative process for various values of the periodicity parameter *η* (*N*_*ϕ*_ = 80 per dimension, generative-process Fourier decay scale *σ* = 4, response sharpness *β* = 2.5, *K* = 3, *N*_samples_ = 1000). **(D)** Example input-current maps sampled from the Gaussian process with *η* = 0.6 (arrow in histogram in C). **(E)** Spatial autocorrelation functions computed from the input-current maps in (D), showing enhanced hexagonal structure compared to the raw maps in (D). **(F)** Network dynamics (*N* = 25,000, *σ* = 2.5, *K* = 1, error-function nonlinearity) visualized in the space of the first three principal components. Black crosses mark initial conditions, colored lines show trajectories over 30 s, and black dots indicate final states.

To generate more realistic grid-cell firing-rate maps, we extended the Gaussian process approach to periodicity in two dimensions. Let the periodic variables be a two-dimensional vector of angles, ***θ*** = (*θ*_1_, *θ*_2_). The correlation function of the Gaussian process is

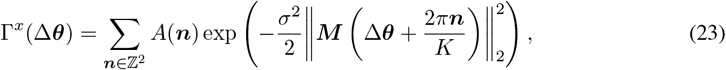

where ***n*** = (*n*_1_, *n*_2_) runs over the two-dimensional integer lattice ℤ^2^ and 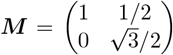 whose columns define the axes of a hexagonal lattice. The weighting function *A*(***n***) controls the periodicity of the patterns and is given by

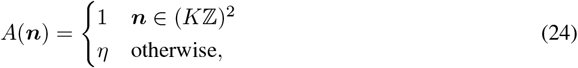

where 0 ≤ *η* ≤ 1 is the parameter controlling pattern periodicity. When *η* = 1, the sampled patterns are periodic with period 2*π/K*, which is the period of the grid activity pattern. For *η <* 1, the firing becomes only approximately periodic over this period but maintains periodicity over the larger period 2*π* and retains an underlying toroidal symmetry that is realized in the target manifold and network dynamics in the large-*N* limit. We set *K* = 3 for generating firing-rate maps and grid-score analyses.

Fig. 10D shows samples of firing-rate maps for various values of *η*, demonstrating increasing grid-like structure as *η* approaches unity. Notably, two-dimensional autocorrelograms of these firing fields (Fig. 10E) often display more pronounced hexagonal structure than the firing fields themselves, consistent with experimental observations [16]. Our approach provides a natural explanation for this phenomenon: the underlying periodic grid pattern is reflected in the autocorrelation function of the process generating the firing-rate maps, rather than in individual maps themselves. A “good grid cell” in our framework is one with a firing-rate map that just happens to resemble the correlation function of the generative process from which it was sampled.

To quantify the grid-like properties of the generated firing-rate maps, we computed grid scores using standard methods (Fig. 10C). A positive grid score indicates hexagonal structure, with higher scores representing more pronounced grid patterns. This approach generates a substantial number of cells that fail to meet the conventional criterion for grid cells (sometimes taken to be grid score *>* 0), despite being integral components of the toroidal attractor that we develop below. This suggests that the prevalent definition of grid cells may be overly restrictive, discarding neurons that play crucial roles in the emergent, functionally symmetric dynamics.

We constructed attractor networks from samples drawn from the generative process with *K* = 1, making the situation more directly analogous to the head-direction model. Using the same least-squares optimization approach described for head-direction cells, but extended to the two-dimensional case, we constructed an attractor network based on *N* = 25, 000 randomly drawn tuning curves. This larger network size is necessary because the number of neurons required for stable dynamics scales exponentially with the manifold dimension (2 in this case), so we need approximately the square of the neurons required for the head-direction system. The resulting network shows rapid convergence to the toroidal manifold from nearby initial conditions (Fig. 10F).

For grid-cell networks constructed using our approach, the weight matrix at large *N* exhibits eigen-value degeneracies in multiples of six, with the first few degeneracies being sixfold, reflecting the hexagonal symmetry of the grid lattice (Sec. A6, Fig. B11). This is analogous to the doublet degeneracy exhibited by head-direction networks.

When written in Cartesian coordinates ***x*** = ***Mθ***, the full *N* → ∞ mean-field description with velocity inputs takes a form similar to classical grid-cell models [49, 59],

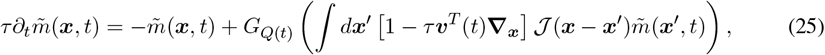

where ***v***(*t*) = (*v*_*x*_(*t*), *v*_*y*_(*t*)) is the Cartesian velocity and ***∇***_***x***_ = (∂_*x*_, ∂_*y*_) is the Cartesian gradient. This dynamical mean-field equation exhibits two-dimensional translation symmetry. Our use of *K* = 1 means that this is a single-bump model; the activity pattern exhibited by 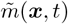 is a single bump of activity moving in response to velocity input.

## 7 Discussion

Classical continuous-attractor models rely on continuously symmetric weights to generate homogeneous tuning curves that tile a low-dimensional variable. Mammalian circuits exhibit far more heterogeneous tuning, raising two questions: (1) what structure, if any, governs this heterogeneity, and (2) how do networks generating such tuning relate to classical models? We addressed these questions by showing that heterogeneous tuning curves in data are consistent with nonlinearly transformed samples from a Gaussian process with rotationally symmetric correlation functions, and that these tuning curves can comprise a stable attractor manifold by recurrent neural networks whose weight matrix is minimum-norm subject to the tuning curve constraints. These networks admit a dynamical mean-field theory that recovers the essential features of classical ring-attractor models, demonstrating that classical continuous-attractor mechanisms could underlie heterogeneous tuning in mammalian circuits.

### 7.1 Is symmetry necessary?

The connection between our model and classical models relies on rotational symmetry of tuning curve statistics. This raises the question: is symmetry necessary in continuous-attractor models? Strictly speaking, no: the universal approximation property of recurrent neural networks [37] implies that networks of *N* neurons can support a continuous attractor manifold of arbitrary shape embedded in a *K*-dimensional subspace, provided that *K* is fixed and *N* can be taken arbitrarily large (Sec. A5).

What is the nature of the effective dynamics in such non-symmetric systems? Our dynamical mean-field framework extends to asymmetric cases and reveals how systems maintain a continuum of stable states without continuous symmetry. Fig. B7 examines systems with non-circulant correlation functions Γ^*x*^(*θ, θ*^*′*^) ≠ Γ^*x*^(*θ − θ*^*′*^), parameterized by an asymmetry factor *f* ≥ 0. As *f* increases, the effective weights 𝒥(*θ, θ*^*′*^) become increasingly non-circulant, yet the system maintains a one-dimensional continuum of stable states. Stability is preserved through compensatory modulation in the gain kernel 𝒰(*θ, θ*^*′*^) that appears in the quadratic term defining *Q*(*t*) in the effective activation function. The non-circulant gain modulation exactly compensates for the non-circulant structure of the weights, enabling continuous-attractor dynamics.

If symmetric manifolds are not strictly necessary, why might biological systems exhibit them? Head-direction circuits appear relatively rigid, suggesting they may reflect lifetime-averaged environmental statistics, which are angularly isotropic—there are no globally preferred directions. This isotropy is broken within specific environments that animals explore over the short term, raising the question of whether head-direction systems show asymmetries reflecting this on timescales of minutes to hours. This would be an interesting experimental test. Additionally, while continuous-attractor models without continuous symmetry can theoretically be constructed, they may be more difficult to form through biologically plausible plasticity mechanisms. Existing models of ring-attractor self-organization [60] produce symmetric weights, and extensions to asymmetric cases, though they may exist, have not been presented to our knowledge.

### 7.2 From world structure to circuit structure

An intriguing question in neuroscience is: what can one infer about the structure of neural circuits from the structure of the world? Near sensory periphery, such inferences are feasible. Recognizing that sounds comprise superposed frequencies led Helmholtz to theorize that neural circuits decompose sounds into frequencies, as occurs in the cochlea [61]. Perceptual observations led Hering to propose color opponency, realized at the level of retinal ganglion cells and beyond [62]. In theoretical postdictions that could in principle have been predictions, Olshausen and Field [63] showed that natural image statistics generate V1-like receptive fields when represented by sparse, overcomplete neural populations. Several authors have noted that the structure discovered by Buck and Axel [64] of many receptors, each binding to multiple odorants resembles a compressed-sensing strategy for olfaction [65, 66].

Such inferences are more challenging deeper in the brain. However, consider the following argument that generalizes the approach taken in this paper. Begin with a continuous variable in the world, such as head direction, and assume that a fixed neural circuit maintains this variable by generating a persistent activity manifold. This manifold must have the same topology as the variable (e.g., a circle for head direction). What about its geometry? As argued above, environmental symmetries over an animal’s lifetime may lead the distribution of neuronal tuning curves to obey the symmetry of the variable (e.g., rotational symmetry). This requirement is much weaker than supposing all tuning curves have identical shapes up to shifts. At large *N*, this distributional symmetry becomes a geometric symmetry of the manifold. The recurrent weights must encode the manifold’s structure to generate it through neuronal dynamics. As we have shown, the geometric symmetry of the manifold appears in the weights as spectral degeneracies (doubly degenerate eigenvalues for a circle, multiples of six for a torus with hexagonal symmetry, and so on). Thus, on quite general grounds, we may expect symmetries of the world to be reflected in the spectral structure of neural-circuit connectivity, even far from sensory periphery.

This argument proceeds from topology to geometry to connectivity: the topology of a variable determines the topology of the neural manifold; environmental symmetries determine its geometry; and this geometry produces degeneracies in the weight-matrix spectrum. We propose that such spectral degeneracies are the best candidates for identifying continuous attractor structure in future mammalian connectome data. Generalizing eigenvalue-based analysis techniques to more complex circuits than considered here (e.g., with multiple cell types) is an important future direction.

### 7.3 Low-rank recurrent neural networks

The optimal weights (Eq. 3) exhibit an approximate low-rank structure, where the number of large eigenvalues is much smaller than the number of neurons *N*. This situates our model within the framework of low-rank recurrent neural networks, a well-studied class of models whose dynamics admit low-dimensional mean-field descriptions [35, 36, 67, 68]. In Sec. A1, we show that distributional symmetries in weight statistics generate corresponding symmetries in the effective low-dimensional dynamics (see also [69]). Our model instantiates this principle: circular symmetry in the connectivity distribution produces circular symmetry in the overlap dynamics. We further demonstrate that eigenvalue degeneracies, while highly suggestive of underlying geometric structure, do not uniquely determine the symmetry group of the dynamics without information about higher-order correlations in the weight statistics.

Previous work has proposed fitting low-rank network weights to neural data [67, 70], proceeding, via empirical optimization, from tasks to weights that can then be analyzed theoretically. In contrast, our approach provides an explicit analytical solution for the weights, enabling an end-to-end theory from data statistics through weight structure to mean-field dynamics. We achieve this analytical tractability by formulating continuous-attractor learning as a regression problem with a vanishing-flow constraint. The more challenging problem of analytically describing learning for inherently dynamical tasks remains largely open, though progress is being made [71].

### 7.4 Extension to general Lie groups

The heterogeneous-tuning network construction presented in this paper could be generalized to arbitrary compact Lie groups beyond the circle and torus considered here. The remaining (compact, non-toroidal) cases are non-abelian groups. As shown in Sec. A1, the resulting network models are expected to exhibit eigenvalue degeneracies inherited from the dimensions of the irreducible representations that comprise the group representation under which the connectivity transforms. These irreducible representations generalize the Fourier modes used in the toroidal cases. Exploring continuous-attractor models with heterogeneous tuning based on non-abelian symmetries represents an important avenue for future work.

### 7.5 From data to mean-field theories via generative processes

An interesting aspect of our analysis is the use of a statistical generative process to extend mean-field approaches to data-derived network models. The overall pipeline we have established—from characterizing single-neuron responses, to developing statistical models of neural heterogeneity, to deriving the necessary circuit mechanisms, to analyzing both the high-dimensional structure and mean-field limit—provides a general framework for understanding how ordered computation emerges within disordered neural circuits. It would be interesting to apply this pipeline to other brain regions and computations. For example, Gaussian-process modeling of neural trajectories has been performed in the motor system [72].

Several works prior to ours and Mainali et al. [29] have used nonlinearly transformed Gaussian processes as generative processes for neuronal tuning within unsupervised “manifold learning” approaches for neural data. Wu et al. [31] applied this approach to olfactory responses, learning a low-dimensional Euclidean olfactory space and neuronal tuning curves over this space. Jensen et al. [30] developed a more general framework for various manifolds, applying it to both fly and mouse head-direction neurons. While these methods share our use of Gaussian processes for modeling neuronal tuning, they differ fundamentally in approach. These methods simultaneously infer both latent-variable values and tuning curves by computing posterior estimates within the generative process. In contrast, we sample tuning curves from the prior distribution and assess whether their statistical features agree with tuning curves computed from spiking data using existing knowledge of the encoded variable. Additionally, these methods use maximum likelihood estimation to fit the full generative model, whereas we fit only the correlation function to match the data.

These methods could be powerfully coupled with our framework. Such coupling would enable application of our modeling and dynamical mean-field theory approach without prior knowledge of the encoded variable, creating an unsupervised version of the present work. It would also provide principled methods for fitting the parameters of the activation function and correlation function.

### 7.6 Order and disorder

The network models we have studied dynamically mimic classical continuous-attractor models while representing a much broader class of implementations. The ordered, translation-invariant architectures of classical models, corresponding to Fourier embeddings, represent only a special, measure-zero set of solutions within this broader space. This may explain why heterogeneous tuning is prevalent in larger brains such as those of mammals: it represents the generic case rather than a specialized solution. This raises the question: What conditions might favor the emergence of classical, translation-invariant architectures? One possibility is the constraint of limited neuronal resources. This could explain why flies, with their smaller nervous systems, implement an anatomical ring-attractor model that closely resembles classical models. The restricted number of available neurons may necessitate a more specialized, symmetrical solution to achieve reliable continuous-attractor dynamics [6]. As for structural perturbations, analysis of structural robustness shows that our disordered networks maintain, or fail to maintain, their attractor properties under weight perturbations comparably to classical circulant models (Sec. B5.1).

## Acknowledgments

We thank A. Litwin-Kumar, J. Zavatone-Veth, J. Cotler, and H. Grier for interesting conversations. This research was supported by the Gatsby Charitable Foundation (D.G.C., L.F.A. and H.S.) and the Kavli Foundation (D.G.C. and L.F.A.). H.S. was additionally supported by the Swartz Foundation and the Kempner Institute for the Study of Natural and Artificial Intelligence at Harvard University.

Anthropic’s Claude and OpenAI’s ChatGPT were used to edit the text and to generate code for both analyses and plotting.

## Appendix A: Theory

### A1 Symmetries, degeneracies, and dynamics in low-rank recurrent neural networks

The optimal weights (Eq. 3) exhibit approximate low-rank structure, where the number of large eigen-values is much smaller than the number of neurons *N* (indeed, the weights are exactly rank 2*D* + 1 if we truncate at *D* Fourier modes). This situates our model within the framework of low-rank recurrent neural networks, a well-studied class of models whose dynamics admit low-dimensional mean-field descriptions [35, 36, 67, 68]. This framework provides a natural setting for understanding: (1) how distributional symmetries in connectivity generate dynamical symmetries, (2) how these symmetries produce eigenvalue degeneracies, and (3) the nuanced relationship between eigenvalue degeneracies and dynamical symmetries.

In the general low-rank recurrent neural network framework, weight matrices take the form

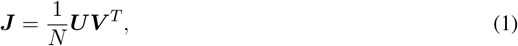

where ***U*** and ***V*** are *N × K* matrices with *K* fixed as *N → ∞*. Corresponding rows of ***U*** and ***V***, denoted by ***u*** and ***v***, are sampled from a joint distribution *P*(***u, v***) that is independent and identically distributed across rows. This construction yields network dynamics that can be analyzed through a *K*-dimensional vector of overlap order parameters,

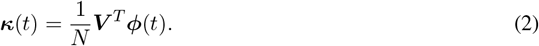

For low-pass filtered overlaps satisfying 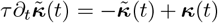, the effective dynamics as *N* → ∞ are

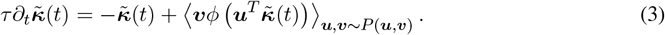

#### A1.1 Connectivity and dynamical symmetries

We now show that invariances in the weight matrix statistics generate corresponding invariances in the effective dynamics.

Suppose the joint distribution *P*(***u, v***) is invariant under transformation by a *K × K* orthogonal matrix ***R*** ∈ *O*(*K*):

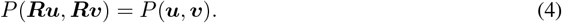

To see that this distributional symmetry translates to a symmetry of the overlap dynamics, use the invariance and change variables (***u***^*′*^, ***v***^*′*^) = (***Ru, Rv***) in the expectation:

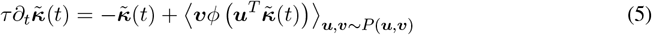

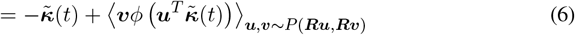

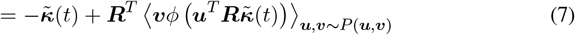

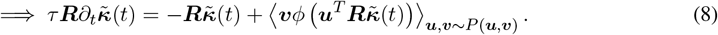

Thus, if 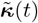 satisfies the effective dynamics Eq. 3, so does 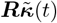. For example, if the system has a stable fixed point ***κ***^*^, then ***Rκ***^*^ is also a fixed point for any ***R*** satisfying the invariance condition. Note that the symmetry of the dynamics depends on the symmetry being obeyed by the full joint probability distribution of loading vectors.

A particularly interesting structure occurs when the set of orthogonal matrices ***R*** under which *P*(***u, v***) is invariant forms a continuous symmetry, specifically, a representation of some Lie group *G*. In this case, the dynamics admit the same continuous symmetry. For example, if the system has a stable fixed point ***κ***^*^, then (generically) its entire orbit {***Rκ***^*^|***R*** ∈ *G*} forms a continuous attractor manifold with the same structure as the group *G*.

A recent preprint of Pezon et al. [69] established a related result within a more specialized framework where neurons have assigned coordinates in a low-dimensional feature space and weight-matrix elements measure feature similarity through a kernel. Our analysis operates directly at the level of loading-vector statistics. The following eigenvalue and group-based analyses are, to our knowledge, novel.

#### A1.2 Eigenvalue degeneracies from continuous symmetries

We now show how continuous symmetries of the weight statistics lead to eigenvalue degeneracy, and moreover show that eigenvalue degeneracy does not uniquely determine the symmetry of the dynamics.

The eigenvalues of the *N × N* weight matrix are the same as the eigenvalues of the *K × K* overlap matrix 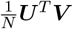, which as *N* → ∞ becomes

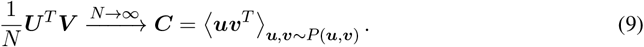

The invariance of *P*(***u, v***) under ***R*** implies, by changing variables (***u***^*′*^, ***v***^*′*^) = (***Ru, Rv***) in the expectation,

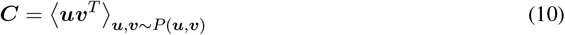

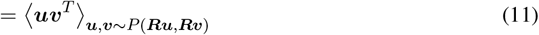

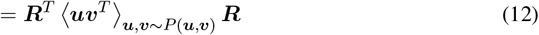

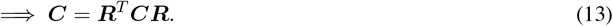

That is, ***C*** commutes with ***R***.

Now suppose that ***R*** is an orthogonal representation of a compact Lie group *G* (compactness is needed to ensure complete reducibility of the representation in what follows). Its action on ℝ^*K*^ can be decomposed into invariant subspaces in which it acts with an irreducible representation. In the following statement we assume for simplicity that each irreducible representation is non-repeated, but this is not essential. Labeling the irreducible representations by dimensions *d*_1_, …, *d*_*L*_, the relation ***C*** = ***R***^*T*^ ***CR*** implies that the matrix ***C*** (and hence ***J***) has eigenvalue degeneracies, with each irreducible representation *ℓ* contributing a set of *d*_*ℓ*_ degenerate eigenvalues. Thus, the eigenvalue degeneracies are determined by the dimensions of the irreducible representations that constitute the representation of the Lie group under which the connectivity transforms.

Note that the eigenvalue degeneracies, and eigenvalues in general, depend only on the second-order statistics of the loading vectors.

#### A1.3 Application to our model

The ring-attractor model studied in this paper instantiates the above principles. At large *N*, the optimized weights of our model (Setting 3) take the form 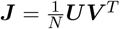 where

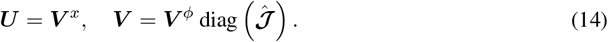

Here, ***V*** ^*x*^ and ***V*** ^*ϕ*^ contain Fourier coefficients, scaled to have unit variance, of input-current and firing-rate tuning curves, respectively, expressed in the sine-cosine basis (constant mode followed by (cos(*kθ*), sin(*kθ*)) pairs for *k* = 1, …, *D*). The vector 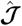 contains (2*D* + 1) Fourier coefficients (Eq. 17) that exhibit doublet degeneracy.

The pairwise covariances ⟨***uu***^*T*^⟩, ⟨***vv***^*T*^⟩, and ⟨***uv***^*T*^⟩ are all diagonal. Elements of ***u*** are marginally Gaussian due to the Gaussian process construction. In contrast, ***v*** exhibits higher-order correlations across modes and with modes of ***u*** due to the nonlinear transformation *ϕ*(*·*) applied in *θ*-space, making the joint distribution *P*(***u, v***) non-Gaussian with complex higher-order structure. Despite this complexity, the distribution possesses an underlying circular *SO*(2) symmetry inherited from the rotation-invariant input-current Gaussian process.

To define this symmetry concretely, let ***R***(*ψ*) ∈ *O*(2*D* + 1) be the (2*D* + 1) *×* (2*D* + 1) matrix

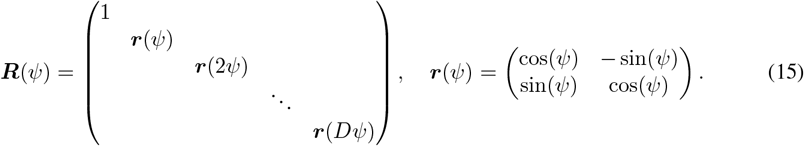

For each angle *ψ*, the transformation ***R***(*ψ*) leaves the distribution invariant:

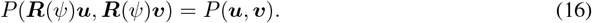

This invariance reflects the fact that rotating the tuning curves *x*^⋆^(*θ*) and *ϕ*^⋆^(*θ*) by an angle *ψ* does not change the likelihood of observing them. Note that it is crucial that the same *ψ* appears in each block for this to hold.

The matrices ***R***(*ψ*) form a representation of *SO*(2) acting on the (2*D* + 1)-dimensional space of Fourier coefficients. This representation is reducible and decomposes into a direct sum of *D* + 1 irreducible representations: one trivial (constant mode) and *D* two-dimensional representations. Each two-dimensional irreducible representation contributes a degenerate pair of eigenvalues to the spectrum of ***J***. Importantly, the presence of multiple irreducible representations beyond the first one is precisely what generates heterogeneity in tuning curves, beyond just amplitude variability as one would get with a single irreducible representation. By the principle above, this distributional symmetry generates *SO*(2) symmetry in the overlap dynamics.

Note that the dynamical mean-field theory in Sec. 5 describes the same dynamics in *θ*-space 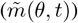, enabling direct comparison to classical models, while the present analysis is in Fourier-coefficient space 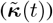.

#### A1.4 Do eigenvalue degeneracies imply dynamical symmetries?

Eigenvalue degeneracies, while highly suggestive of underlying symmetry, do not strictly determine the symmetry group of the dynamics. This is because eigenvalue degeneracies depend only on the second-order statistics of the loading vectors, whereas the symmetry group of the dynamics is determined by the full distribution *P*(***u, v***).

To illustrate this, consider a modification of our model where the joint distribution *P*(***u, v***) is Gaussian with the same second-order moments as in our actual model. This Gaussianization preserves the weight-matrix eigenvalues and in particular their doublet degeneracy. However, the symmetry group of the dynamics expands to have the product structure (*SO*(2))^*D*^ rather than *SO*(2). This enlarged group allows independent rotations of each doublet:

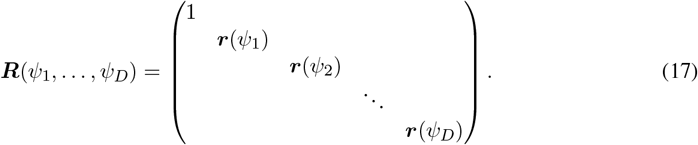

**Figure A1:**
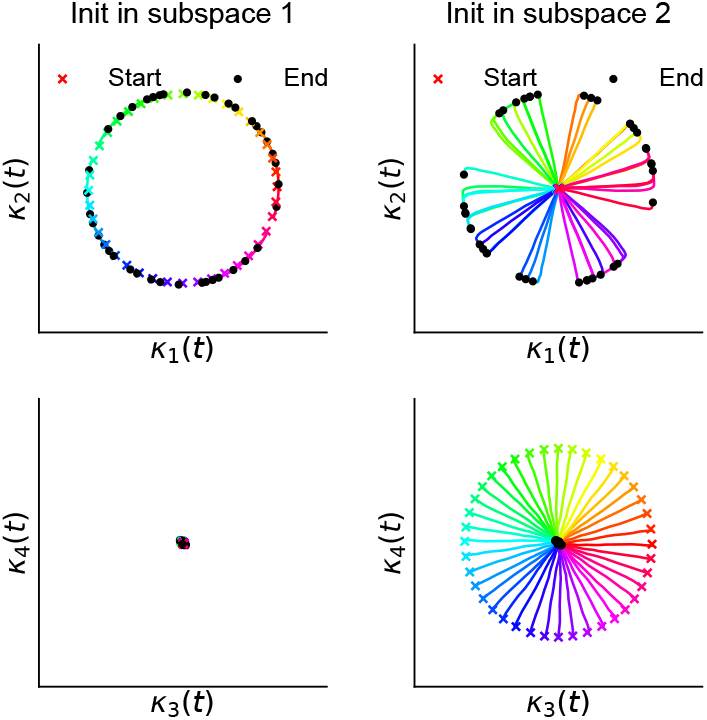
Rank-four Gaussian network with (*SO*(2))^2^ symmetry exhibiting winner-take-all behavior (*N* = 1000, *K* = 4). Colored lines show evolution of the four components of ***κ***(*t*) over 10 s. Crosses indicate initial states, dots indicate final states. **Left:** Initialization in dominant subspace. **Right:** Initialization in subdominant subspace. In both cases, activity converges to the dominant subspace.

Under this Gaussian modification, the fixed-point structure is a *D*-dimensional torus rather than a one-dimensional ring. In our model, the higher-order correlations of *P*(***u, v***) arising from the nonlinear transformation *ϕ*(*·*) constrain the symmetry group of the dynamics to *SO*(2) rather than (*SO*(2))^*D*^.

There is an interesting caveat, however: in the Gaussian construction described here, while the dynamics possess the product-group symmetry (*SO*(2))^*D*^, only a single two-dimensional subspace corresponding to the highest-variance doublet is stable, and the projections onto the remaining two-dimensional subspaces decay to zero. To demonstrate this, we constructed a rank-four network (*K* = 4) where, for simplicity, corresponding columns of ***U*** and ***V*** are perfectly correlated (i.e., ***U*** = ***V***), with the first two columns having variance 10 and the second two having variance 5. The overlap vector ***κ***(*t*) is four-dimensional. We initialized the system along a ring in either the dominant subspace (first doublet) or the subdominant subspace (second doublet). In both cases, only the components of ***κ***(*t*) corresponding to the dominant subspace survive asymptotically, and we observe quasi-continuous-attractor behavior (Fig. A1). This winner-take-all behavior is analogous to that seen in networks with Gaussian loadings storing discrete fixed points [36]; our construction represents a generalization of this result to continuous manifolds.

To conclude: While eigenvalue degeneracies are highly non-generic features and thus serve as suggestive indicators of potential dynamical symmetry, they do not uniquely specify the symmetry group of the dynamics without additional information about higher-order structure. Although the eigenvectors ***V*** ^*x*^ and ***V*** ^*ϕ*^ appear disordered in scatterplots (Fig. 7F, Fig. B6), and thus naive techniques such as spectral embedding do not work, they contain higher-order correlations that encode the *SO*(2) symmetry. Whether this structure can be reliably extracted from data remains an open question. Eigenvalue degeneracies appear to be the most robust and readily identifiable signature of continuous-attractor organization in connectivity data.

### A2 Computing the weight matrix

We use the same method to construct the weight matrix ***J*** in Setting 2 and Setting 3.

#### A2.1 Problem setup

We seek to find ***J*** that minimizes the loss function of Eq. 2 subject to the constraint *J*_*ii*_ = 0. In this section, we omit the superscripts ⋆ from 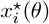 and 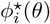 since there are no dynamic variables from which we must disambiguate them. As in the analysis of Sec. 4, we define **Φ** as an *N*_*θ*_ *× N* matrix whose columns contain the tuning curves *ϕ*_*i*_(*θ*), and ***X*** as an *N*_*θ*_ *× N* matrix containing the corresponding input currents *x*_*i*_(*θ*). Specifically, *X*_*ai*_ = *x*_*i*_(*θ*_*a*_) and Φ_*aj*_ = *ϕ*_*j*_(*θ*_*a*_), where we sample *θ* uniformly on [0, 2*π*) at 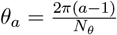 for *a* = 1, …, *N*_*θ*_. In the data-derived network case, *N* = *N*_data_.

#### A2.2 Discrete-continuous correspondence

The general dictionary for converting between continuous and discrete representations is

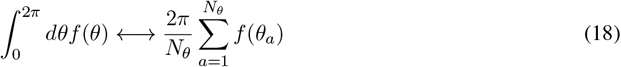

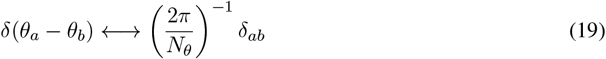

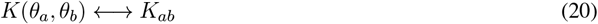

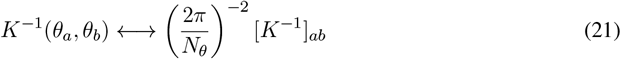

The delta function and inverse formulas follow from the requirements that 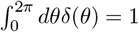 and 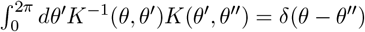, respectively.

#### A2.3 Loss function and unconstrained solution

The loss function (reproduced from Eq. 2) is

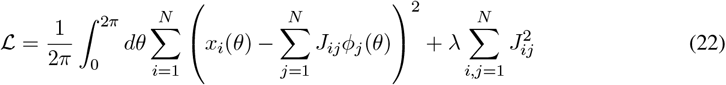

We first solve the **unconstrained** optimization problem. Differentiating with respect to *J*_*ij*_ gives:

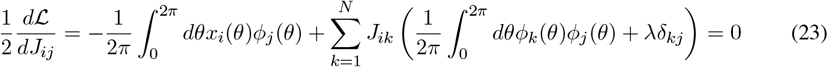

Converting to the discretized setting gives

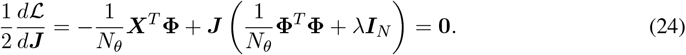

Solving this system yields

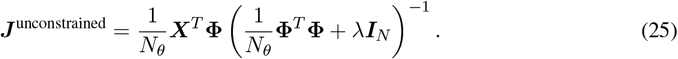

Using the identity

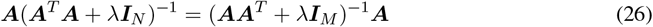

for a *M × N* matrix ***A***, this can be rewritten as

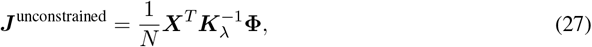

where the kernel matrix is

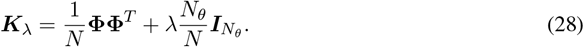

In the continuous limit, this becomes

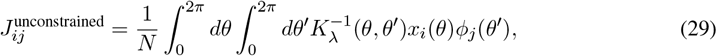

where the kernel matrix is

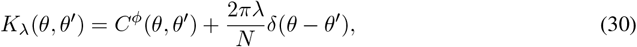

and 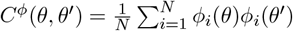 is the empirical correlation function of the tuning curves (Eq. 4).

#### A2.4 Constrained solution

We now enforce the constraint *J*_*ii*_ = 0 (the **constrained** solution). Note that the optimization problem decouples across rows of ***J***, where each row predicts *x*_*i*_(*θ*) from regressors *ϕ*_*j*_(*θ*) for *j* = 1, …, *N*. Constraining *J*_*ii*_ = 0 means excluding regressor *ϕ*_*i*_(*θ*) and adjusting the weights from the other regressors accordingly. This gives, upon modifying Eq. 27 and Eq. 28:

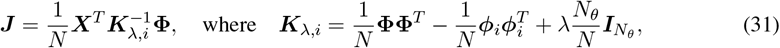

where ***ϕ***_*i*_ is the *N*_*θ*_-dimensional vector containing *ϕ*_*i*_(*θ*). Taking the continuous limit yields Eq. 3 of the main text, reproduced here:

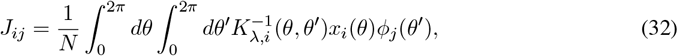

where the kernel matrix is

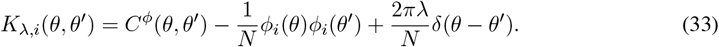

Note that this formula for *J*_*ij*_ applies only for *i* ≠ *j*; for *i* = *j, J*_*ii*_ = 0. This formula for the kernel makes it clear that the constraint of zero on-diagonals provides a small, 𝒪(1*/N*) correction to the off-diagonals relative to the unconstrained solution. Thus, at large *N*, it may be omitted.

Unfortunately, the above formula requires inverting a separate matrix for each neuron *i*, which is computationally expensive. This can be circumvented using the Sherman-Morrison formula (rank-one update to a matrix inverse). Here, we instead use Lagrange multipliers to obtain the same result. The modified loss function becomes

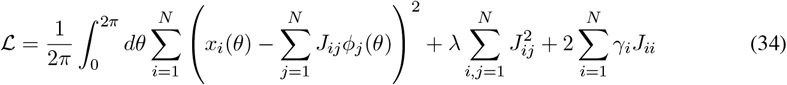

where *γ*_*i*_ are Lagrange multipliers for *i* = 1, …, *N*. Differentiating with respect to *J*_*ij*_ gives

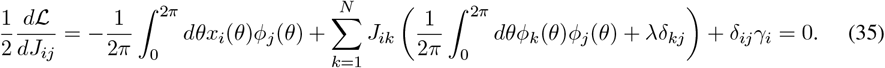

In discretized form,

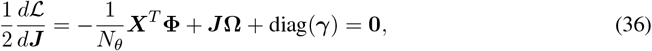

where we have defined the *N × N* matrix **Ω**,

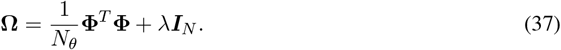

In the continuous limit,

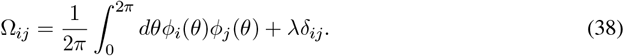

Solving for ***J*** yields

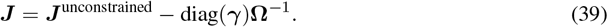

The Lagrange multipliers are determined by the constraint *J*_*ii*_ = 0, yielding

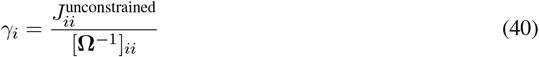

Therefore, the final constrained solution is

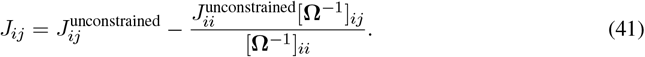

This is the formula we use in practice.

### A3 Details of large-N analysis of optimized weights

#### A3.1 Fourier conventions in Sec. 4

The eigenvalues of a circulant kernel matrix—for example, Γ^*x*^(*θ* − *θ*^*′*^), where *θ* and *θ*^*′*^ serve as the indices— are given by the Fourier transform of Γ^*x*^(Δ*θ*) in the complex exponential basis. These eigenvalues have doublet degeneracy, 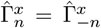, where *n* = −*D*, …, *D*. Within each two-dimensional degenerate subspace, we are free to choose any eigenbasis. In Sec. 4 of the main text (whose details are given below) and Sec. A1 of the Theory Appendix, we adopt a sine-cosine basis, which is a rotation of the complex-exponential basis, because we require real singular vectors for the analysis.

Note that, when using this basis, we do *not* use the sine-cosine Fourier coefficients of Γ^*x*^(Δ*θ*). These coefficients possess only nonzero cosine terms and thus are not the correct eigenvalues of the kernel matrix, which come from the complex-exponential Fourier transform.

#### A3.2 Weight-matrix analysis details

For finite *N*, the solution to the optimization problem can be expressed as follows. Recall that ***X*** and **Φ** are *N*_*θ*_ *× N* matrices containing the input-current and firing-rate tuning curves. We start from Eq. 27 and Eq. 28 and define *λ*_*θ*_ = *N*_*θ*_*λ*. We have

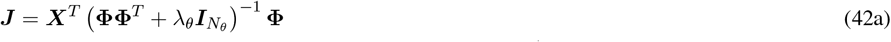

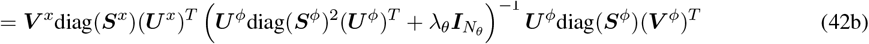

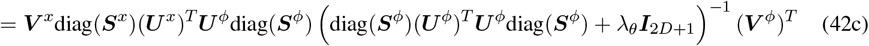

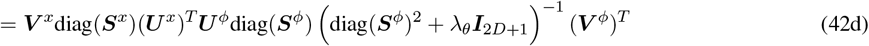

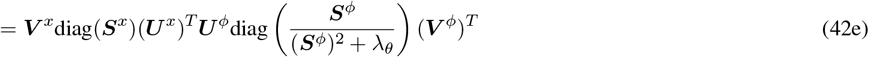

The steps applied are: (a→b) substitution of SVDs; (b→c) using the identity Eq. 26; (c→d) using orthonormality of ***U*** ^*ϕ*^ to simplify (***U*** ^*ϕ*^)^*T*^ ***U*** ^*ϕ*^ = ***I***_2*D*+1_; and (d→e) is cosmetic.

Taking *N* → ∞ and using the convergence of left singular vectors to Fourier modes (***U*** ^*x*^ → ***F***, ***U*** ^*ϕ*^ → ***F***) and singular values to Fourier coefficients 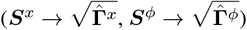, we obtain:

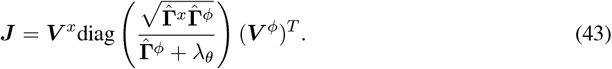

Taking the *λ*_*θ*_ → 0 (minimum-norm) solution yields:

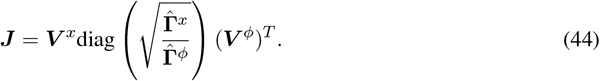

To establish the right singular vector overlaps, we rearrange the SVDs (for any *N*) to obtain

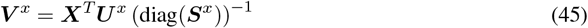

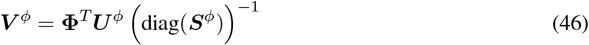

Therefore:

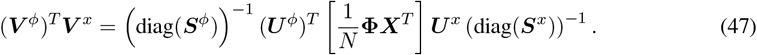

As *N* → ∞, the empirical *ϕ*-*x* correlation function (the bracketed term) converges to a limiting *N*_*θ*_ *× N*_*θ*_ matrix whose (*θ, θ*^*′*^) element is

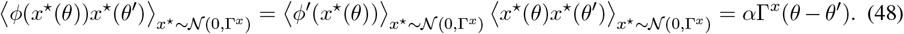

Here we applied Price’s theorem, which is valid since *x*(*θ*) is Gaussian, and 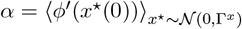 is the average slope of the activation function with respect to the Gaussian distribution of input currents (Eq. 16). Since the ***U*** matrices converge to Fourier modes at large *N* (***U*** ^*x*^ → ***F***, ***U*** ^*ϕ*^ → ***F***), we have

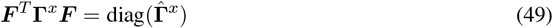

Combining these results gives

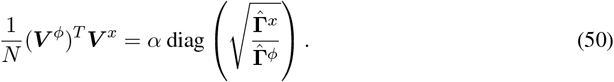

This demonstrates that, at large *N*, non-corresponding columns of ***V*** ^*x*^ and ***V*** ^*ϕ*^ are orthogonal, while corresponding columns are aligned, but not perfectly aligned. Using Eq. 44 and Eq. 50, one can verify that columns of ***V*** ^*x*^ are right eigenvectors of ***J*** and columns of ***V*** ^*ϕ*^ are left eigenvectors of ***J***, with eigenvalues

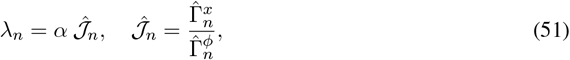

as given in the main text.

#### A3.3 Attempting to extract information about the latent ring structure from the eigen-vectors despite their disordered nature

One can attempt to extract information about the latent ring structure from the eigenvectors despite their disordered nature. We took the top two columns of ***V*** ^*x*^ and defined angular coordinates using the *N* two-dimensional embedding coordinates [51]. Despite the independent and identically distributed nature of these coordinates (which form an isotropic Gaussian cloud, as expected), this angular ordering reveals the noisy circulant structure, shown in the data-derived network case in Fig. 3B.

One can ask whether this “sort-the-neurons-and-check-for-circulant-structure” method is viable for detecting latent ring structure in connectomes. To test this, we constructed null-model matrices that lack doublet degeneracy in their singular-value spectra and have variable approximate rank (Fig. A2; details of analysis in figure caption). We find that being approximately low-rank is sufficient for revealing noisy circulant structure upon sorting the entries of these null-model matrices; doublet degeneracy (i.e., an actual latent ring structure) is not necessary. This demonstrates that the method produces false positive detections of circular symmetry, hallucinating circulant patterns from low-rank structure alone.

**Figure A2:**
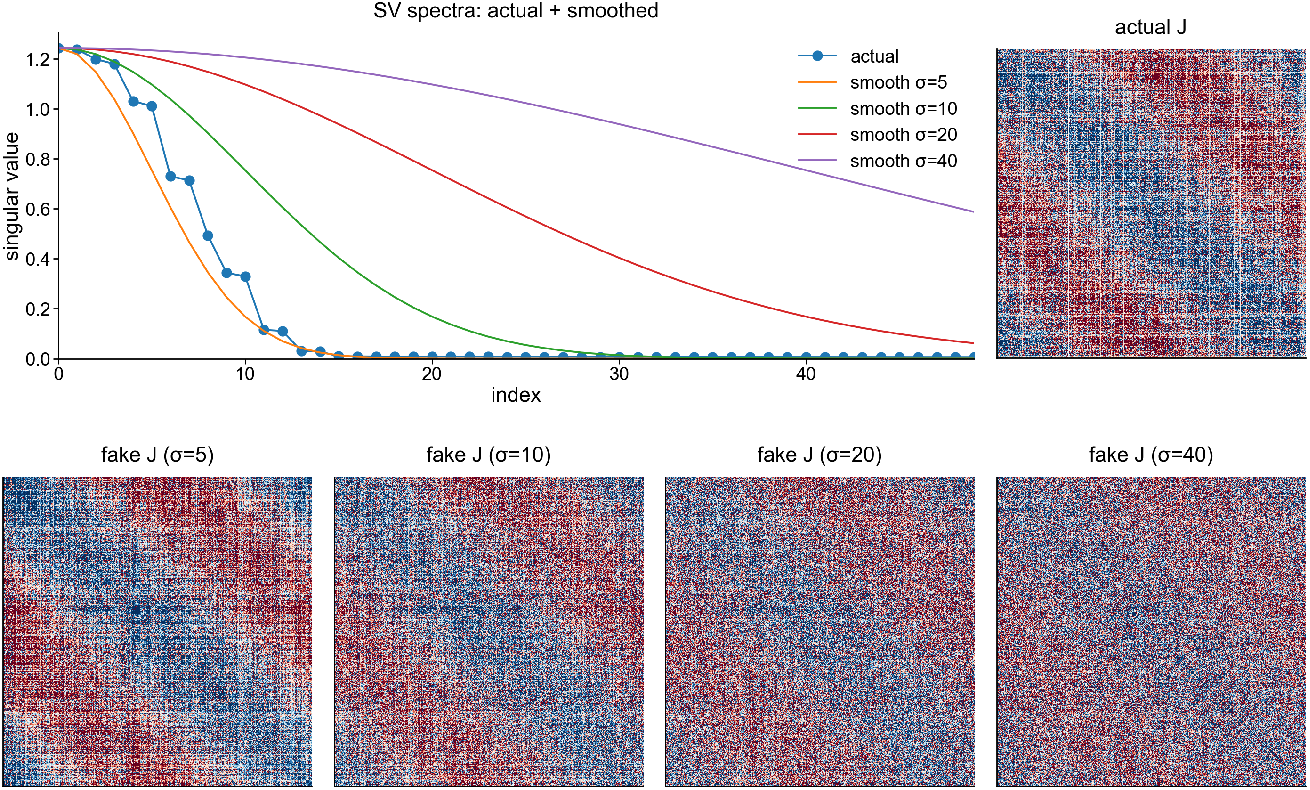
Spectral embedding hallucinates circulant structure from low-rank matrices without circular symmetry (*N* = 2500, generative-process Fourier decay scale *σ* = 1.42, response sharpness *β* = 2.76). **Top left:** Singular value spectra of the optimal weight matrix (exhibiting doublet degeneracy) and four null models with smoothed singular value spectra. Null models have singular values obtained by applying Gaussian smoothing with scales *σ*_smooth_ ∈ {5, 10, 15, 20} to the original spectrum, eliminating degeneracy and varying the approximate rank. These matrices are constructed as ***J*** ^null^ = ***O***diag(***S***)***O***^*T*^, where ***O*** is a Haar random orthogonal *N × N* matrix. **Top right:** Optimal weight matrix ***J*** with neurons sorted by angular coordinates derived from the top two left singular vectors, revealing circulant structure. **Bottom row:** Four null model matrices sorted using identical procedures, showing hallucinated circulant structure that arises purely from low-rank organization rather than circular symmetry. The false positive detection is strongest for small *σ*_smooth_ (highly low-rank) and diminishes as *σ*_smooth_ increases.

### A4 Details of dynamical mean-field theory

#### A4.1 Derivation of dynamical mean-field theory equations

We begin by deriving the dynamical mean-field equations for zero velocity input (*ω*(*t*) = 0), which capture the essential features of the derivation while simplifying the presentation. This allows us to describe the transient evolution of the overlap order parameter from arbitrary initial conditions. We then extend the analysis to incorporate velocity inputs. The problem whose solution we derive here is a special case of the more general problem analyzed in Supplement: Dynamics of learned attractors using a path-integral approach.

Starting with the network dynamics (Eq. 1) with *b* = 0 and the solution for the weights *J*_*ij*_ (Eq. 3), we have

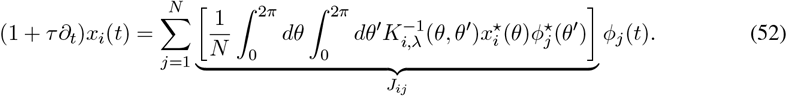

In the limit *N* → ∞ with *λ* = 0, the inverse kernel 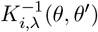 converges to [Γ^*ϕ*^]^*−*1^(*θ* − *θ*^*′*^), where Γ^*ϕ*^(Δ*θ*) is invertible due to all Fourier components being nonzero. Moving the sum over *j* inside the integrals and using the definition of *m*(*θ, t*) from Eq. 6, we obtain

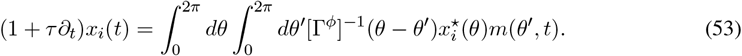

We introduce a temporally smoothed version of the overlap order parameter,

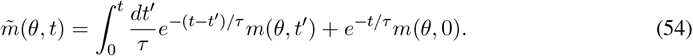

This satisfies

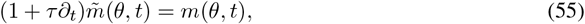

as stated in the main text. Applying (1 + *τ*∂_*t*_)^*−*1^ to both sides of the dynamics equation (Eq. 53) yields

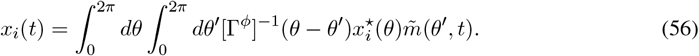

Applying *ϕ*(*·*) to both sides of this equation, and then taking the inner products of both sides with 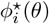 and dividing them by *N*, gives

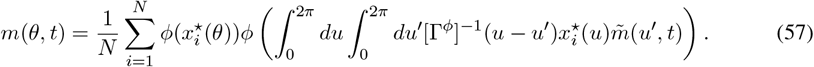

The key step in deriving the mean-field equations is to replace the empirical average over neurons (*i* index) with an expectation over the generating Gaussian process. That is,

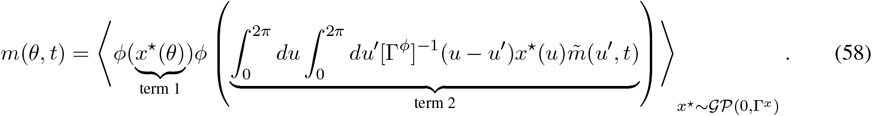

This replacement is exact for *N → ∞*. Terms 1 and 2 are jointly Gaussian with zero mean. Their variances and covariance fully determine the right-hand side of the equation and are given by

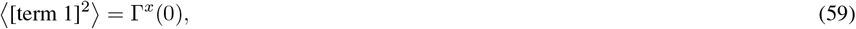

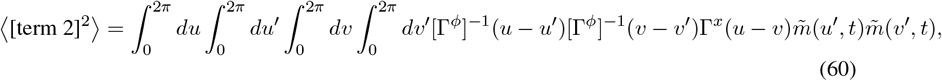

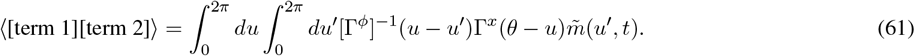

We define kernels in Fourier space:

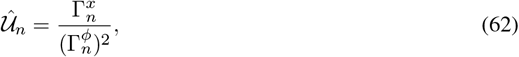

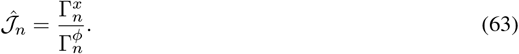

The second equation (Eq. 63) defines the effective rotation-invariant weights stated in the main text (Eq. 17). These kernels allow us to express the correlations between terms 1 and 2 as

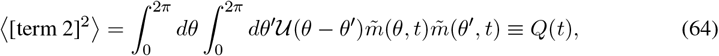

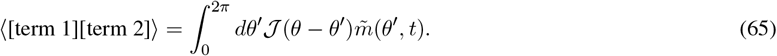

Now, let 𝒢(*C*_11_, *C*_22_, *C*_12_) denote the expectation

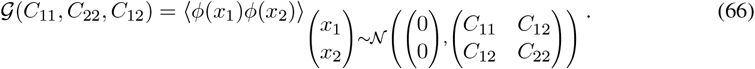

This can be explicitly evaluated using the activation function 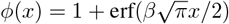, which yields

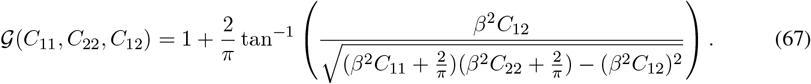

Then, Eq. 58 can be written as

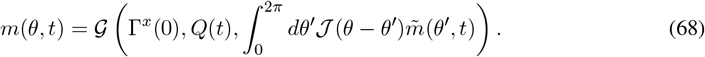

Defining

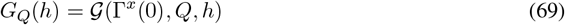

and using the relationship between *m*(*θ, t*) and 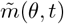, we obtain

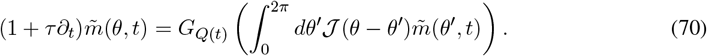

These are the mean-field dynamics from the main text (Eq. 18) with *ω*(*t*) = 0.

#### A4.2 Velocity-dependent dynamics

We now extend this derivation to include nonzero angular velocity *ω*(*t*). The network dynamics are modified to include an angular velocity-dependent term,

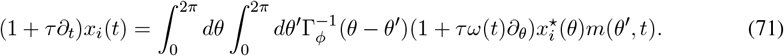

To solve Eq. 71, we again apply (1 + *τ*∂_*t*_)^*−*1^ to both sides, and now introduce a variable 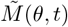 defined as

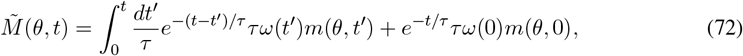

which satisfies

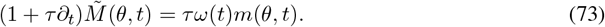

Then, we have

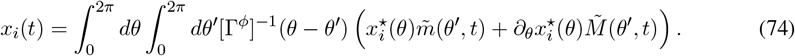

We now massage Eq. 72 by expanding *ω*(*t*^*′*^) in a Taylor series around *t* (dropping the initial transient for simplicity),

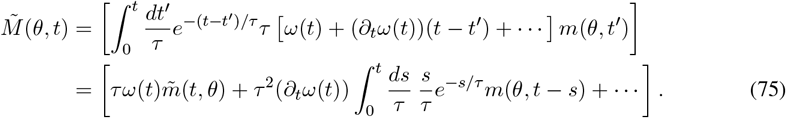

The expansion in square brackets is in powers (*τ*∂_*t*_)^*n*^*τω*(*t*) for *n* = 0, 1, 2, …, which we will assume can be truncated after *n* = 0. Note that this assumption means that the angular acceleration, given by ∂_*t*_*ω*(*t*) (times appropriate factors of *τ*), is small, but the angular velocity itself may be large. This gives

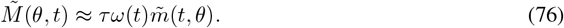

Substituting Eq. 76 back into Eq. 74, we obtain

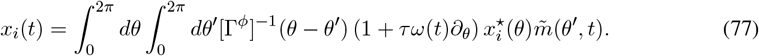

Following the same averaging procedure as in the zero-velocity case, but now accounting for the velocity-dependent terms in Eq. 77, the correlations are modified in two ways:

1. The kernel 𝒰(Δ*θ*) that determines the activity-dependent gain parameter *Q*(*t*) (the variance of input currents) is modified to include an angular velocity-dependent term,

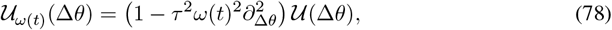

where 𝒰(Δ*θ*) is the original kernel defined through its Fourier transform 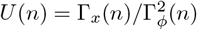.
2. The correlation between input currents and target tuning curves acquires an additional angular velocity-dependent term,

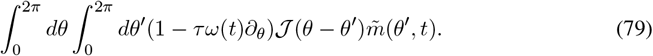

These modifications lead to the full dynamical mean-field equation for the overlap order parameter,

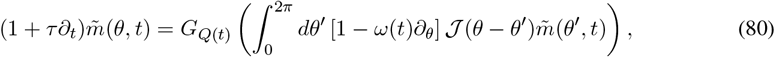

where *Q*(*t*) uses 𝒰_*ω*(*t*)_(Δ*θ*) (Eq. 78) in the quadratic term.

#### A4.3 Jacobian analysis of the mean-field dynamics

To analyze the stability of solutions to the mean-field equations, we need to compute the Jacobian matrix *M*(*θ, θ*^*′*^), which describes how perturbations to the overlap order parameter evolve in time. This requires expressions for derivatives of the effective activation function *G*_*Q*_(*h*) with respect to both *h* and *Q*. Recall that *G*_*Q*_(*h*) is defined through the function 𝒢(*C*_11_, *C*_22_, *C*_12_), which computes the expectation in Eq. 66.

We define auxiliary quantities

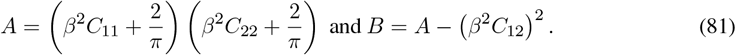

Then, the required derivatives are

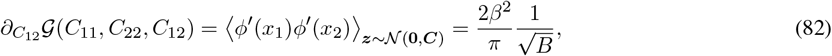

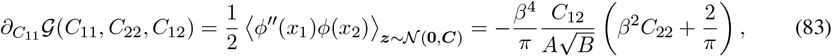

where ***C*** = ((*C*_11_, *C*_12_), (*C*_12_, *C*_22_)).

For the bump solution 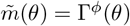, the Jacobian takes the form

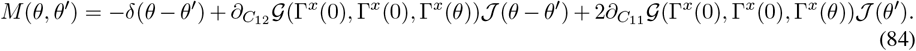

Each of the three terms represents a distinct contribution to the dynamics: exponential decay (first term), direct coupling through the effective weights (second term), and modulation through the activity-dependent gain *Q*(*t*) (third term).

For the homogeneous solution 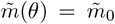, we first determine 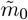 by numerically solving the self-consistent equations

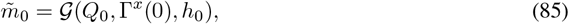

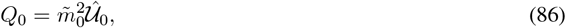

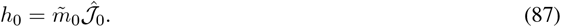

The Jacobian at this solution is then

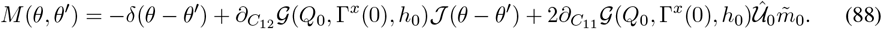

Finally, we explain how to analyze the dynamical mean-field theory equations in scenarios when the dynamics of *Q*(*t*) are excluded by fixing *Q* = Γ^*x*^(0), as discussed in the main text. We simply omit the terms involving 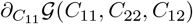 in both Jacobians (and, if we would like to find the new stable uniform state, denoted by 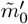 in the main text, we fix *Q*_0_ = Γ^*x*^(0) in the self-consistent equations above rather than allowing it to depend on 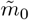). The presence of a positive eigenvalue of the Jacobian for the bump state when we fix *Q* = Γ^*x*^(0), shown in Fig. 9F, demonstrates that the activity-dependent gain modulation through *Q*(*t*) is essential for bump stability.

#### A4.4 Finite domain of effective activation function

The effective activation function *G*_*Q*_(*h*) (Eq. 69) is defined only for

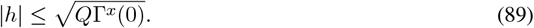

Beyond this range, the 2 × 2 covariance matrix of the underlying bivariate Gaussian is non-positive-definite.

This bound holds in the dynamical mean-field system, as we now show. Using matrix-vector notation for clarity, we have:

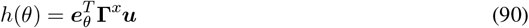

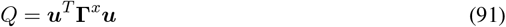

where ***u*** = (**Γ**^*ϕ*^)^*−*1^***m***(*t*) and ***e***_*θ*_ selects angular component *θ*. Since **Γ**^*x*^ is positive definite, the Cauchy-Schwarz inequality implies that

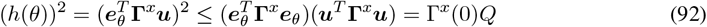

#### A4.5 Reciprocal symmetry of J at large N

It is straightforward to show that

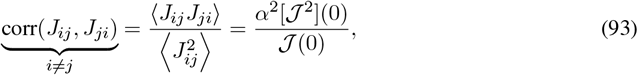

where

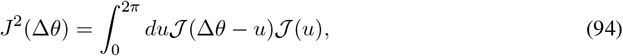

where 𝒥(Δ*θ*) are the circulant interactions in the dynamical mean-field theory, and *α* is given by Eq. 16.

#### A4.6 Emergence of Mexican-hat interactions from tuning-curve statistics

The Mexican-hat profile emerges from the neuronal activation function’s effect on correlation structure, specifically, from the generation of a certain quartic term in the Fourier transform of the effective weights.

To illustrate this mechanism, consider correlation functions on one-dimensional infinite position space using coordinate *x* and frequency *k* rather than *θ* and *n* (circularity is not essential). Define the input-current correlation function as Γ^*x*^(Δ*x*) = exp (−Δ*x*^2^*/*2*ℓ*^2^), corresponding to the Gaussian form used in our generative process. The firing-rate correlation function becomes Γ^*ϕ*^(Δ*x*) = *G*(1, 1, Γ^*x*^(Δ*x*)) through the activation function. We take the activation function for simplicity to be 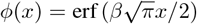.

**Figure A3:**
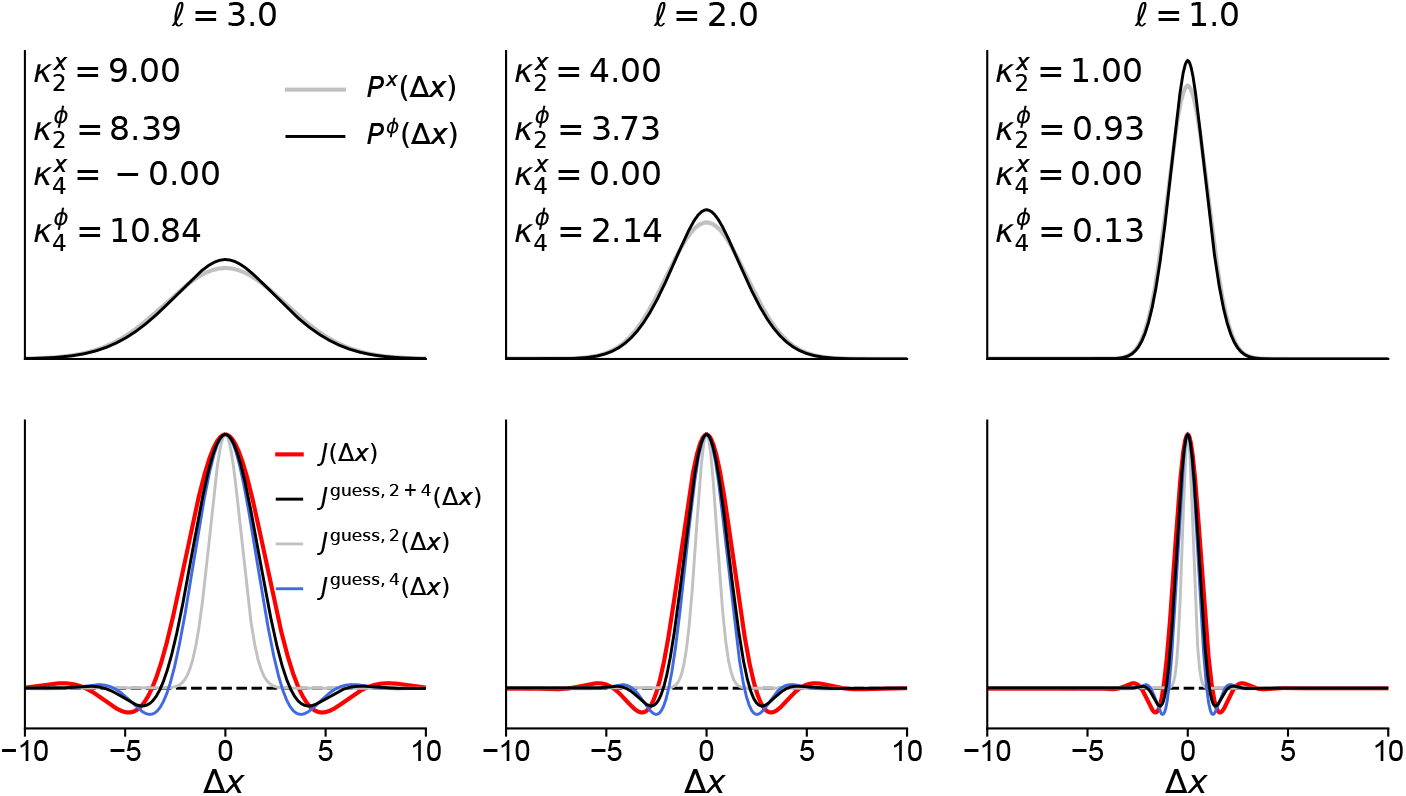
Origin of Mexican-hat connectivity profile. As defined in the main text, Γ^*x*^(Δ*x*) = exp (− Δ*x*^2^*/*2*ℓ*^2^) and Γ^*ϕ*^(Δ*x*) = 𝒢(1, 1, Γ^*x*^(Δ*x*)) with activation function 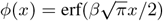. Columns correspond to length scales *ℓ* = 3, 2, 1 (left to right). **Top row:** Input-current correlation *P*^*x*^(Δ*x*) and firing-rate correlation *P*^*ϕ*^(Δ*x*) (both normalized to probability distributions) with their second and fourth cumulants displayed. **Bottom row:** Comparison of the full effective weights 𝒥(Δ*x*), computed from Fourier-coefficient ratios, with various approximations: 𝒥_approx,2+4_ includes both quadratic and quartic terms; 𝒥^approx,2^ excludes the quartic term; and 𝒥^approx,4^ excludes the quadratic term. The quartic term is essential for generating the Mexican-hat structure. Including the quadratic term further improves approximation accuracy.

Normalizing these to probability distributions 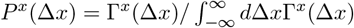 and similarly for *P* ^*ϕ*^(Δ*x*), we obtain cumulants 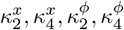. Due to the Gaussian form of Γ^*x*^(Δ*x*) (a distinct notion from the Gaussian process over the input currents), we have 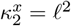 and 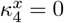. What happens to 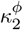 and 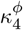 is nontrivial.

The effective weights in Fourier space are:

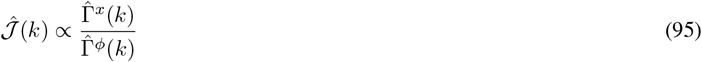

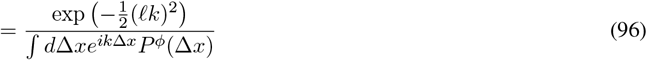

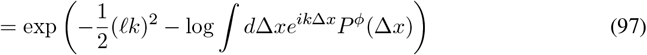

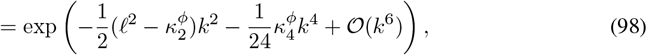

where in the last line we recognized the cumulant generating function of *P* ^*ϕ*^(Δ*x*).

The neuronal activation function “tightens” tuning, a well known effect leading to 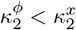. However, a less considered aspect is what happens to the fourth cumulant 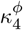. We find that the neuronal activation function leas to positive fourth cumulant, 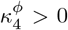. This quartic term in Fourier space is essential for the Mexican-hat profile. If it were zero, 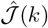 would be purely Gaussian, producing a Gaussian (not Mexican-hat) profile in position space. Indeed, as demonstrated in Fig. A3, the observed quadratic and quartic terms are sufficient for generating a Mexican-hat shape.

### A5 Existence of continuous attractors with arbitrary manifolds via universal approximation

Consider dynamics of a *K*-dimensional vector ***κ***(*t*) governed by

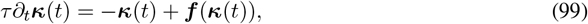

where ***f*** : ℝ^*K*^ → ℝ^*K*^ is a continuous evolution function. The decay term −***κ***(*t*) imposes no loss of generality. These dynamics may possess a continuous attractor manifold of dimension ≤ *K*.

Universal approximation (see, e.g., Chong [73]) guarantees that, for sufficiently large *N*, one can find *N × K* matrices ***U*** and ***V*** such that

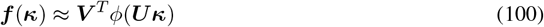

to arbitrary accuracy uniformly over ***κ*** restricted to a compact set (e.g., encompassing the explored state space of the dynamics), provided that *ϕ*(*·*) is non-polynomial. Defining ***x***(*t*) = ***Uκ***(*t*) yields

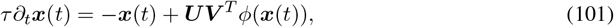

which is a rank-*K* recurrent neural network embedding the original dynamical system in a *K*-dimensional subspace spanned by the columns of ***U***.

One might worry that choosing ***U*** to implement the approximation distorts the geometry of the ***κ***(*t*) dynamics through a non-orthonormal embedding. However, the approximation property holds with probability one even when elements of ***U*** are chosen randomly [73], in which case the columns of ***U*** become orthogonal at large *N*.

The construction used here of a low-rank recurrent neural network that emulates a given dynamical system by representing the evolution function via a one-hidden layer neural network is essentially the same as that proposed by the Neural Engineering Framework [74]. The random-***U*** construction suggested above would result in a ring-attractor model with a random embedding.

### A6 Fourier transforms

#### A6.1 Conventions and facts

Here, we consolidate various conventions and facts about Fourier transforms. Let 𝕋^*d*^ = (𝕊^1^)^*d*^ denote the *d*-dimensional torus. We present the one-dimensional case (the circle, 𝕊^1^) which extends easily to higher dimensions.

For a function *F*(*θ*) with *θ* ∈ 𝕊^1^, we define the Fourier coefficients and their inversion by

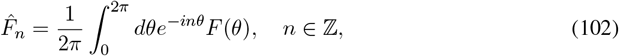

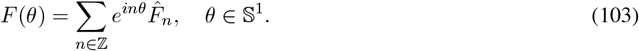

The forward transform maps a function of the continuous circular variable *θ* to a discrete set of coefficients indexed by *n*, while the inverse transform reverses this mapping. These relations are inverses of one another due to the orthogonality identities

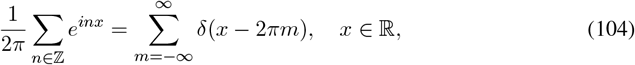

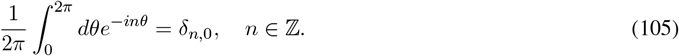

The sum of delta functions in Eq. 104 is called a Dirac comb.

For functions *f*(*x*) on the real line (*x* ∈ ℝ), we define the Fourier transform and its inverse by

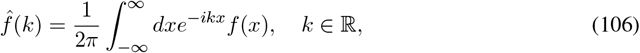

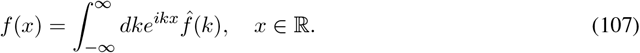

The forward transform maps a function of continuous variable *x* to a function of continuous frequency *k*, and the inverse transform reverses this mapping. The inversion property follows from 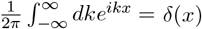.

Multiplying the Dirac comb identity (Eq. 104) by a function *f*(*x*) and integrating over *x* ∈ ℝ yields the Poisson summation formula:

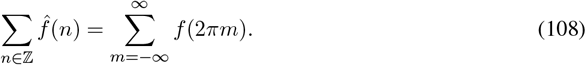

Furthermore, shifting the function before summing, *f*(*x*) → *f*(*x* + *θ*), gives

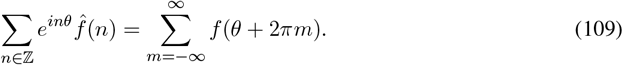

This relationship says that constructing a circular function from Fourier coefficients obtained by sampling the Fourier transform 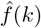 at integer frequencies *k* = *n* yields a 2*π*-periodic “wrapped” version of *f*(*x*). Conversely, given a wrapped function 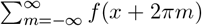, its Fourier coefficients are obtained by sampling the Fourier transform 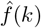 at integer frequencies *k* = *n*.

#### A6.2 Fourier coefficients correlation functions

Consider first the wrapped Gaussian correlation function from the main text (Eq. 9, reproduced here):

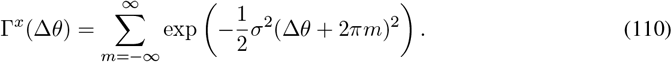

We identify 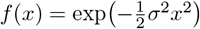, which has Fourier transform

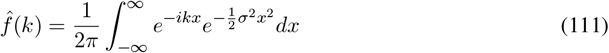

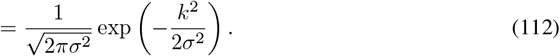

Therefore, the Fourier coefficients of Γ^*x*^(Δ*θ*) are

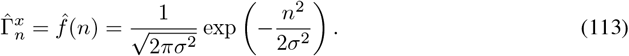

There is a 2-fold degeneracy of the Fourier coefficients 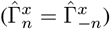 for all *n* except *n* = 0, for which there is no degeneracy. In the more abstract language of Sec. A1, these degeneracies arise from the irreducible representations of *SO*(2), with the trivial representation for *n* = 0 and two-dimensional representations for each doublet.

The two-dimensional analogue of the above analysis applies to the grid-cell case. The original correlation function is (Eq. 23, reproduced here in the *K* = 1 case for simplicity):

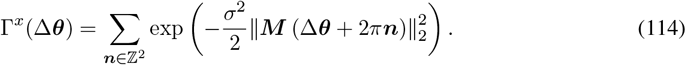

This function has a 12-fold symmetry Γ^*x*^(Δ***θ***) = Γ^*x*^(***g*** Δ***θ***) where ***g*** ∈ *D*_6_, the dihedral group of order 12. Using the Poisson-summation relation, the Fourier coefficients are

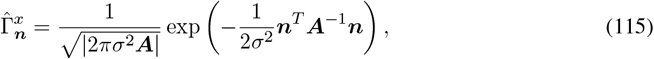

where 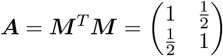 and 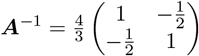. This can be further expressed as

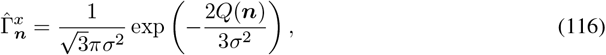

where

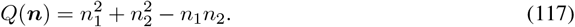

Thus, all integer pairs ***n*** ∈ ℤ^2^ with the same value of *Q*(***n***) yield identical Fourier coefficients. The degeneracy of a given eigenvalue is therefore the number of integer solutions to *Q*(***n***) = *Q*, which is a classical counting problem for the hexagonal lattice. Explicitly, one has

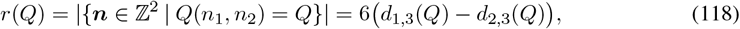

where *d*_1,3_(*Q*) and *d*_2,3_(*Q*) denote the numbers of divisors of *Q* congruent to 1 and 2, respectively, modulo 3. Every degeneracy *r*(*Q*) is a multiple of 6. For several of the first few representable values of *Q* (including *Q* = 1), the degeneracy is exactly 6. In the more abstract language of Sec. A1, these degeneracies arise from the irreducible representations of 𝕋^2^ with hexagonal (*D*_6_) symmetry.

## Appendix B: Methods

### B1 List of main-text symbols

- Network dynamics
  - *x*_*i*_(*t*) – Input current to neuron *i* at time *t*
  - *ϕ*_*i*_(*t*) – Firing rate of neuron *i* at time *t*
  - *ϕ*(*x*) – Single-neuron activation function (softplus or error function)
  - *τ* = 50 ms – Network time constant
  - *N* – Number of neurons in a given network model
- Experimental data and target manifolds
  - *N*_data_ = 1533 – Number of neurons classified as head-direction cells available in dataset [45]
  - *θ* – Head direction angle (parameterizes manifold), circular variable
  - 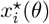 – Target input current for neuron *i* at angle *θ*
  - 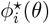 – Target firing rate for neuron *i* at angle *θ*
  - *N*_*θ*_ – Number of angular bins for discretization (typically 100 for data processing, 500–10,000 for simulations)
  - ***X*** – Matrix of target currents, *N*_*θ*_ *× N*
  - **Φ** – Matrix of target firing rates, *N*_*θ*_ × *N*
  - Comparison between generative model and data
    * *I*_HD_ – Head-direction information content [54]
    * *ρ*^flip^ – Flip symmetry correlation
    * *z*_thresh_ – Peak detection threshold for counting peaks in tuning curves
- Network construction and connectivity
  - *J*_*ij*_ – Synaptic weight matrix element from neuron *j* to neuron *i*
  - ***J*** – Full synaptic weight matrix, *N × N*
  - *b* – Bias current for each neuron
  - *c* – Uniform inhibition strength for mean-mode stabilization
  - *λ* – Regularization parameter in optimization; *λ* = 10^−6^ unless otherwise noted
  - ℒ (***J***) – Loss function for weight optimization
  - *C*^*ϕ*^(*θ, θ*^*′*^) – Empirical two-point correlation function: 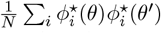
  - *K*_*i,λ*_(*θ, θ*^*′*^) – Kernel matrix for optimization (subtracts out the *i*-th regressor due to constraint *J*_*ii*_ = 0)
- Order parameters and network state
  - *m*(*θ, t*) – Overlap order parameter: 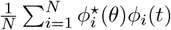
  - 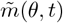 – Temporally low-pass filtered version of 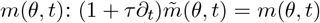
  - *ψ* – Angular center of activity bump, circular variable
  - *ω* (*t*) – Angular velocity input
- Statistical generative process (used in Setting 2 and Setting 3; Sec. B3.1)
  - Γ^*x*^(Δ*θ*) – Two-point correlation function of input current Gaussian process
  - Γ^*ϕ*^(Δ*θ*) – Two-point correlation function of firing rates resulting from input-current Gaussian process samples: 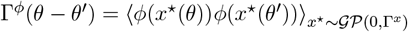
  - **Γ**^*x*^ – Discretized two-point correlation-function matrix for input currents, *N*_*θ*_ × *N*_*θ*_
  - **Γ**^*ϕ*^ – Discretized two-point correlation-function matrix for firing rates, *N*_*θ*_ *× N*_*θ*_
  - *σ* – Fourier decay scale of input-current correlation function (Setting 2 and Setting 3; Sec. B3.1)
  - *β*– Sharpness parameter of activation function (Setting 2 and Setting 3; Sec. B3.1)
  - *b* – Soft threshold in softplus activation function (Setting 2 only; Sec. B3.1)
  - *Z*[*x*^⋆^] – Normalization functional (Setting 2 only)
- Fourier analysis and spectral structure
  - *n* – Fourier mode index; unbounded case *n* ∈ ℤ; with frequency cutoff *n* = −*D*, …, *D*
  - *D* – Frequency cutoff for Fourier modes (results in 2*D* + 1 non-cutoff modes)
  - **Fourier transforms** (using complex exponential basis *e*^*inθ*^):
    * Forward: 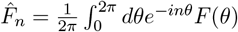
    * Inverse: 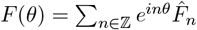
  - 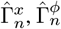 – Fourier coefficients (complex exponential basis) of correlation functions Γ^*x*^(Δ*θ*) and Γ^*ϕ*^(Δ*θ*), computed via 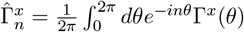; exhibit doublet degeneracy 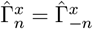
  - 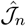 – Fourier coefficients (complex exponential basis) of effective weights 𝒥 (Δ*θ*): given by 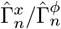
  - *λ*_*n*_ – Eigenvalues of weight matrix ***J*** : given by 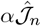
  - **Singular value decomposition quantities** (use sine-cosine modes: constant mode, then (cos(*kθ*), sin(*kθ*)) pairs for *k* = 1, …, *D*, with *n* = 0, …, 2*D* indexing all modes)
    * 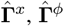 – Vectors of Fourier coefficients for input currents and firing rates, (2*D* + 1)-dimensional, with doublets after first component
    * ***U*** ^*x*^, ***U*** ^*ϕ*^ – Left singular vector matrices of input currents and firing rates, *N*_*θ*_*×* (2*D* + 1); at large *N*, both converge to Fourier mode matrix ***F***
    * ***V*** ^*x*^, ***V*** ^*ϕ*^ – Right singular vector matrices of input currents and firing rates, *N ×* (2*D*+1); rows contain Fourier coefficients of individual neuron tuning curves 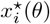 and 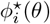
    * ***S***^*x*^, ***S***^*ϕ*^ – Singular value vectors for input currents and firing rates, (2*D*+1)-dimensional; at large *N*, converge to 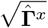 and 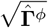
    * ***F*** – Matrix of Fourier modes in sine-cosine basis (so that they are real), *N*_*θ*_ *×* (2*D* + 1)
- Dynamical mean-field theory
  - 𝒥 (Δ*θ*) – Effective rotation-invariant weights in mean-field theory; circulant function with Fourier coefficients given by 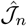
  - *Q*(*t*) – Activity-dependent gain parameter (quadratic functional of 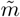)
  - *G*_*Q*_(*h*) – Effective activation function in mean-field equations
  - 𝒰_*ω*_(Δ*θ*) – Angular velocity-dependent kernel for *Q*(*t*) computation
  - *α* – Average slope of activation function: 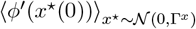
  - *M*(*θ, θ*^*′*^) – Jacobian matrix for stability analysis
- Grid cell extensions
  - ***θ*** = (*θ*_1_, *θ*_2_) – Two-dimensional angular coordinates for grid cells, toroidal variable
  - ***x*** = ***Mθ*** – Two-dimensional Cartesian coordinates for grid cells
  - ***M*** – Transformation matrix for hexagonal geometry: 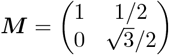
  - ***v***(*t*) = (*v*_*x*_(*t*), *v*_*y*_(*t*)) – Two-dimensional Cartesian velocity
  - ***n*** = (*n*_1_, *n*_2_) – Two-dimensional integer lattice vector
  - *A*(***n***) – Weighting function in grid cell correlation function
  - *η* – Periodicity parameter in grid cell correlation function, 0 ≤ *η* ≤ 1; *η* = 1 gives perfect periodicity
  - *K* – Periodicity parameter in grid cell correlation function, integer ≥1; period of grid pattern is 2*π/K*

### B2 Data processing

We analyzed neural recordings collected by Duszkiewicz et al. [19] (see [45] for the publicly available dataset) from mice exploring a square environment, comprising 1,533 excitatory neurons that were previously identified as head-direction cells. Nine of these mice had viral expression for optogenetic manipulations, though we excluded periods of optogenetic stimulation and processed this data identically to non-optogenetic sessions.

Neural and behavioral data were stored in Neurodata Without Borders (NWB) format. For each mouse, we extracted the animal’s head direction and spike times during periods of square environment exploration. Head direction angles were binned into 100 equally sized bins spanning the full 2*π* radians. For each neuron, we computed raw tuning curves by dividing the spike count in each angular bin by the time spent in that bin.

To reduce noise in the resulting histogram-based tuning curve estimates while preserving non-noise features, we implemented a cross-validated smoothing procedure. For each neuron, we performed the following.

1. Divided the recording session into eight temporal segments of equal duration, labeled alternately as partition 1 or 2 (i.e., 1-2-1-2-1-2-1-2).
2. Computed raw tuning curves independently for each partition.
3. Applied Gaussian smoothing kernels of varying widths to the tuning curve from each partition, with widths logarithmically spaced from 0.1 to 100 bins (corresponding to 2*π/*1000 to 2*π* radians).
4. Evaluated the Poisson log-likelihood of the spiking data from one partition given the smoothed tuning curve from the other partition. Let 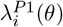 denote the smoothed tuning curve estimate for neuron *i* obtained from partition 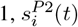 represent the binned spike counts from partition 2, and *θ*(*t*) be the animal’s head direction at time *t*. The Poisson log-likelihood is:

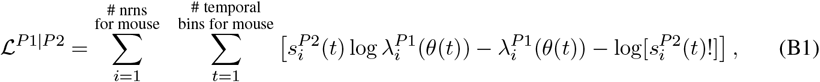

with ℒ^*P* 2|*P* 1^ defined symmetrically.
5. Selected the smoothing width that maximized this cross-validated log-likelihood.
6. Computed final tuning curves by averaging the optimally smoothed curves from both partitions.

Finally, we normalized all tuning curves to have a mean firing rate of unity (arbitrary units) across all head directions. This normalization allowed direct comparison of tuning properties across neurons and animals. These normalized, smoothed tuning curves formed the foundation for subsequent analysis.

### B3 Overview of methods

#### B3.1 Settings 1, 2, and 3

Throughout this paper, we use slightly different choices of activation functions, network dynamics, and weight construction procedures, each suited to a specific modeling or analysis context. For reference, we consolidate these choices here, under the umbrellas of three different settings.

- **Setting 1: Experimental data-derived network model**. For constructing a network model directly from experimental tuning curves (Sec. 2.3), we use the softplus function:

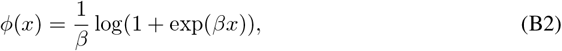

where *β* controls the sharpness of the activation function. We set *β* = 2 in all data-derived analyses. This activation function contains no bias term; instead, we incorporate bias current *b* in the network dynamics (Eq. 1, reproduced here):

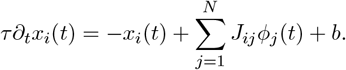

The weights are constructed via least-squares optimization (Sec. A2) to satisfy the vanishing-flow condition on the target manifold. The firing-rate target manifold is defined by the tuning curves themselves, while the input-current target manifold results from applying *ϕ*^−1^(·) to the firing-rate manifold. Thus, using an invertible activation function (i.e., *β <* ∞) is important. After fitting the weights with *b* = 0, we introduce a parameter *c >* 0 to stabilize the mean activity mode; in particular, we shift *J*_*ij*_ → *J*_*ij*_ − *N* ^−1^*c* and set *b* = *c* (Sec. B4.2). These dynamics may receive an additional input corresponding to the modulation of *J*_*ij*_ by angular velocity *ω*(*t*).
- **Setting 2: Experimental data-fitted statistical generative process**. For the statistical model of experimental tuning curves (Sec. 3.2), we use a normalized softplus function (Eq. 10, reproduced here):

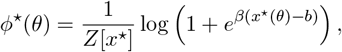

where *β* controls the sharpness, *b* sets a soft threshold for firing, and

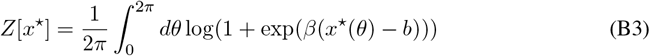

is a normalization functional ensuring unit mean firing rate over *θ*. We use fitted parameters (*σ, β, b*) ≈ (1.42, 2.76, 1.73), where *σ* is the Fourier decay scale of the generative-process correlation function and does not appear in the activation function itself. This activation function is used for statistical modeling and is not embedded within a dynamical model. Correspondingly, no weight matrix is constructed in this context.
- **Setting 3: Analytically tractable statistical generative process and network model**. For large-*N* theoretical analysis of the weights and dynamics (Sec. 4, Sec. 5), we use the error-function activation:

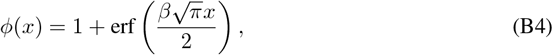

where *β* controls the sharpness. We typically use (*σ, β*) = (1.42, 2.76), where *σ* is the Fourier decay scale of the generative-process correlation function and does not appear in the activation function. We occasionally vary *σ* and *β* to examine parameter dependencies. This activation function contains no bias term. The corresponding network dynamics are (Eq. 1 with *b* = 0, reproduced here):

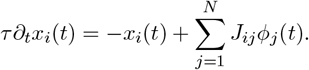

The weight matrix *J*_*ij*_ requires no uniform inhibition since the error function saturates. The weights are constructed via the same least-squares optimization procedure as in the data-derived case, but applied to tuning curves sampled from the generative process.

In Setting 1 and Setting 2, we use two mathematically equivalent forms of bias: an external term *b* in the network dynamics (Setting 1) and a threshold within the activation function (Setting 2). A change of variables *x*_*i*_ → *x*_*i*_ + *b* transforms the former into the latter.

#### B3.2 Non-saturating and saturating activation functions

For the data-derived network model (Setting 1) and generative process (Setting 2), we use unbounded activation functions for the following reasons.

1. This choice is more neurobiologically realistic: cortical neurons typically operate below saturation in their dynamic range [75].
2. The experimental tuning curves show no obvious signs of saturation at high activation levels, making estimation of a saturation point ill-posed.

For theoretical analysis (Setting 3), we switch to bounded nonlinearities for the following reasons discussed in the introduction to Sec. 4.

1. This enables closed-form evaluation of certain integrals in the mean-field derivations, e.g., the function 𝒢 (*C*_11_, *C*_22_, *C*_12_) (Sec. A4.1, Eq. 66).
2. Dynamically, saturation precludes runaway activity, removing the need for the uniform inhibition parameter *c* used in Sec. 2.3.
3. Since we drop the soft threshold *b*, firing rates are non-sparse over *θ* and the normalization functional *Z*[*x*^⋆^] is not needed to amplify small tuning features.

Within each class (non-saturating or saturating), the specific functional form is not critical: different non-saturating nonlinearities (e.g., ReLU versus softplus) or different saturating nonlinearities (e.g., tanh versus error function) would not meaningfully alter network behavior or theoretical conclusions.

#### B3.3 Simulation details

All simulations used network time constant *τ* = 50 ms and Euler integration with timestep Δ*t*.

We simulated recurrent neural networks (Setting 1 and Setting 3) by integrating the rate-based dynamics:

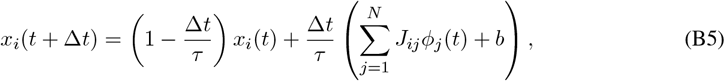

For data-derived networks (Setting 1), we used Δ*t* = 1 ms. For networks constructed from synthetic tuning curves (Setting 3), we used Δ*t* = 5 ms unless otherwise noted. Network construction used regularization parameter *λ* = 10^−6^ unless otherwise noted.

We simulated the evolution of the overlap order parameter 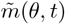 by integrating the dynamical mean-field equations using Euler integration:

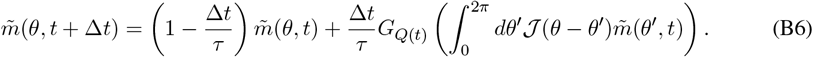

For main text analyses, we used Δ*t* = 0.001. For supplementary analyses, we used Δ*t* = 5 ms.

### B4 Details of data-derived network model

#### B4.1 Double normalization

We performed an additional normalization step for the data-derived network construction. As demonstrated in Sec. 3.1, the data are consistent with distributional circular symmetry, and since *N*_data_ is large, we expect 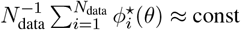. for all *θ*. Furthermore, when pooling tuning curves across mice, as we do here, some degree of circular uniformity is probable due to the possibility of mouse-specific arbitrary phase shifts.

**Figure B1:**
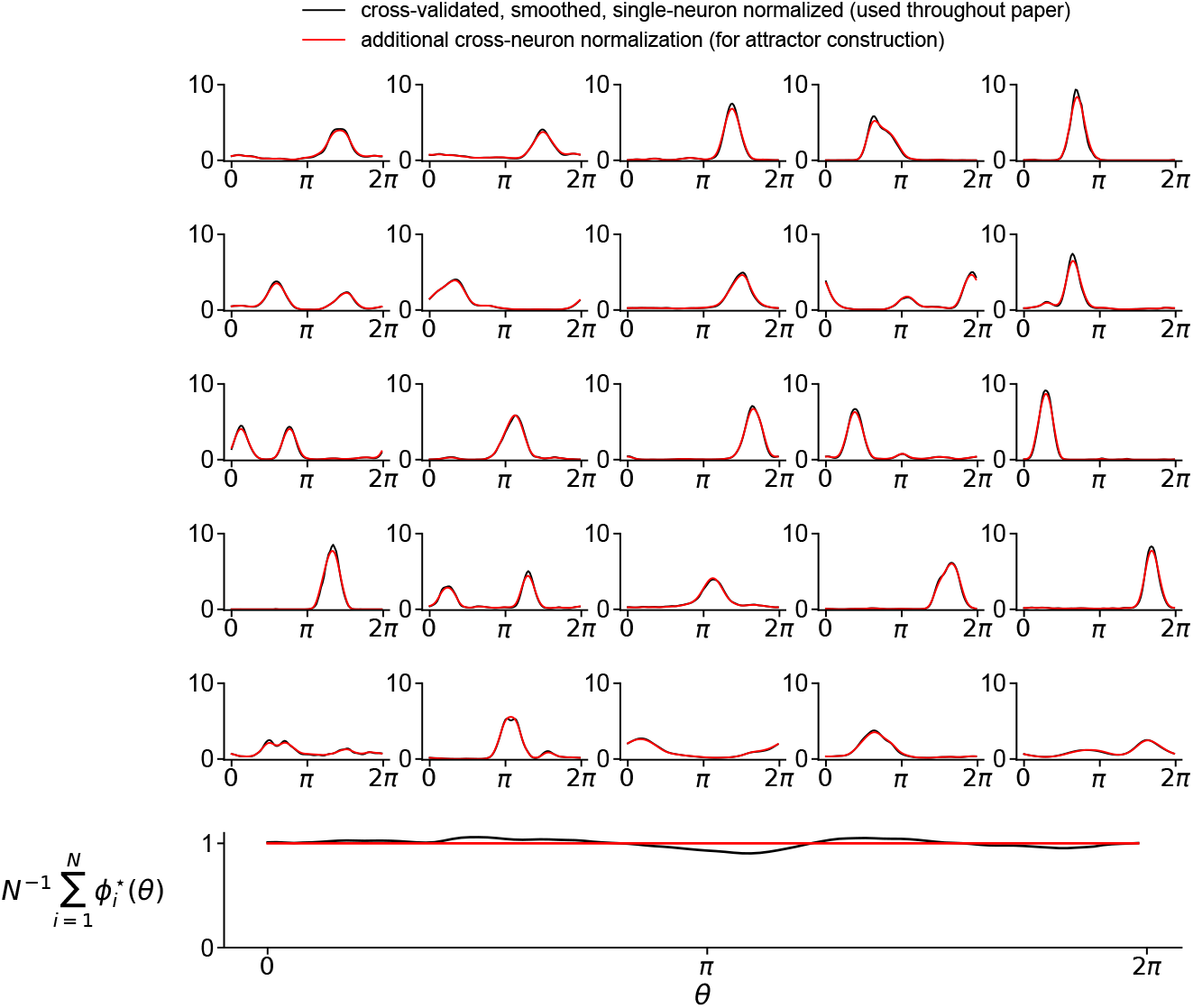
Effect of double normalization on tuning curves. **Top:** Sample of 25 randomly selected tuning curves from the full dataset (*N*_data_ = 1533). Black lines show cross-validated, smoothed tuning curves with single-neuron normalization (used throughout the paper). Red lines show the same curves after additional cross-neuron normalization (used for network construction). The corrections are visually imperceptible at the single-neuron level. **Bottom:** Population-averaged firing rate 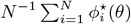 as a function of head direction *θ*. The cross-neuron normalization enforces exact unity at all angles (red line), correcting small deviations from the single-neuron normalized data (black line). The magnitude of these corrections demonstrates that the double normalization introduces only minor adjustments to individual tuning curves.

While the integral over *θ* is exactly unity for each *i* by explicit normalization (Sec. B2), the population average over *i* for each *θ* exhibits small fluctuations around unity. To make this *i* average exactly unity for each *θ*, we simultaneously normalized both the neuronal (*i*) and angular (*θ*) averages to unity using the Sinkhorn-Knopp algorithm, which iteratively normalizes both axes until convergence. This results in a tiny correction to each tuning curve, displayed in Fig. B1.

We make a few further comments about this double normalization. First, in the limit *N* → ∞, achieving the condition 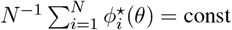. for each *θ* does not require normalization along *θ* for each *i* (although, as we have just described, in our case, this normalization is imposed as well). At large *N*, this condition results (up to 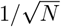 fluctuations) from the distributional circular symmetry of tuning curves, for which we have provided statistical evidence in the mouse data.

Second, even in finite-*N* systems without exact enforcement of the *θ*-independent *i*-average condition (and thus without the reparameterization symmetry under the parameter *c*), one could presumably configure broad inhibition and positive bias currents that stabilize some manifold, albeit not a specific manifold of interest. The fact that it is not exactly a prespecified manifold could be considered problematic from the perspective of our modeling framework, but is not problematic for biological circuits.

**Figure B2:**
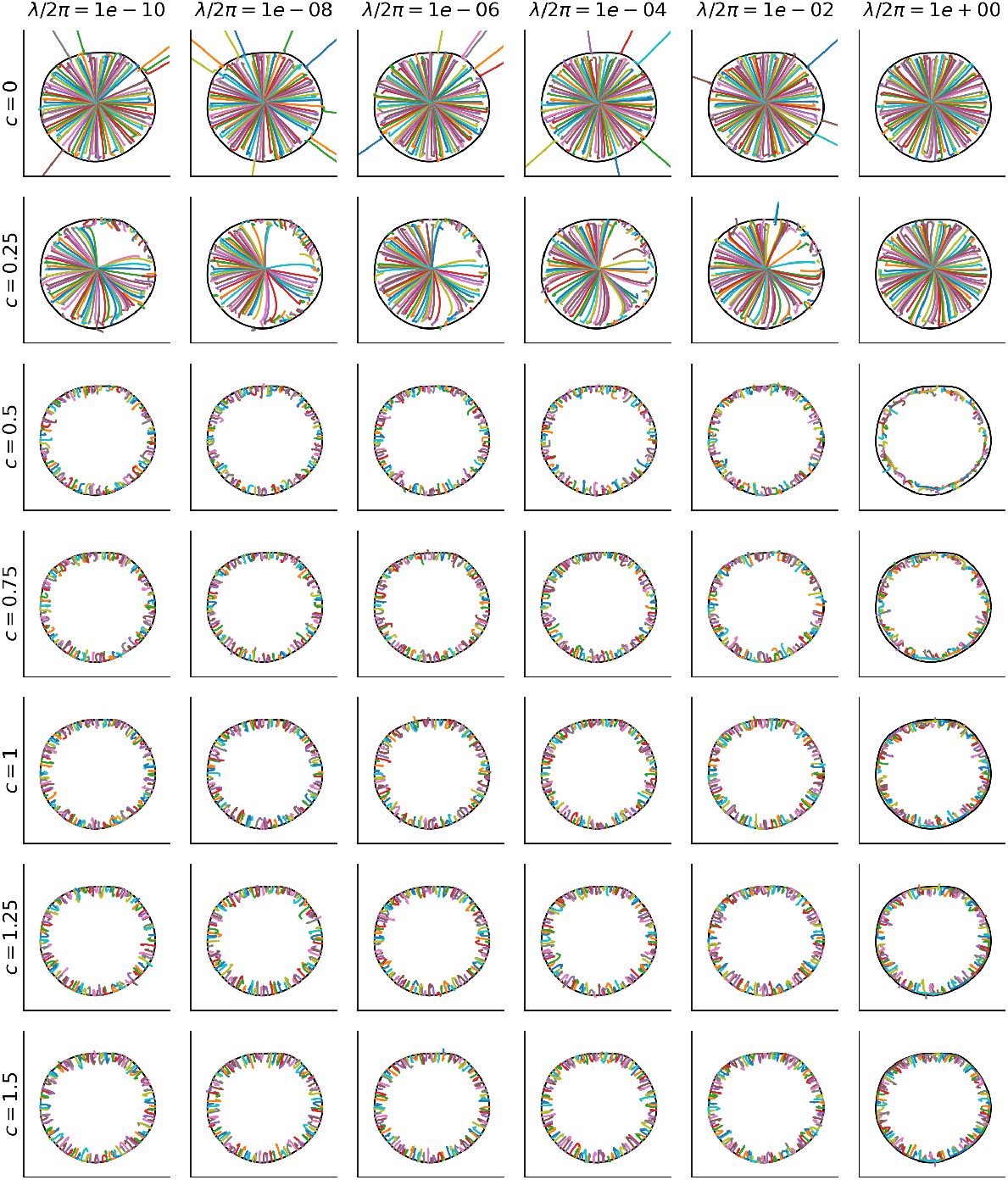
Attractor stability across parameter values (*N* = *N*_data_ = 1533). Each panel shows the plane of the top two principal components of the target manifold (black) with trajectories from 100 initial conditions perturbed from the target manifold by independent Gaussian noise across neurons. Columns correspond to different regularization values *λ*, while rows show different mean-mode stabilization values *c* starting from *c* = 0. Successful attractor behavior requires mean-mode stabilization *c* ≳ 0.5 to ensure convergence to the target manifold rather than collapse to the origin or divergence.

#### B4.2 Stability and c

With *c >* 0, the dynamics become:

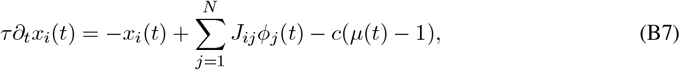

where 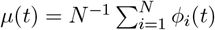 is the population average. Since *µ*(*t*) = 1 when the network lies on the target manifold, this term subtracts input current when *µ*(*t*) *>* 1 and adds input current when *µ*(*t*) *<* 1, maintaining *µ*(*t*) = 1. For *c* ≫ 1, the mean mode exhibits exponential decay to unity with timescale *τ/c*.

#### B4.3 Variants on the weight matrix

For the circulant and noisy circulant variants on the sorted data-derived weight matrix of Sec. 2.3 (Fig. 3C,D), we multiply each matrix by a factor of 1.275 to ensure bump formation. This factor was chosen since it leads to bump formation and produces bumps of similar height to those in networks with the data-derived weight matrix.

#### B4.4 Impact of *λ* and c

We analyzed how attractor stability depends on both *c* (mean-mode stabilization) and *λ* (regularization) in Fig. B2. In the paper, we use *λ* = 10^−6^ and *c* = 1; these figures show the full parameter dependence.

Consider first Fig. B2. The parameter *c* serves two stabilization purposes. When *c* = 0, activity either diverges or collapses to zero depending on initial conditions. For *c >* 0 but small, the collapse-to-origin problem persists even when divergence is prevented. Values *c* ≥ 0.5 stabilize the mean activity mode across all tested initial conditions. Increasing *λ* leads to the stabilization of a different manifold than the target manifold with smaller radius.

Fig. B10 shows that the flow error 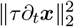 increases with larger *λ* values, as expected, but also increases at small *λ* due to numerical issues. The value *λ* = 10^−6^ used in the paper was chosen to balance between minimizing these sources of error.

#### B4.5 Overlap order parameter provides the MAP estimate of head direction under certain assumptions

Here we consider observed firing rates ***r*** = (*r*_1_, …, *r*_*N*_). This is a random vector, distinct, for example, from the network state vector ***ϕ***(*t*) studied in the paper. Assume that each neuron’s firing rate is conditionally independent given head direction *θ*, with Gaussian noise around the tuning curve,

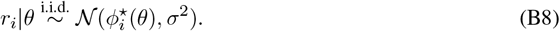

The likelihood is then

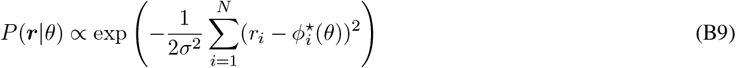

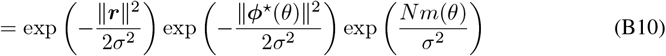

where 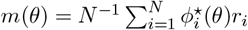 is the overlap order parameter. Under a uniform prior 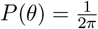 and the large-*N* assumption that 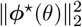 is approximately constant across *θ* due to the circulant statistics of tuning curves (or exactly constant due to explicit double normalization), the posterior probability depends on *θ* only through the exponential term containing *m*(*θ*):

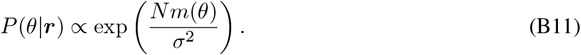

Since *P*(*θ*|***r***) is monotonically increasing in *m*(*θ*), the maximum *a posteriori* estimate is

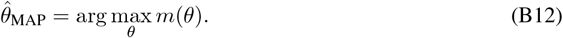

### B5 Structural and global stability

#### B5.1 Structural stability of the manifold

The dynamics of classical ring-attractor models degrade under introduction of structural (i.e., weight) noise. In our model, there is already disorder present in the form of a random embedding, but the question of structural stability remains, namely, what happens when noise is introduced atop this connectivity?

We analyzed structural robustness comparing disordered and circulant networks (Fig. B3). To ensure fair comparison, we constructed both a disordered weight matrix and a circulant counterpart with matching eigenvalues. In particular, the circulant matrix was built using the dynamical mean-field-derived *J* (Δ*θ*) profile discretized over *N* neurons, with additional global inhibition (− 1 subtracted from all weights) to enable unimodal bump formation.

**Figure B3:**
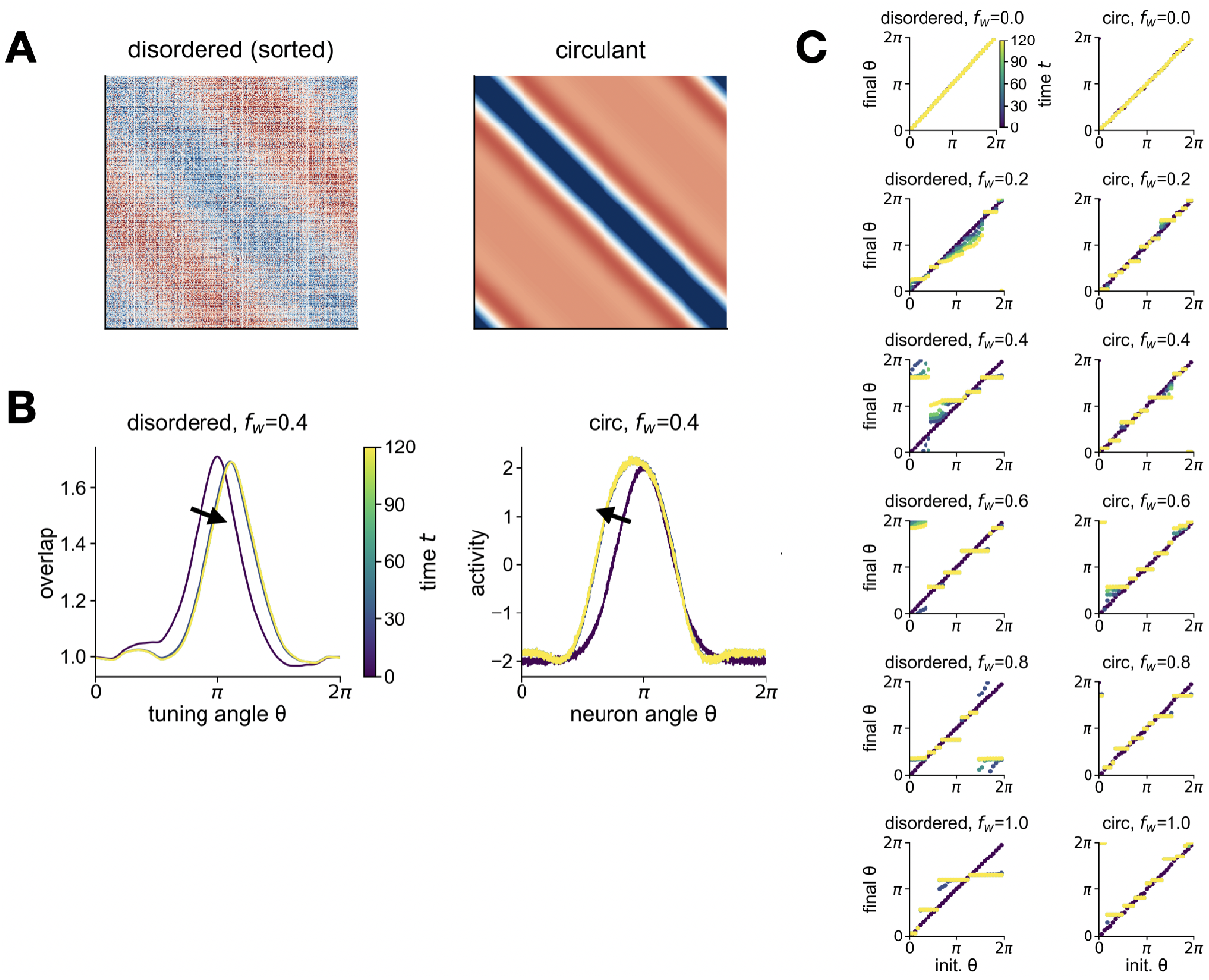
Structural stability comparison between disordered and circulant networks (*N* = 2500, generative-process Fourier decay scale *σ* = 1.42, response sharpness *β* = 2.76). **(A)** Connectivity matrices: disordered network with neurons sorted by center-of-mass angles (left) and corresponding circulant network (right). **(B)** Example trajectories at noise level *α*_noise_ = 0.4 showing angular drift over time for both network types (colors indicate different times), where independent Gaussian noise with standard deviation *α*_noise_ *×* [empirical standard deviation of weights] is added to each weight. **(C)** Angular stability analysis across noise levels. Each panel shows initial angles (x-axis) versus final angles after 30 s evolution (y-axis) for disordered (left column) and circulant (right column) networks. Perfect diagonal alignment indicates stable continuous attractors; deviations reflect noise-induced breakdown. Both network types show comparable degradation patterns as noise strength *α*_noise_ increases from 0 to 1.

We tested both networks under analogous conditions: initialization with slightly noisy patterns distributed uniformly around their respective attractor manifolds, followed by evolution under different levels of additive Gaussian weight noise *α*_noise_ (expressed as a fraction of the standard deviation of the noiseless weights). For each network type, we tracked the angular position of activity patterns over time, defined by the peak of the overlap order parameter for disordered networks and the peak across neurons for circulant networks.

Both network types exhibit comparable degradation of continuous attractor properties under weight noise (Fig. B3C), with clear breakdown of the continuous manifold structure emerging by *α*_noise_ = 0.4. At large noise levels, disordered networks tend to collapse onto a slightly smaller number of discrete attractor states compared to classical models. Conversely, at small noise levels, disordered networks appear to show somewhat more gradual degradation, maintaining approximate continua of fixed points for longer than their circulant counterparts. Overall, both implementations exhibit generally comparable structural stability.

#### B5.2 Global stability of the manifold

To assess the global stability of the target manifold in finite-size networks, we examined whether initial conditions far from the manifold converge to it or to spurious attractors elsewhere in state space. We used the model parameters from 4 (Setting 3; B3.1) with *N* = 2500.

**Figure B4:**
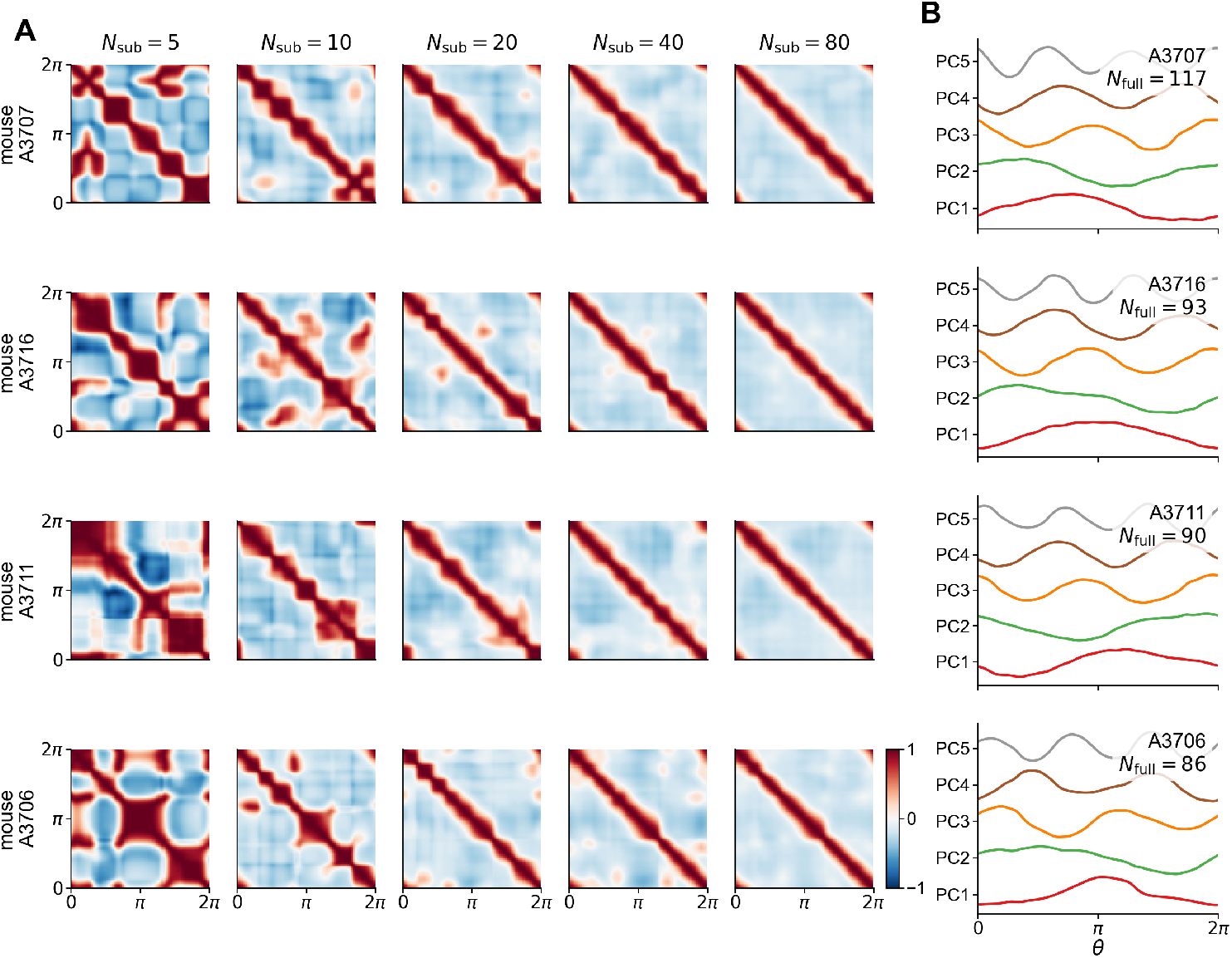
Convergence to circulant forms and Fourier modes in experimental data. **(A)** Convergence to circulant forms of centered correlation matrices (same format as Fig. 4C, which considered uncentered covariance matrices). **(B)** Fourier modes obtained by diagonalizing the full correlation matrices from (A) within each mouse.

We initialized the network at i.i.d. Gaussian states with zero mean and varying standard deviations per neuron, ranging from 1 to 1000. For reference, when the standard deviation per neuron is 𝒪 (1), the projection of initial conditions onto the leading two principal components of the target manifold is 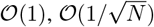 and thus very close to the origin in principal-component space. Initial conditions become comparable in magnitude to the ring in the top-2 principal-component space when the initial-condition standard deviation is on the order of 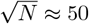.

For each standard deviation, we simulated network dynamics for 30 s and recorded both initial and final states. We then projected these states onto the top two principal components of the target manifold to visualize convergence (Fig. B8).

Across all noise levels tested, initial conditions reliably converged to points on the target manifold, with no evidence of spurious attractors far from the manifold.

**Figure B5:**
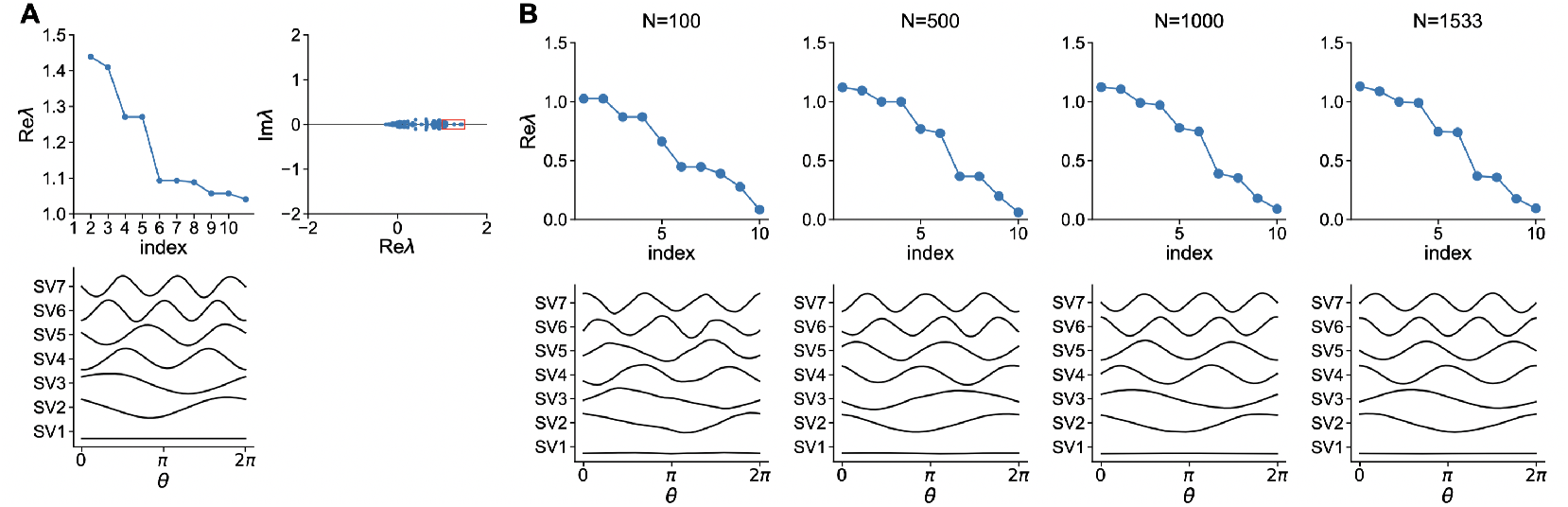
Spectral signatures in data-driven and finite-sample networks. **(A)** Spectrum of the data-derived optimal weight matrix *J*_*ij*_ from Sec. 2.3 and Fig. 3 (*N* = *N*_data_ = 1533). **Top left:** Real parts of eigenvalues ranked by their real parts, showing approximate doublet degeneracy. **Top right:** Complex-plane plot of eigenvalues, demonstrating that they are approximately real despite *J*_*ij*_ not being transpose symmetric. **Bottom:** Left singular vectors of the *N*_*θ*_ *× N*_data_ data matrix **Φ** used to construct the data-derived weight matrix (with double normalization). These spectral signatures derived from the generative process setting (Setting 2; Sec. B3.1) also hold in this data-driven setting (Setting 1; Sec. B3.1). **(B)** Convergence of spectral signatures with increasing sample size. Each column shows eigenvalue spectrum (**top**) and singular vectors (**bottom**) for networks constructed from generative process samples of increasing size (error-function nonlinearity, generative-process Fourier decay scale *σ* = 1.42, and response sharpness *β* = 2.76 as in Sec. 4). Rightmost column shows *N*_data_ = 1533 samples.

**Figure B6:**
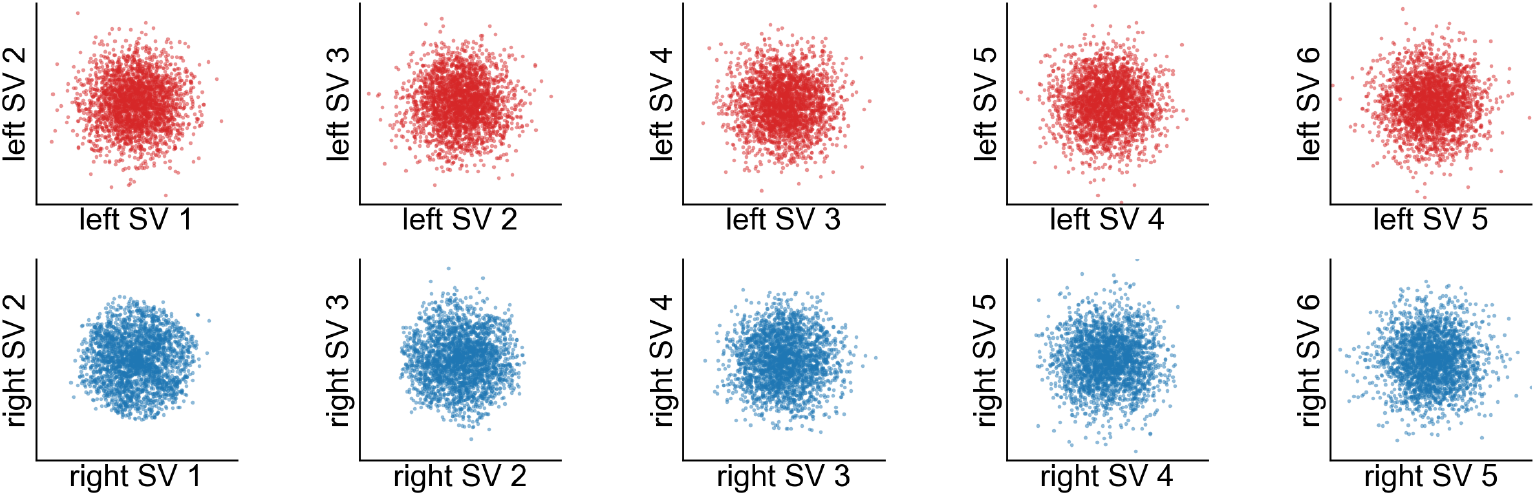
Loadings of left and right singular vectors of the optimal weight matrix (*N* = 2500, generative-process Fourier decay scale *σ* = 1.42, response sharpness *β* = 2.76, error-function nonlinearity). **Top (red):** Loadings of left singular vectors of the optimal weight matrix ***J*** constructed using the error-function generative process. The left singular vectors converge to independent Fourier coefficients of the Gaussian input-current tuning curves. **Bottom (blue):** Loadings of right singular vectors. The right singular vectors converge to non-independent Fourier coefficients of the non-Gaussian firing-rate tuning curves, appearing disordered despite some correlations.

**Figure B7:**
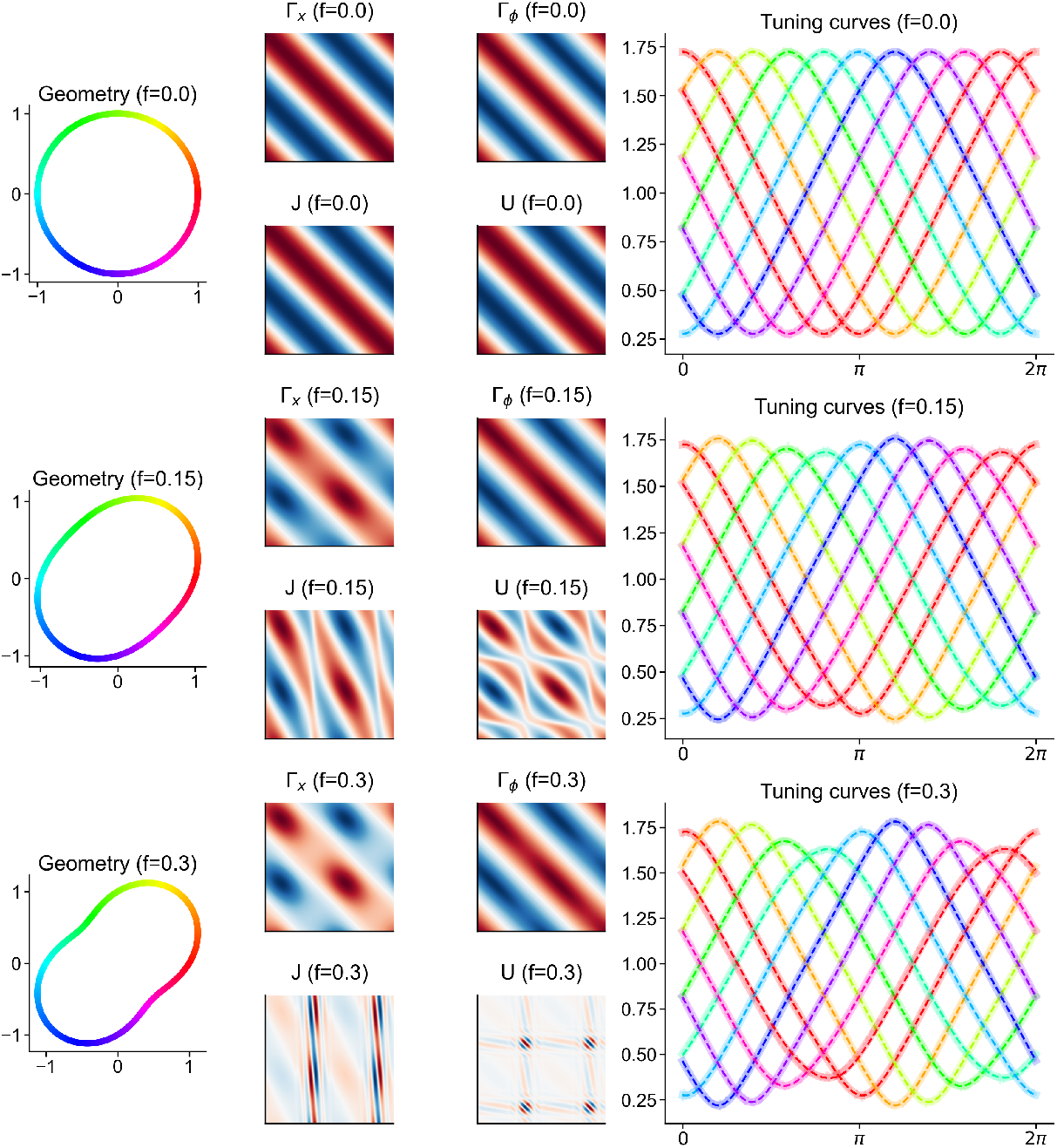
Dynamical mean-field theory of asymmetric continuous attractors (*β* = 2.5, *λ* = 10^−3^, error-function nonlinearity). Panels from top to bottom show increasing values of the asymmetry parameter *f*. For each value of *f* : **Left panel:** Geometric structure of the asymmetric circular manifold. **Middle panels:** Input-current correlation function Γ^*x*^(*θ, θ*^*′*^), firing-rate correlation function Γ^*ϕ*^(*θ, θ*^*′*^), non-circulant interactions 𝒥 (*θ, θ*^*′*^), and gain kernel 𝒰 (*θ, θ*^*′*^). **Right panel:** Angular profiles of the overlap order parameter 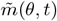 initialized at Γ^*ϕ*^(*θ* − *ψ*) with small Gaussian perturbations (standard deviation = 0.01) for 10 different values of *ψ* (thick solid lines) and the unperturbed states to which they relax after 30 s (thin dotted lines).

**Figure B8:**
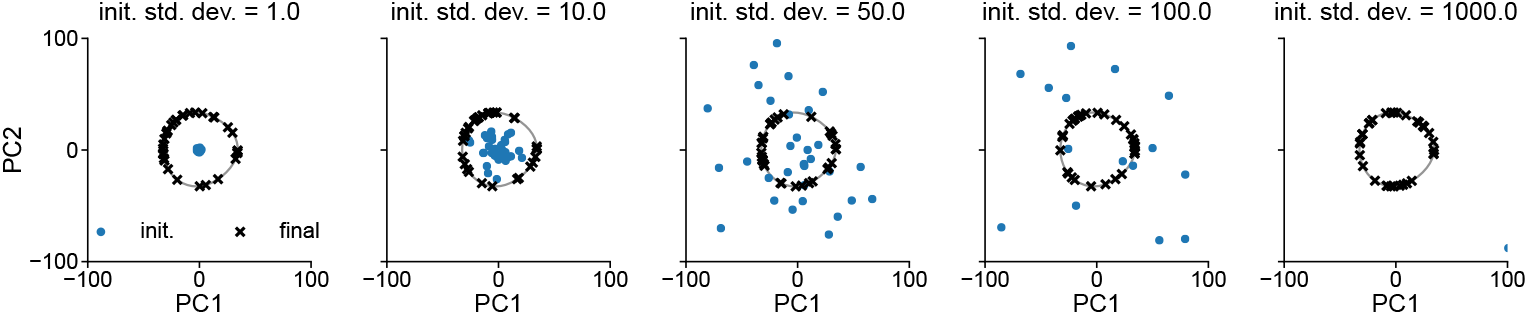
Global stability of the target manifold (*N* = 2500, error-function nonlinearity, generative-process Fourier decay scale *σ* = 1.42, response sharpness *β* = 2.76). Each panel shows initial conditions (blue dots) and final states after 30 s (black crosses) projected onto the top two principal components of the target manifold (gray line). Initial-condition standard deviation per neuron increases from left to right. All trajectories converge to the ring, with no spurious attractors observed.

**Figure B9:**
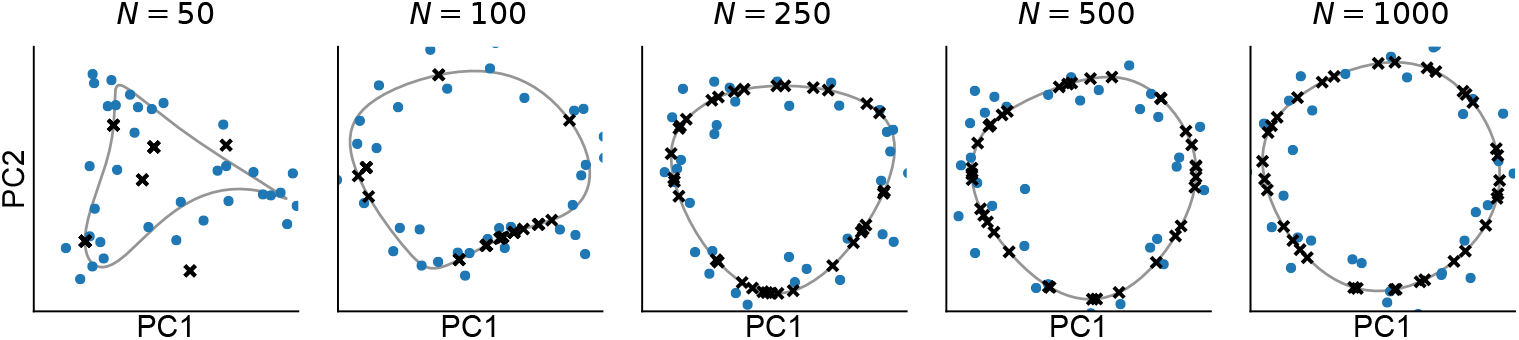
Convergence of dynamics to mean-field behavior with increasing *N* (error-function nonlinearity, Fourier decay scale *σ* = 1.42, response sharpness *β* = 2.76). As in Fig. B8, each panel shows initial conditions (blue dots) and final states after 30 s (black crosses) projected onto the top two principal components of the target manifold (gray line). Initial condition noise is scaled by 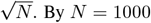, the target manifold resembles a geometric ring and initial conditions reliably converge to it. At small *N*, trajectories sometimes converge to fixed points away from the target manifold. Fig. B8 shows that at large *N* no such spurious off-manifold states are reached, as predicted by the DMFT.

**Figure B10:**
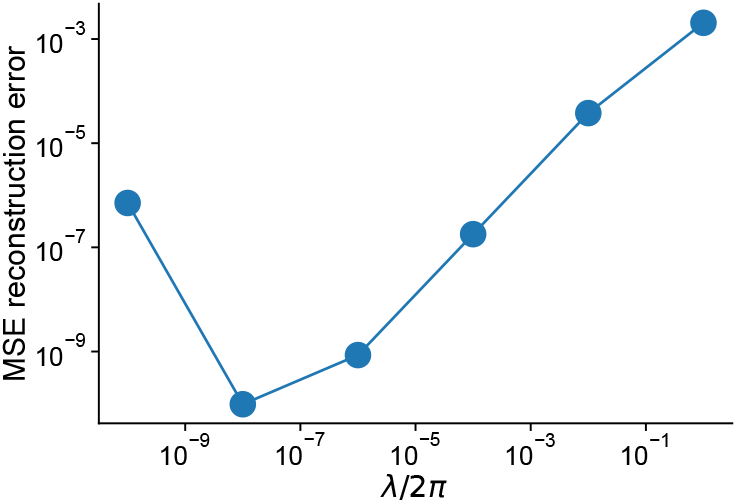
Flow error as a function of regularization parameter for the data-derived network construction (*N* = *N*_data_ = 1533). Flow error is measured as the *L*^2^ norm of *τ∂*_*t*_***x***(*t*) evaluated on the target manifold, as a function of the regularization parameter *λ*.

**Figure B11:**
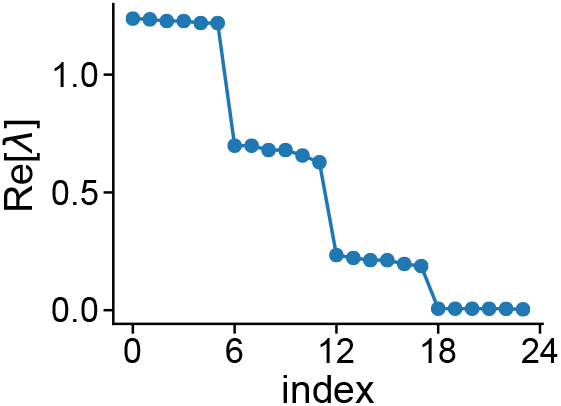
Eigenvalues of the weight matrix ***J*** constructed for the grid-cell model (*N* = 2,500, generative-process Fourier decay scale *σ* = 4, response sharpness *β* = 3, *K* = 1). Eigenvalues versus index showing multiple-of-six degeneracies characteristic of hexagonal symmetry. Same degeneracies are observed for *K >* 1. Network size was reduced from simulations to make diagonalization computationally feasible.

**Figure B12:**
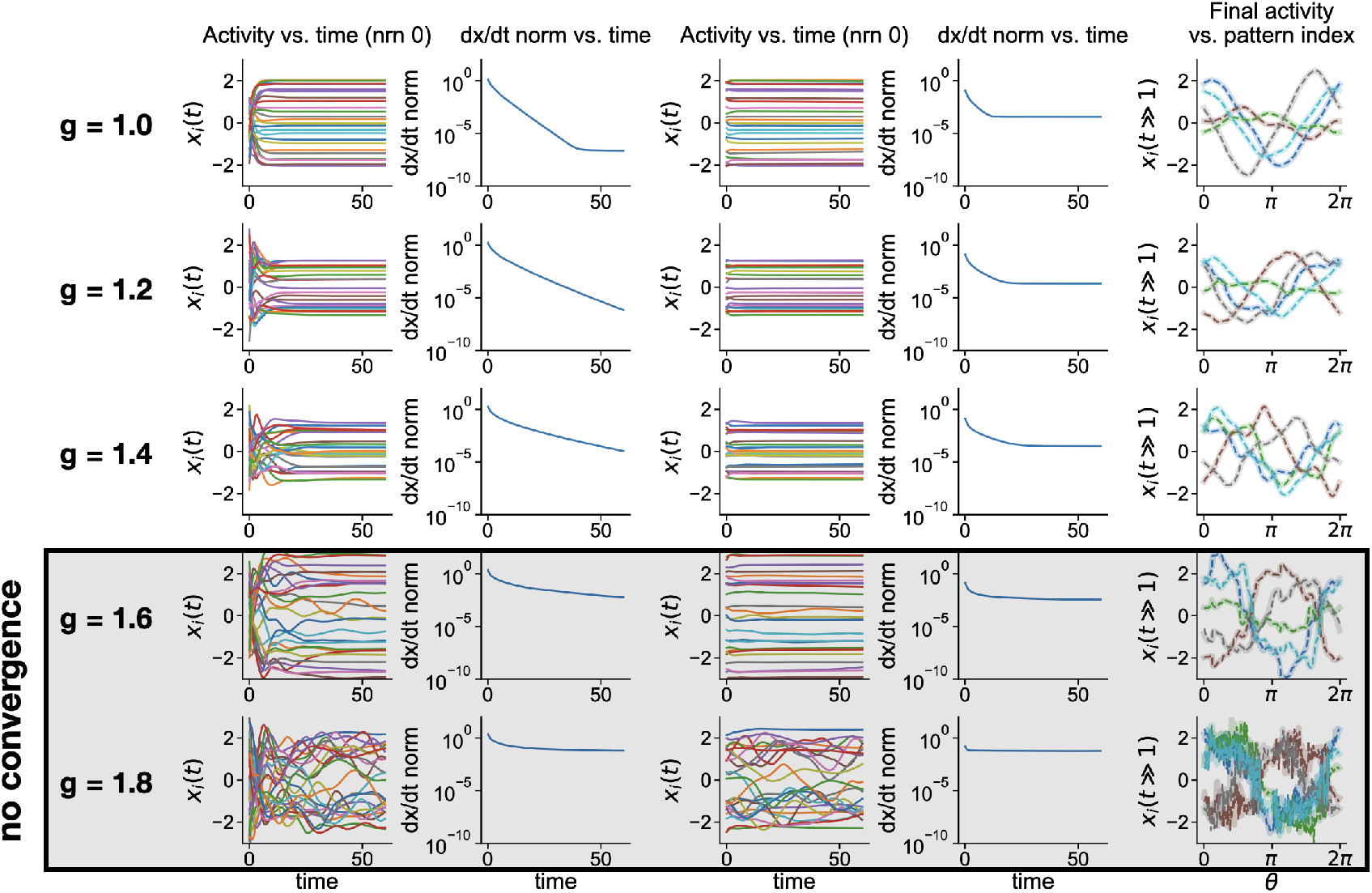
Simulations of the Darshan and Rivkind model [32]. Rows show different values of the random connectivity strength *g*. **First two columns:** Open-loop operation with teacher signal. Left column shows activity (input current) for a single neuron (nrn 0), where each line represents a different teacher signal at different angular coordinates *θ*. Second column shows ∥*∂*_*t*_***x***(*t*) ∥ _2_ versus time. **Third and fourth columns:** Closed-loop (autonomous) operation initialized near open-loop final states. Third column shows activity trajectories for a single neuron as in the leftmost column. Fourth column shows ∥*∂*_*t*_***x***(*t*) ∥ _2_ versus time. **Final column:** Tuning curves defined by the manifold of fixed points, plotting final activity of a given neuron versus coordinate *θ*, insofar as such a stable manifold is formed. Large *g* is necessary for the teacher to dominate chaotic activity, which, in turn, is necessary but not sufficient to form a stable closed-loop manifold [32, 33]. Chaotic activity prevents use as a memory system (left columns). At the same time, large *g* is required to generate heterogeneous tuning curves (right column). The network model constructed in our paper is based on a minimum-norm principle that does not exhibit this tradeoff between stability and heterogeneity.

## Supplement: Dynamics of learned attractors

### S1 Introduction

Here, we revisit reservoir-computing recurrent neural networks that are first trained with open-loop, teacher-forced inputs before operating autonomously in closed-loop mode. We consider the closed-loop dynamics resulting from open-loop learning of fixed points. Rivkind and Barak [33] analyzed local stability around fixed points by computing a response function whose poles correspond to outlier eigenvalues of the Jacobian matrix. Their approach has three limitations. First, it addresses only local linear stability, providing no description of global nonlinear dynamics. Second, the relationship between response-function poles and the full nonlinear dynamics remains unclear. Third, their analysis relies on heuristic Gaussianity assumptions that prevent generalization to structured reservoir architectures.

We revisit this problem through dynamical mean-field theory. Using a path-integral formulation, we derive equations describing the global nonlinear dynamics through order parameters: overlap functions *m*^*µ*^(*t*) that depend on a single time variable and correlation functions *C*(*t, t*^*′*^) that depend on two times. For Gaussian feedback weights, the dynamical mean-field description reduces to an effective recurrent neural network of *P* units, where *P* is the number of prescribed fixed points. Each unit represents an order parameter *m*^*µ*^(*t*) measuring the overlap between the high-dimensional network state and a specific learned pattern. These units interact through an effective weight matrix determined by target-signal statistics and use an activation function that, for *g >* 0, depends on the full history of the system through the two-time correlation function *C*(*t, t*^*′*^).

By linearizing these equations around fixed points, we demonstrate that the order-parameter dynamics are governed by a Jacobian whose eigenvalues precisely match the poles of the response function computed by Rivkind and Barak [33]. This approach eliminates heuristic assumptions, extends naturally to structured reservoirs, provides a complete description of the nonlinear dynamics, and establishes a clear interpretation of response-function poles as eigenvalues governing order-parameter dynamics.

While we focus on the general problem, this formulation can be specialized to continuous manifolds of fixed points, as done by Darshan and Rivkind [32] for ring attractors.

In general, this supplement is self-contained and independent of the main text, though we sometimes refer to it, particularly in Sec. S8 where we provide a taxonomy of ring-attractor models. Variables and parameters are normally represented by the same symbols as those in the main text to which they are analogous, with the exception of what we call here Γ^*µν*^ and Ψ^*µν*^. The former is analogous to Γ^*ϕ*^(*θ, θ*^*′*^) in the main text, and the latter is analogous to Γ^*x*^(*θ, θ*^*′*^). In contrast to the main text, we set the single-neuron time constant *τ* = 1 (equivalently, measuring time in units of *τ*).

### S2 Problem formulation

We consider weight matrices of the form

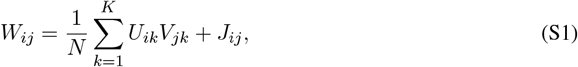

where *N* is the number of neurons, *K* is the rank of the low-rank component, and *P* is the number of training examples. We take the scaling limit *N* → ∞ with *K* and *P* finite.

The weight matrix comprises two components: a learned low-rank structure and random reservoir weights. The low-rank component consists of fixed feedback weights *U*_*ik*_ and learned readout weights *V*_*ik*_. Letting ***u***_*i*_ denote the *i*-th row of the feedback weights *U*_*ik*_, we assume that averages of the form 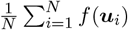 converge to limiting values, denoted ⟨*f*(***u***)⟩_*u*_, as *N* → ∞. The readout weights *V*_*ik*_ are learned using least-squares regression as described below. Finally, the reservoir weights are sampled independently as 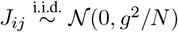, where *g* scales the reservoir connectivity strength. Without the low-rank structure, the network exhibits chaotic activity for *g >* 1 as *N* → ∞ [76].

During training, the system operates in open-loop mode where feedback is clamped to each of *P* teacher signals. Let 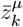 denote the teacher input for example *µ* = 1, …, *P*. For input *µ*, the open-loop network dynamics are

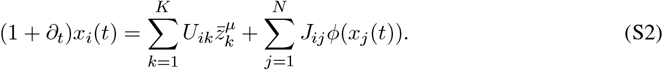

We assume the teacher inputs have sufficient magnitude to suppress chaotic activity if *g >* 1. The network settles to fixed points 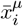 satisfying

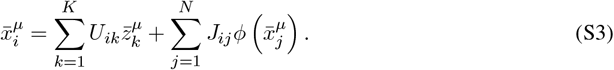

The teacher input correlation matrix is defined as

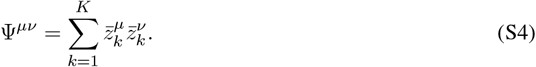

After learning the readout weights (described below), the system operates in closed-loop mode with dynamics

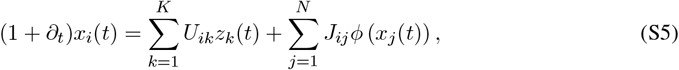

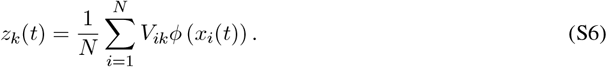

The readout weights *V*_*ik*_ are determined through least-squares regression to ensure the closed-loop system regenerates the open-loop fixed-point activity. The loss function is

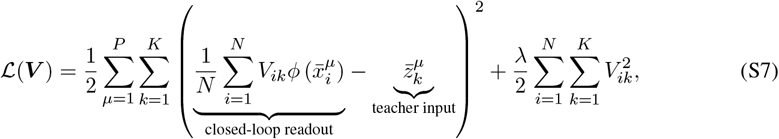

where *λ* ≥ 0 is a regularization parameter. Setting the gradient with respect to *V*_*ik*_ to zero yields the optimal readout weights

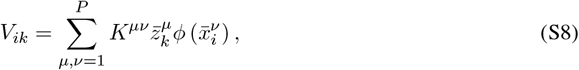

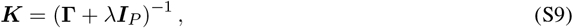

where Γ^*µν*^ is the activity correlation matrix defined below (Eq. S10). In the limit *λ* → 0, the training error vanishes (since there are finitely many training examples but infinitely many neurons), and the open-loop fixed points 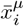 become fixed points of the closed-loop system, though their stability is not guaranteed.

### S3 Order parameters

To analyze the closed-loop dynamics, we define three order parameters:

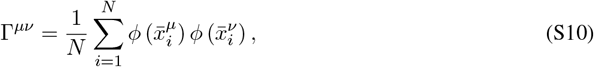

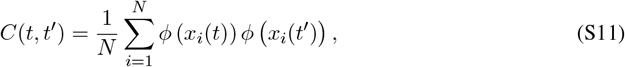

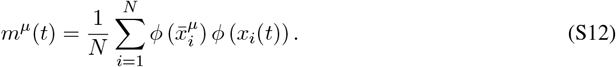

The first is the activity correlation matrix between open-loop fixed points, the second is the two-time correlation function of closed-loop activity, and the third is the overlap function measuring alignment between open-loop fixed points and closed-loop states. Using the parameters Γ^*µν*^ and *m*^*µ*^(*t*), the closed-loop readout can be expressed as

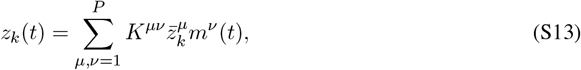

where *K*^*µν*^ is the regularized inverse of Γ^*µν*^ (Eq. S9).

### S4 Single-site dynamics

The dynamical mean-field theory reduces the high-dimensional network dynamics to self-consistent equations for the order parameters. These equations emerge from the saddle-point solution of a path integral (derived in the Appendix) and describe two effective single-neuron or “single-site” processes: one for open-loop fixed points and one for closed-loop dynamics.

These effective single-site processes are given by

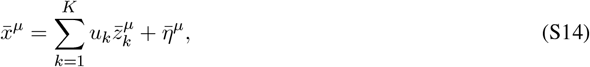

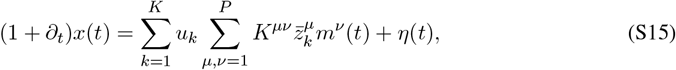

where ***u*** = (*u*_1_, …, *u*_*K*_) represents the feedback weights for a single neuron, i.e., a single row of *U*_*ik*_. The effective noise terms 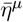 and *η*(*t*) are zero-mean Gaussian random variables with correlations given by

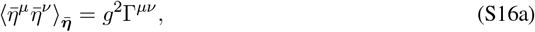

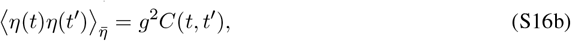

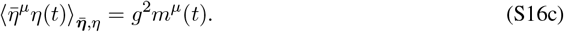

Here we have established the convention that 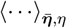 denotes an average over 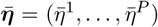 and *η*(*t*). The order parameters obey self-consistency relations

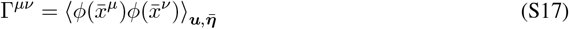

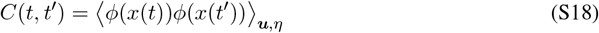

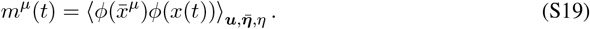

### S5 Differential equations for order parameters

To derive differential equations governing the temporal evolution of the order parameters, we define temporally low-pass filtered versions of the time-dependent order parameters,

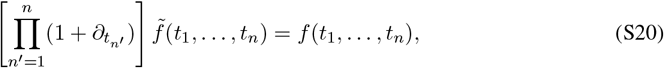

where we will use *n* ∈ {1, 2}. From the self-consistency relations given above, the order parameters satisfy the following differential equations:

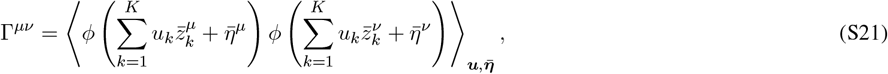

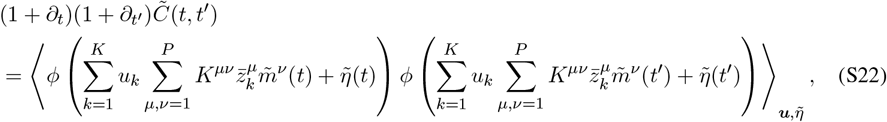

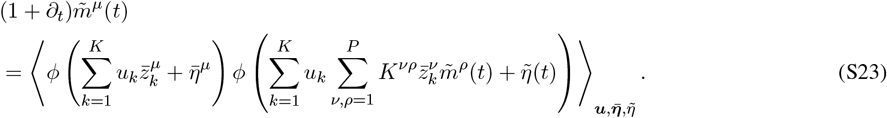

While unwieldy, these provide a closed set of differential equations that govern the evolution of the order parameters. The equation for 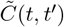 could be numerically integrated on a grid.

### S6 Linearization around fixed points

Having derived the above differential equations, we now analyze the linear dynamics of the order parameters around fixed points. We will set *λ* → 0, in which case fixed points exactly match the teacher signal and ***K*** = **Γ**^−1^. Assuming the closed-loop network state is at fixed point *µ* = **•**, the fixed points of the order parameters are

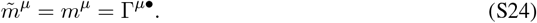

As a sanity check, we compute the closed-loop readout at this fixed point, finding

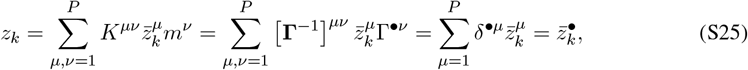

as expected. Finally, the fixed point of the closed-loop two-point function is

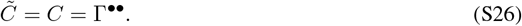

Now, consider perturbations of activity given by

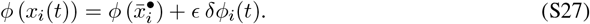

Corresponding perturbations to the order parameters can be written as

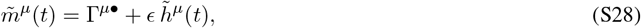

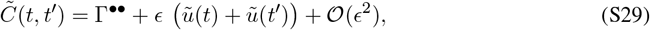

where we have noted that the first-order perturbation to 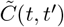 must linearly decompose into terms depending on *t* and *t*^*′*^ that we have denoted 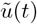 and 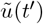. In fact, we have *u*(*t*) = *h*^**•**^(*t*), so the dynamics can also be expressed in terms of *P* variables rather than *P* + 1. However, keeping these variables separate provides additional information about the spectral bulk, as we will show below.

Since the dynamics governing the first-order perturbations are linear, we can remove the tildes, and obtain

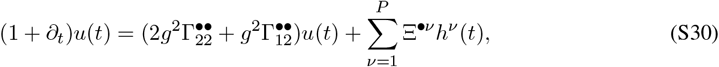

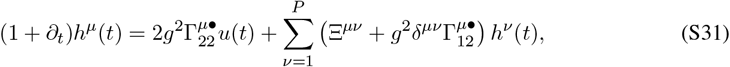

where we have defined

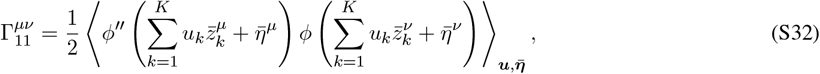

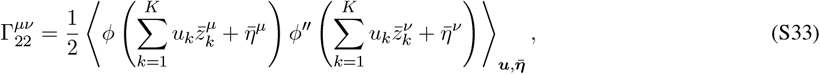

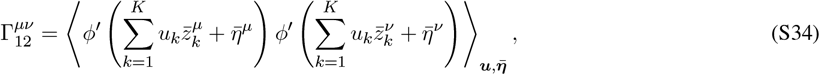

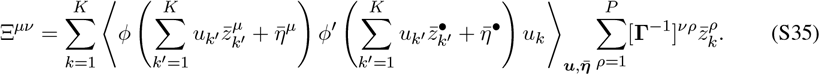

One can identify the (*P* + 1) *×* (*P* + 1) dynamics matrix, or Jacobian, governing these dynamics as

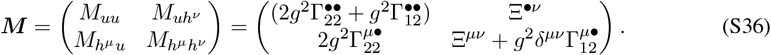

This matrix ***M*** has a left eigenvector (i.e., a right eigenvector of ***M*** ^*T*^) given by

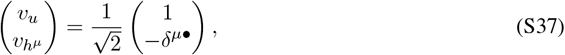

with eigenvalue 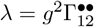. This can be verified by direct calculation. It follows that

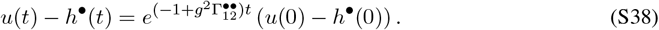

So if we have *u*(*t*) = *h*^**•**^(*t*) at *t* = 0, this holds for all *t*. Furthermore, if we were to non-physically initialize *u*(*t*) and *h*^**•**^(*t*) to be different, this difference decays with rate 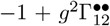, which is the maximum real part of the spectral bulk of the full network’s *N*-dimensional Jacobian at the fixed point. We interpret this to mean that activity outside of the “learned subspace” decays with this timescale.

Assuming *u*(*t*) = *h*^**•**^(*t*) for all *t*, we obtain the *P × P* Jacobian 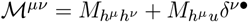, which, upon substituting the formulas for the matrix elements, is

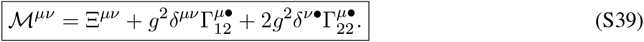

In summary, the linear dynamics of the first-order perturbations to the overlaps, *h*^*µ*^(*t*), are given by

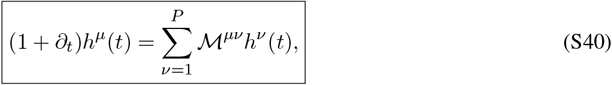

with the *P × P* matrix ℳ^*µν*^ given by Eq. S39.

#### S6.1 Relation to response function

We now explicitly relate the eigenvalues of ℳ ^*µν*^ to the poles of the closed-loop readout response function, which were the output of the calculation of [33]. To match their analysis, we add a source term *w*_*k*_(*t*) to the closed-loop dynamics of the full, *N*-dimensional network,

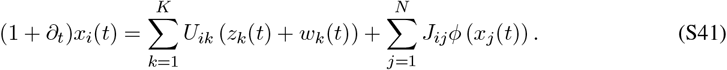

The prior work computed the Fourier space version, denoted *G*_*kℓ*_(*ω*), of the *R × R* response-function matrix

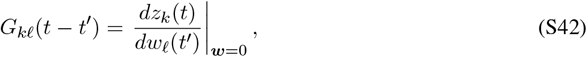

evaluated at a fixed point. When we add this source term, the first-order perturbations to the overlap order parameters get new contributions,

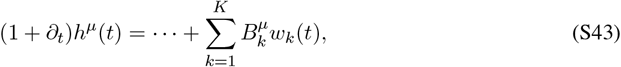

where we have defined a matrix 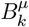 and a matrix 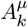 (used below) as

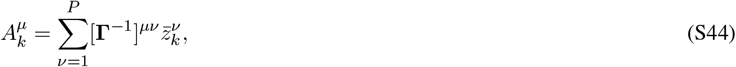

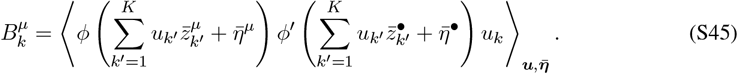

We then have

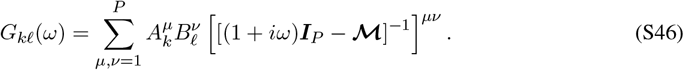

Therefore, the poles of this response function are *ω* = −*i*(−1 + *λ*) where *λ* is an eigenvalue of ℳ^*µν*^. This establishes the connection between the dynamical mean-field theory presented here and the response-function calculation of Rivkind and Barak [33].

### S7 Gaussian feedback weights

#### S7.1 Full dynamical mean-field theory for Gaussian weights

For Gaussian feedback weights, we can derive simplified dynamical mean-field equations. Let 𝒢 (·,·, ·) perform a bivariate Gaussian integral using the activation function *ϕ* (·) (equivalent to Eq. 66, reproduced here):

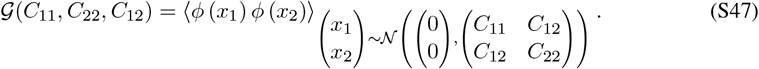

We adopt a simplified notation when applying this to *µ, ν* and *t, t*^*′*^ dependent kernels

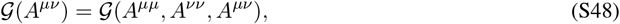

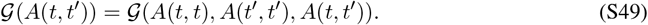

The self-consistent equations are

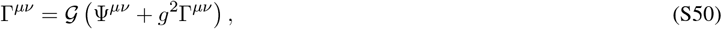

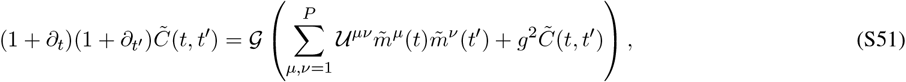

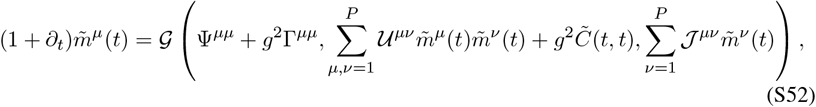

where we have defined *P × P* matrices

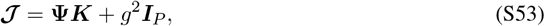

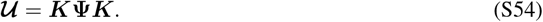

These are analogous to the *N*_*θ*_ *× N*_*θ*_ circulant effective interactions and gain kernel that are central to the dynamical mean-field theory of the model presented in the main text.

#### S7.2 Interpretation as an effective recurrent neural network

The dynamical mean-field equations derived above can be rewritten to highlight their interpretation as an effective recurrent neural network of *P* units. In particular, we have

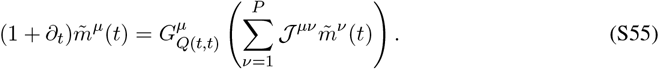

Here, we have defined the effective activation function (analogous to Eq. 69):

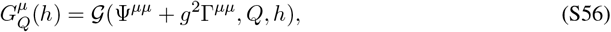

We have also defined *Q*(*t, t*^*′*^) to be the two-time correlation function of the input currents, satisfying (analogous to Eq. 19):

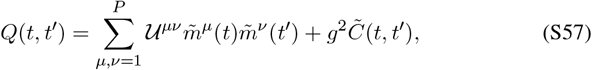

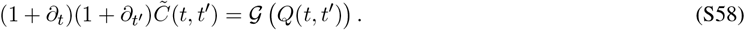

The dynamical mean-field theory of Eq. S55 describes an effective recurrent neural network of *P* units, where each unit corresponds to an order parameter 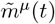 representing the overlap between the high-dimensional network state and a learned pattern. The units interact through an effective weight matrix 𝒥^*µν*^ defined via the teacher correlation structure (Ψ^*µν*^), the regularized inverse firing-rate correlation structure (*K*^*µν*^), and the random connectivity strength (*g*^2^).

The activation function 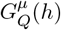 is modulated by the quantity *Q*(*t, t*), which, for *g >* 0, depends on the full history of the system; for *g* = 0, *Q*(*t, t*) depends instantaneously on the system state, making the analogous to a conventional recurrent neural network more direct (though the activation function is modulated by the global system state through *Q*(*t, t*)).

#### S7.3 Linearization around fixed points with Gaussian weights

When applying the Gaussian assumption to the linearized dynamics around fixed points, we can further simplify the Jacobian. By integrating by parts

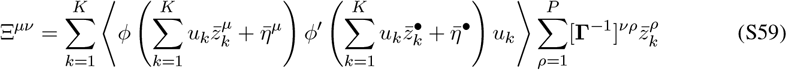

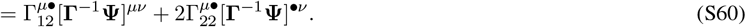

This gives

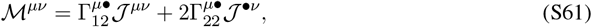

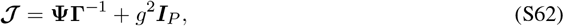

where the expression for **𝒥** is the same as Eq. S53 but with ***K*** = **Γ**^−1^ since *λ* = 0.

### S8 Taxonomy of ring-attractor models through dynamical mean-field theory

We now specialize the general framework a continuous manifold by taking the limit *P*→ ∞ after *N* → ∞. To embed a geometric ring into the network, we define the teacher signal using scaled Fourier modes:

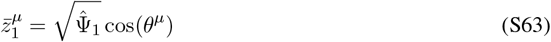

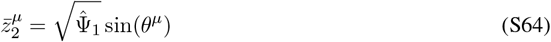

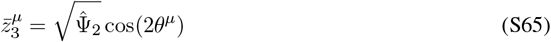

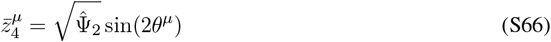

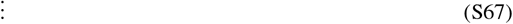

where *θ*^*µ*^ = 2*π*(*µ* − 1)*/P* for *µ* = 1, …, *P*, and 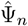 are Fourier coefficients. The rank *K* determines how many Fourier modes are included: *K* = 2 uses only the first harmonic, *K* = 4 uses the first two harmonics, and so on. The resulting teacher correlation matrix has the circulant form

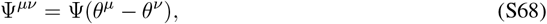

where 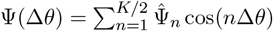 is the inverse Fourier transform of the coefficients 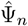.

This continuous-manifold limit yields a taxonomy of ring-attractor models unified by the dynamical mean-field framework. Models are classified by three parameters:

- **Feedback rank** *K*: number of Fourier modes in the teacher signal
- **Random connectivity strength** *g*: controls chaotic fluctuations in reservoir
- **Embedding type**: Gaussian (random *U*_*ik*_) versus Fourier (structured *U*_*ik*_ = *u*_*k*_(*θ*_*i*_))

Combinatorial selection of these parameters generates eight possibilities:

1. ***K* = 2, *g* = 0, Fourier embedding:** Model of Ben-Yishai et al. [2]; classical ring-attractor model with cosine connectivity.
2. ***K >* 2, *g* = 0, Fourier embedding:** Classical ring-attractor model with higher-order Fourier modes in connectivity.
3. ***K* = 2, *g >* 0, Fourier embedding:** Model of Darshan and Rivkind [32].
4. ***K >* 2, *g >* 0, Fourier embedding:** Generalization of Darshan and Rivkind [32]; reduces to case 2 as *g* → 0 (not published).
5. ***K* = 2, *g* = 0, Gaussian embedding:** Model of Beiran et al. [36] and of Mastrogiuseppe and Ostojic [35] with *g* = 0 (up to (non-)Gaussianity, see below); ring attractor with heterogeneous tuning-curve amplitudes.
6. ***K >* 2, *g* = 0, Gaussian embedding:** Model of this paper.
7. ***K* = 2, *g >* 0, Gaussian embedding:** Generalization of Darshan and Rivkind [32]; reduces to case 5 as *g* → 0 (not published).
8. ***K >* 2, *g >* 0, Gaussian embedding:** Generalization of Darshan and Rivkind [32]; reduces to case 6 as *g* → 0 (not published).

Case 5 requires a couple of comments. First, Beiran et al. [36] and Mastrogiuseppe and Ostojic [35] constructed a rank-two ring-attractor model using jointly Gaussian loadings with appropriately configured correlations. In case 5, ***U*** is marginally Gaussian but the full joint distribution over ***U*** and ***V*** is non-Gaussian due to the nonlinearity (Sec. A1). As explained in Sec. A1, this difference is minor for *K* = 2, as in case 5 (since, for both Gaussian and non-Gaussian loadings, the symmetry group of the loading distribution is the circle) but becomes pertinent for *K >* 2 (in which case Gaussian and non-Gaussian loadings lead to different symmetry groups for the loading distribution).

Second, Mastrogiuseppe and Ostojic [35] studied the same ring-attractor structure as in Beiran et al. [36] with added i.i.d. random connectivity scaled by *g >* 0. While this might appear equivalent to case 7 in our taxonomy (which reduces to case 5 as *g* → 0), they differ in the following way. In Mastrogiuseppe and Ostojic [35], the low-rank and i.i.d. components are independent, analogous to the circulant-plus-noise constructions examined in Sec. 2.3 and Sec. 4.2, though using a Gaussian rather than Fourier embedding. In contrast, the framework upon which the taxonomy is based involves learning in an open-loop setting where the low-rank component is correlated with *J*_*ij*_.

#### S8.1 Comparison to Darshan and Rivkind model

Recall that our model achieves tuning heterogeneity through multiple Fourier modes (*K >* 2), without random background weights (*g* = 0), and a Gaussian embedding; while the Darshan and Rivkind [32] model achieves heterogeneity through two modes (*K* = 2), random background weights (*g >* 0), and a Fourier embedding. The conceptual and functional consequences that result from these differences include the following.

- **Interpretability of effective dynamics**. Because the Darshan and Rivkind [32] model uses a Fourier embedding, its dynamical mean-field theory does not reduce to an effective recurrent neural network, obscuring its relationship to classical continuous-attractor dynamics. Furthermore, even with a Gaussian embedding, *g >* 0 would prevent such interpretation due to history dependence in the effective activation function. While this limitation limits theoretical understanding, it is not a functional problem.
- **Stability under increasing heterogeneity**. The Darshan and Rivkind [32] model is non-minimum-norm and operates near the chaotic regime as its mechanism for generating tuning heterogeneity. That introduces the functional problem that, once tuning curves deviate substantially from cosine-like profiles, chaotic fluctuations destabilize the manifold, preventing its use as a memory system (Fig. B12)—though unlike circulant-plus-noise constructions (Sec. 2.3), collapse to a few discrete fixed points is avoided in the non-chaotic regime. Given tuning curves produced by the Darshan and Rivkind [32] model, one could apply the minimum-norm prescription to obtain a version of our model with those tuning-curve statistics, thereby eliminating chaotic instability at large heterogeneity. This raises the question of why a neural circuit would implement the Darshan and Rivkind [32] weight matrix. One possibility is that operating near chaotic instability proves computationally useful in circuits that must combine memory of a continuous variable with dynamic computations.
- **Flexibility for matching experimental data**. The Darshan and Rivkind [32] model produces only a one-dimensional family of tuning-curve statistics parameterized by *g*. In contrast, our minimum-norm prescription allows specification of arbitrary correlation functions for input currents, providing greater flexibility for matching experimental observations. While not infinitely flexible, this approach provides sufficient degrees of freedom to achieve quantitatively accurate fits to experimental head-direction tuning curves with only three fitted parameters.

### S9 Path-integral derivation

We write down a path integral, within the Martin–Siggia–Rose–Janssen–De Dominicis framework, that describes both the open- and closed-loop activity [77]. This requires the introduction of a set of conjugate fields, denoted with hats, that enforce the fixed-point (open-loop) and dynamical (closed-loop) network equations through integral representations of delta functions.

Using from the start the least-squares solution for *V*_*ik*_, the path integral depends on *J*_*ij*_ and *U*_*ik*_ and is given by

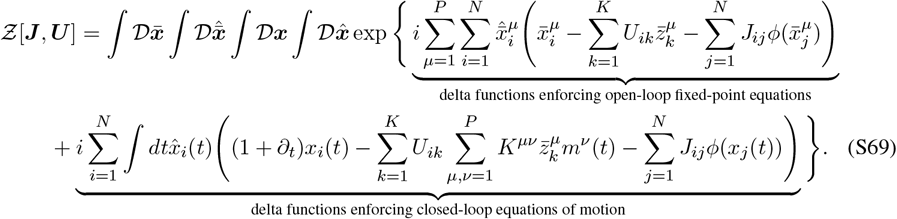

Here 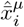 and 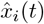 are the conjugate fields that enforce the open-loop and closed-loop dynamics, respectively. We first average over the random reservoir weights, which multiply both open-loop and closed-loop terms in the action. Averaging over Gaussian *J*_*ij*_ generates cross-terms between these contributions, yielding three types of terms (open-open, closed-closed, and open-closed) that correspond directly to the three order parameters. In particular, the open-closed overlap *m*^*µ*^(*t*) emerges automatically from this procedure:

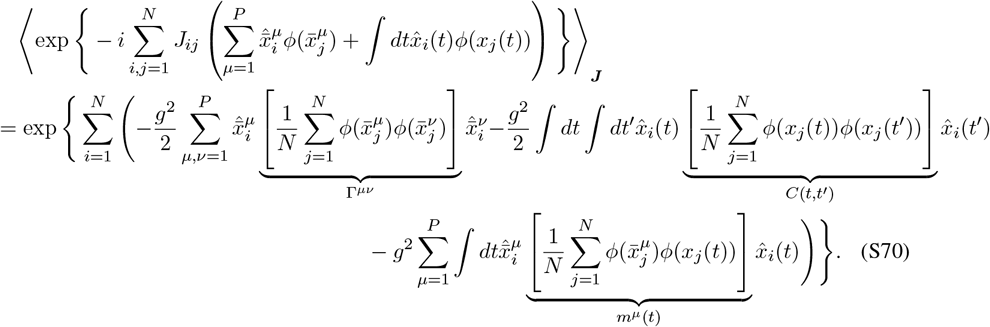

To express the path integral as a sum over the neuron index *i*, we introduce the order parameters indicated in Eq. S70 and enforce their definitions through further integral representations of delta functions. This involves introducing additional conjugate fields 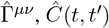, and 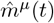:

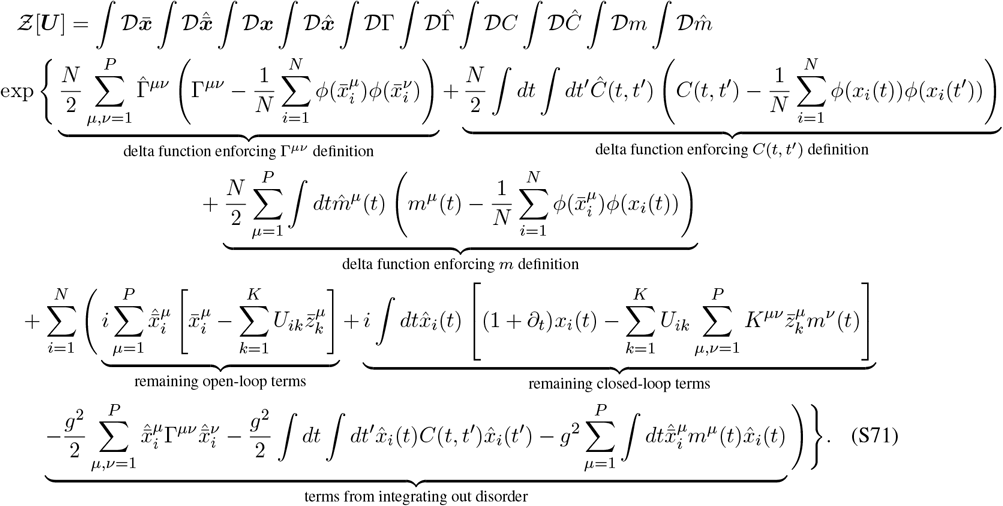

This can be expressed more compactly as

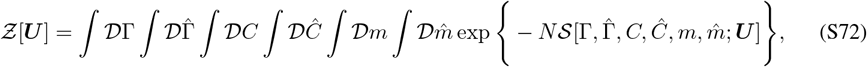

where the order-one action is defined as

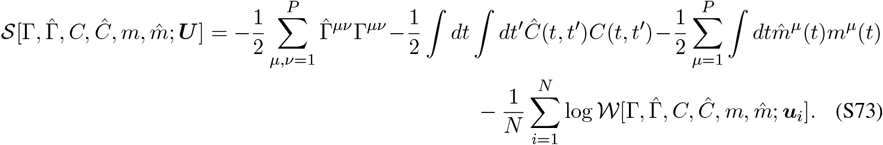

The single-site path integral 𝒲 […; ***u***], which depends on a single row ***u*** of the feedback weight matrix, is given by

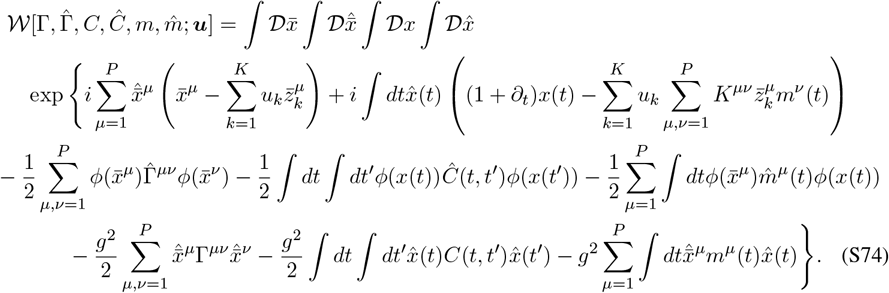

The final three terms can be reformulated using jointly Gaussian random fields. We introduce a Gaussian field *η*(*t*) and a Gaussian vector 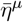 with zero means and covariances given by (identical to Eq. S16):

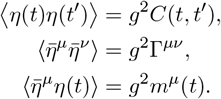

This allows us to rewrite the single-site path integral as

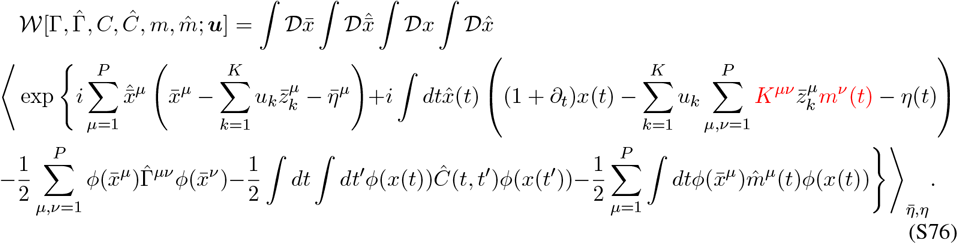

The derivation is exact to this point. We now take the *N* → ∞ limit, where the path integral is dominated by the saddle point at which derivatives of the action vanish. In anticipation of this large *N* limit, we make the replacement in the action:

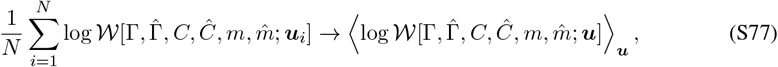

which removes the lingering *i* dependence. The saddle-point equations are obtained by setting derivatives of the action to zero:

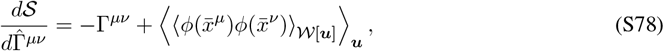

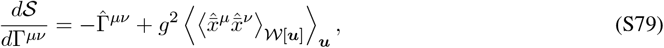

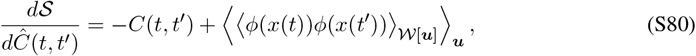

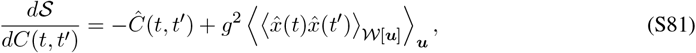

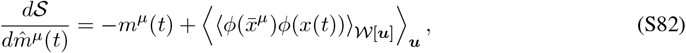

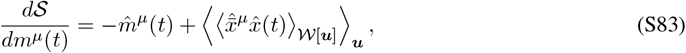

where ⟨*· · ·*⟩_𝒲[***u***]_ denotes an average over the single-site open- and closed-loop processes described by 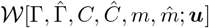. When taking these derivatives, we exclude contributions from the highlighted order parameters *K*^*µν*^ and *m*^*ν*^ (*t*) in Eq. S76, since these quantities appear directly in the original high-dimensional dynamics rather than being introduced through conjugate field pairs in the path-integral formulation. Using the vanishing of pure-hat expectation values, we obtain the saddle-point equation solutions. We indicate that these are the solutions using a * subscript:

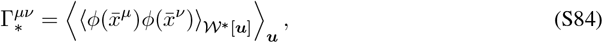

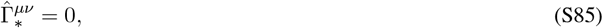

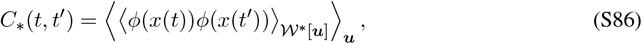

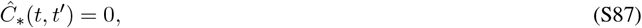

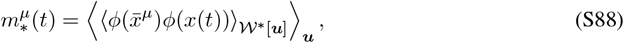

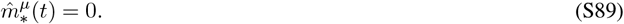

Here 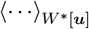 denotes an average within the dynamic process described by 𝒲[Γ_*_, 0, *C*_*_, 0, *m*_*_, 0; ***u***]. Since this paper considers only the saddle-point values of order parameters, we drop the * subscript.

